# Towards an evolutionarily appropriate null model: jointly inferring demography and purifying selection

**DOI:** 10.1101/2019.12.18.881516

**Authors:** Parul Johri, Brian Charlesworth, Jeffrey D. Jensen

**Author notes:** Corresponding authors: Parul Johri, Life Sciences C, Room 340, 427 East Tyler Mall, Arizona State University, Tempe, AZ 85281, Jeffrey D. Jensen, Life Sciences C, Room 340A, 427 East Tyler Mall, Arizona State University, Tempe, AZ 85281.

## Abstract

The question of the relative evolutionary roles of adaptive and non-adaptive processes has been a central debate in population genetics for nearly a century. While advances have been made in the theoretical development of the underlying models, and statistical methods for estimating their parameters from large-scale genomic data, a framework for an appropriate null model remains elusive. A model incorporating evolutionary processes known to be in constant operation - genetic drift (as modulated by the demographic history of the population) and purifying selection – is lacking. Without such a null model, the role of adaptive processes in shaping within- and between-population variation may not be accurately assessed. Here, we investigate how population size changes and the strength of purifying selection affect patterns of variation at neutral sites near functional genomic components. We propose a novel statistical framework for jointly inferring the contribution of the relevant selective and demographic parameters. By means of extensive performance analyses, we quantify the utility of the approach, identify the most important statistics for parameter estimation, and compare the results with existing methods. Finally, we re-analyze genome-wide population-level data from a Zambian population of *Drosophila melanogaster*, and find that it has experienced a much slower rate of population growth than was inferred when the effects of purifying selection were neglected. Our approach represents an appropriate null model, against which the effects of positive selection can be assessed.

## INTRODUCTION

At the founding of population genetics in the early 20th century, R.A. Fisher, J.B.S. Haldane, and S. Wright developed much of the mathematical and conceptual framework underlying the study of population-level processes controlling variation observed within- and between-species. However, as shown by decades of published interactions between them, they held differing views regarding the relative importance of adaptive vs. non-adaptive processes in driving evolution (Provine 2001). As pointed out by J.F. Crow (2008), these issues were not really resolved, but “rather they were abandoned in favor of more tractable studies”. With the advent of the Neutral Theory (Kimura 1968, 1983; King and Jukes 1969; Ohta 1973), the evolutionary importance of stochastic effects due to finite population size, as earlier advocated by Wright, received renewed attention.

In the following decades, further theoretical developments as well as the availability of large-scale sequencing data have validated the important role of genetic drift (Kimura 1983; Walsh and Lynch 2018). However, subsequent research on the indirect effects of selection on patterns of variability at linked neutral sites has re-ignited previous debates (Kern and Hahn 2018; Jensen *et al.* 2019). In particular, it remains unclear whether the large class of deleterious variants envisaged in the Neutral Theory, and their effects on linked neutral sites (background selection, BGS), are sufficient to explain genome wide patterns of variation and evolution, or whether a substantial contribution from the effects of beneficial variants on linked neutral sites (*i.e*., selective sweeps), is also required (see review of Stephan 2010).

The primary difficulty in answering this question stems from our lack of an appropriate neutral null model - that is, a model incorporating genetic drift as modulated by the demographic history of the population, as well as a realistic distribution of fitness effects summarizing the pervasive effects of both direct and indirect purifying selection. Without a model incorporating these evolutionary processes, which are certain to be occurring constantly in natural populations, it is not feasible to quantify the frequency with which adaptive processes may also be acting to shape patterns of polymorphism and divergence.

It can, however, be difficult to distinguish the individual contributions of positive and purifying selection from demographic factors such as changes in population size, as all of these evolutionary processes may affect allele frequency distributions and patterns of linkage disequilibrium in similar ways. For example, both purifying selection and population growth can distort gene genealogies at linked neutral sites in a similar fashion (Charlesworth *et al.* 1993, 1995; Kaiser and Charlesworth 2009; O’Fallon *et al.* 2010; Charlesworth 2013; Nicolaisen and Desai 2013), and result in a skewing of the site frequency spectrum (SFS) towards rare variants. In fact, demographic inference is often performed using either synonymous or intronic sites, which are close to sites in coding regions, but the contribution of the effects of selection at linked sites are generally ignored. Patterns of variation in these regions may be skewed by the effects of either negative selection (Zeng 2013; Ewing and Jensen 2016) or positive selection (Messer and Petrov 2013), and this could strongly affect the accuracy of the inferred demographic model (Ewing and Jensen 2016; Schrider *et al.* 2016). In other words, selection may cause demographic parameters to be mis-estimated in such a way that population size changes are over or under-estimated.

In addition, the extent of BGS can vary considerably across the genome. Although it is necessarily a function of the number and selective effects of directly selected sites, as well as the rate of recombination (Hudson and Kaplan 1995; Nordborg *et al.* 1996; Charlesworth 1996, 2013), the interaction between these parameters and the underlying demographic history of the population remains poorly understood, even for simple models. Furthermore, existing analytical work (Zeng and Charlesworth 2010b; Zeng 2013; Nicolaisen and Desai 2013) has often been done under the assumption of demographic equilibrium, and is mostly restricted to describing strongly selected mutations with a fixed selection coefficient. However, population genomic data suggest the existence of a wide distribution of fitness effects of deleterious mutations (DFE), with a significant proportion of weakly selected mutations with 2*N*_e_*s* < 10 (reviewed by Bank *et al.* 2014b), where *N*_e_ is the effective population size and *s* is the reduction in fitness for mutant homozygotes. In regions of low crossing over, interference among such mutations may result in large distortions of the underlying genealogies (Kaiser and Charlesworth 2009; O’Fallon *et al.* 2010; Good *et al.* 2014), so that the consequences of a wide DFE are not well described by the analytical results.

We first investigate the joint effects of demography, the shape of the DFE, and the number of selected sites in shaping linked neutral variation. Next, we utilize the decay of BGS effects away from the targets of selection, by examining regions spanning coding / non-coding boundaries, in order to jointly infer the DFE of the coding region and the demographic history of the population. By performing extensive performance analyses and quantifying both power and error associated with our approximate Bayesian computation (ABC) approach (Beaumont *et al.* 2002), the method is shown to perform well across arbitrary demographic histories and DFE shapes. Importantly, by utilizing patterns of variation and divergence across coding and non-coding boundaries, this approach avoids the assumption of synonymous site neutrality inherent to approaches based on comparisons of nonsynonymous and synonymous site variability and divergence, an assumption that has been shown to be violated in many organisms of interest (Chamary and Hurst 2005; Lynch 2007; Zeng and Charlesworth 2010a; Lawrie *et al.* 2013; Choi and Aquadro 2016; Jackson *et al.* 2017), and which can result in serious mis-inference (Matsumoto *et al.* 2016). In applying this approach to genome-wide data from a Zambian population of *Drosophila melanogaster*, our results show that this population has experienced a very mild 1.2-fold growth in size, considerably less than previous estimates which did not correct for the BGS-induced skew of the SFS (*e*.*g*., Ragsdale and Gutenkunst 2017; Kapopoulou *et al.* 2018). In addition, we estimate that ∼25% of all mutations in exons are effectively neutral in this population, and find little evidence for wide-spread selection on synonymous sites.

## METHODS

### Simulations

The discrete generation simulation package SLiM 3.1 (Haller and Messer 2019) was used to simulate a functional element of length *L*, which is flanked by neutral non-functional regions. The functional region experiencing purifying selection is described by a DFE that is modeled as a discrete distribution with four bins (Figure 1a) representing effectively neutral (*γ* < 1), weakly deleterious (1 ≤ *γ* < 10), moderately deleterious (10 ≤ *γ* < 100), and strongly deleterious (100 ≤*γ*< 10000) classes of mutations, where *γ*= 2*N*_e_*s*. Semi-dominance is assumed, so that the fitness of mutant heterozygotes is exactly intermediate between the values for the two homozygotes (a dominance coefficient, *h,* of 0.5). Fitness effects are assumed to follow a uniform distribution within each of the four bins. In order to infer the extent of purifying selection, we estimated the fraction of mutations in each bin, referred to as *f*_0_, *f*_1_, *f*_2_ and *f*_3_, respectively (Figure 1a), such that 0 ≤*f*_i_ ≤ 1, and Σ_i_ *f*_i_ = 1, for *i* = 0 to 3. In addition, in order to limit the computational complexity, we restricted values of *f*_i_ to multiples of 0.05 (*i.e*., *f*_i_ ϵ {0.0, 0.05, 0.10, …, 0.95, 1.0} ∀ *i*). These constraints allowed us to sample 1,771 different DFE realizations, and to work independently of arbitrary assumptions regarding DFE shape.

**Figure 1:**
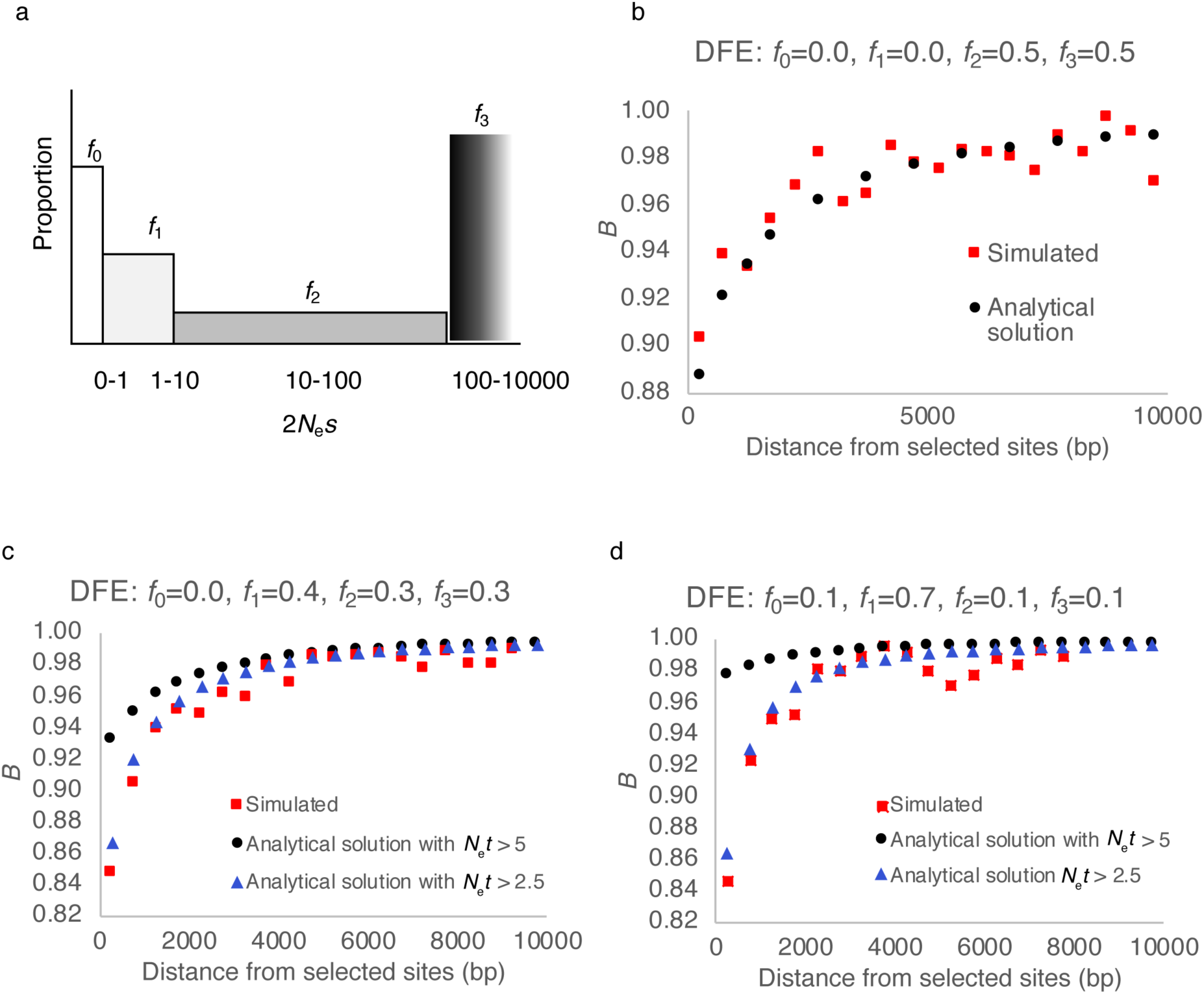
(a) An example of a discrete DFE with four classes of mutations. The proportion of each class of mutation, *f*_i_, lies between 0 and 1. (b) Nucleotide site diversity relative to the neutral expectation (*B* = *π*/*π*_0_) as a function of the distance from the directly selected sites (length 1 kb), as predicted by the analytical solution (black points) and as observed in simulations (red points). (c, d) Analytical predictions and simulated values for a DFE with larger contributions from the weakly deleterious class of mutations. Note that, for the analytical solutions, the two classes of results represent cases where mutations with 2*N*_e_*t* < 5 (black circles) and 2*N*_e_*t* < 2.5 (blue triangles) were ignored.

### Simulations under demographic equilibrium

Simulations were performed for 4 different values of *L* – 0.5 kb, 1kb, 5kb, and 10kb. The intergenic regions were assumed to be 10kb in length, and simulations were restricted to the intergenic region on one side of the functional region. For the purpose of power analyses and testing, we used population genetic parameter values that approximately resemble those for *Drosophila* populations. The population size in nature was assumed to be 10^6^ and the recombination rate (1 × 10^-8^ per site per generation) and mutation rate (1 × 10^-8^ per site per generation) were constant across the simulated region. Although we have not included gene conversion in this study, it will be an important addition in future studies. The simulations were performed with *N*_sim_ = 5000 diploid individuals, and the recombination and mutation rates were scaled proportionally to maintain realistic values of their products with *N*_e_. Such scaling is important for reducing computation time and has been found to be largely accurate, with some exceptions that are not relevant here (Comeron and Kreitman 2002; Hoggart *et al.* 2007; Kim and Wiehe 2009; Uricchio and Hernandez 2014).

We used a burn-in period of 80,000 generations, and an additional 20,000 (= 4*N*_sim_) generations were allowed for further evolution. For every set of parameter combination (*i.e*., *f*_0_, *f*_1_, *f*_2_ and *f*_3_) we performed 1000 replicate simulations.

### Simulations under non-equilibrium demography

Simulations with demographic changes were performed specifically to match the details of the *D. melanogaster* genome. A set of 94 exons belonging to the *D. melanogaster* genome were chosen according to certain criteria (see Results). Simulations were performed using the length of each exon, together with 4 kb of flanking intergenic sequence. The mutation rate was assumed to be 3.0 × 10^-9^ per site per generation, which is consistent with both pedigree (Keightley *et al.* 2014) and mutation accumulation (Keightley *et al.* 2009) studies, although somewhat higher mutation rates have been estimated in other studies (Schrider *et al.* 2013; Assaf *et al.* 2017). Ancestral and current population sizes were sampled from a uniform prior between 10^5^-10^7^ and with values of *f*_i_ chosen as described above. The nucleotide site diversity at 4-fold degenerate sites was 0.019 for the Zambian population of *D. melanogaster*, giving an estimate of *N*_e_ of 1.6 × 10^6^. A scaling factor of 320 corresponding to *N*_e_/*N*_sim_ (= 1.6 × 10^6^ / 5000) was used to perform all simulations with demographic changes. With population size changes, the time of change was assumed to be fixed at *N*_sim_ generations, as inferred in previous studies (Terhorst *et al.* 2017; Kapopoulou *et al.* 2018). A total of 10 replicates were performed for each exon, resulting in 940 replicates for every parameter combination. These simulations were conducted using the computational resources of Open Science Grid (Pordes *et al.* 2007; Sfiligoi *et al.* 2009).

### Calculation of summary statistics

First, we fitted a logarithmic function to the recovery of nucleotide site diversity (*π*) around the functional region such that *π* = *slope*\*ln(*x*) + *intercept*, where *x* is the distance of the site from the functional region in base pair. We used the *slope* and *intercept* of the fit to define the number of bases required for a 50%, 75%, and 90% recovery of nucleotide diversity, with 50% and below being defined as the “linked neutral” region and 50% and above as the “neutral” region. This analysis provides for three non-overlapping regions: (1) functional (experiencing direct selection), (2) linked-neutral (experiencing observable levels of background selection), and (3) neutral (experiencing low / unobservable levels of background selection). The following statistics were calculated for each of these three types of regions: nucleotide site diversity (*π*), Watterson’s *θ*, Tajima’s *D*, Fay and Wu’s *H* (both absolute and normalized), number of singletons, haplotype diversity, LD-based statistics (*r*^2^, *D*, *D*′), and divergence (*i.e*., number of fixed mutations per site per generation after the burn-in period). Simulations for any particular set of parameters were run with 1000 replicates and the means and variances of the above statistics across replicates were used as summary statistics for ABC. In addition to these variables, the six statistics summarizing the characteristics of the recovery of *π* in linked neutral regions were included. Together, these amount to 72 initial summary statistics. All statistics were calculated using the Python package pylibseq (Thornton 2003). The sample size was kept constant at 100 genomes (*i.e*., 50 diploid individuals). It should be noted that some statistics are strongly dependent on the number of sites used in the calculations, and the sizes of linked and neutral regions varied for every set of parameter combination, although this effect is captured in the individual prior distributions.

### ABC

We used an approximate Bayesian computation (ABC) approach, using the R package, “abc”(Csilléry *et al.* 2012), to co-estimate the DFE characterizing a functional region, as well as the population history. The relationship between the parameters and summary statistics were modeled with a linear regression method (ridge regression) and a non-linear correction regression method (a neural net), using default parameters provided by the package: https://cran.r-project.org/web/packages/abc/abc.pdf. The neural net method in the “abc” package by default fits a single-hidden-layer neural network with 5 units in the hidden layer. To infer posterior estimates, a tolerance of 0.05 was applied (*i.e*., 5% of the total number of simulations were accepted by ABC to estimate the posterior probability of each parameter). Cross-validation was performed by leaving out one randomly chosen simulation from which the summary statistics from that simulation were used to infer the parameters. A 100-fold cross-validation procedure was used to assess performance as well as to choose the tolerance value determining acceptance. The weighted medians of the posterior estimates for each parameter were used as point estimates.

### Ranking of summary statistics

Ranking of summary statistics was performed separately for both demographic equilibrium and non-equilibrium cases, using two different methods. The first approach consisted of performing Box-Cox transformations on all 72 summary statistics to correct for non-linear relations between statistics and parameters, using the function “boxcox” in R. Specifically, we used the code provided in Figure 9 of the ABCtoolbox manual (Wegmann *et al.* 2009) to find partial least squares components using R. The squared correlation coefficient, *r*^2^, between the transformed statistics and parameters was then used to rank each statistic for every parameter separately and a statistic was considered to be significantly correlated with the parameter if the *p*-value was less than 0.05 (Bonferroni corrected for multiple testing with a significance cutoff of 0.05/72). The second approach involved a modified version of the algorithm proposed by Joyce and Marjoram (2008) for ranking statistics. With this algorithm, we started with the entire set of 72 statistics. Each statistic was removed from the set and cross-validation using 20 randomly sampled simulations was used to identify the statistic that corresponded to the least error (*i.e*., the removal of which causes the least reduction in accuracy). The same algorithm was performed iteratively until only two statistics remained. This method was performed for each parameter separately, was replicated 10 times, and the average ranking across these replicates was used to obtain the final ranking. The second approach was extremely time consuming and was thus only used to rank the statistics for inferring the DFE under demographic equilibrium.

### Comparisons with DFE-alpha

Simulations were performed under demographic-non-equilibrium models, with 100 replicates of 94 exons each, and ancestral population sizes of 10,000 for all. Functional regions were simulated with 30% of sites being neutral, which were used to calculate the neutral SFS required by DFE-alpha. *Est_dfe* (Schneider *et al.* 2011) was used on the unfolded SFS to perform demographic inference and to infer the deleterious DFE. The proportion of adaptive mutations was fixed at 0.0. Final estimates of the DFE were obtained as *N*_w_*s* where *N*_w_ is the weighted population size inferred by *est_dfe*.

### *Drosophila* data application

Release 5 of the *D. melanogaster* genome assembly (Hoskins *et al.* 2007) and annotation version 5.57 were used, downloaded from ftp://ftp.flybase.net/genomes/Drosophila_melanogaster/dmel_r5.57_FB2014_03/gff/. Crossing over rates estimated by Comeron *et al*. (2012) for every exon and flanking intergenic region were obtained from the *D. melanogaster* Recombination Rate Calculator (https://petrov.stanford.edu/cgi-bin/recombination-rates_updateR5.pl) (Fiston-Lavier *et al.* 2010), and explicitly utilized for each specific region considered. These rates were halved to obtain sex-averaged rates of recombination (Campos *et al.* 2017) as all regions were restricted to autosomes. We excluded all genes that have a crossing over rate 5-fold larger or smaller than the average (*i.e.,* we used only genes with a crossing over rate of between 0.44 and 11 cM/Mb). Consensus sequences of all Zambia lines were downloaded from http://www.johnpool.net/genomes.html (Lack *et al.* 2015). IBD tracks and admixture tracks were masked using scripts provided by the same site. Individuals with known inversions were entirely excluded from the analysis (Kapopoulou *et al.* 2018).

The final set consisted of 76 haploid genomes. PhastCons scores calculated with respect to 15 insect taxa were downloaded from the UCSC genome browser (https://genome.ucsc.edu/). For each of the 94 exons, summary statistics were calculated using pylibseq (Thornton 2003) for the coding region and for 2kb intergenic regions flanking both sides. In order to exclude sites in intergenic regions that might be under direct selection, a phastCons cutoff score of 0.8 was used to calculate all statistics. That is, sites that had a greater than or equal to 80% probability of being a conserved noncoding element identified by phastCons, were excluded when calculating statistics.

For the purpose of inferring derived alleles and for calculating branch-specific rates of substitution, we used the ancestral sequence to the *D. melanogaster* genome provided to us by the authors of Kolaczkowski *et al.* (2011). The ancestral sequence reconstruction had been performed by maximum likelihood over 15 insect genomes available in the UCSC genome browser (Karolchik *et al.* 2004). Sites with missing ancestral sequence were excluded from analysis. Branch-specific rates of substitution (also referred to as divergence in this study) were calculated by identifying derived alleles that were fixed in the *D. melanogaster* Zambian population (*i.e.*, polymorphic sites were removed). After excluding sites with missing ancestral information, with IBD and admixture tracks, and which were likely to belong to a non-coding conserved element, we had on average 1062 sites per exon, 556 sites per linked region, and 666 sites per neutral regions.

It should be noted that for the purpose of performing inference using ABC, substitution rates in simulations were calculated per base pair for 25,000 generations. We thus normalized all rates obtained from simulations by the expected neutral substitution rate (*i.e*., *µ*_sim_*τ*_sim_ = 320 × 3×10^-9^ × 25000 = 0.024, where *μ*_sim_ is the scaled mutation rate and *τ*_sim_ is the number of generations used in the simulations for calculating divergence). Divergence estimates from *D. melanogaster* were normalized by an expected neutral substitution rate of *μτ* = 3 × 10^-9^ × 21333333 (the estimated divergence time) = 0.064 (Li *et al.* 1999; Halligan and Keightley 2006) where *μ* is the mutation rate in *D. melanogaster* and *τ* is the time to the ancestor of *D. melanogaster* and *D. simulans* in number of generations. In addition, inference was performed using divergence estimates only in the exonic regions. ABC inference for *Drosophila* was performed using the abc package in R, with linear regression aided by neural net with default parameters. Each inference was performed 50 times, and the mean of point estimates obtained were reported as the final parameter estimates.

### Data and code availability

The following data are publicly available on https://github.com/paruljohri/BGS_Demography_DFE: 1) Aligned sequences of the single-exon genes and their corresponding intergenic regions used in this study, including derived alleles and fixed substitutions; 2) Scripts to calculate statistics from simulations and from empirical data as well as the code used to perform simulations; 3) Values of all calculated statistics obtained for all parameter combinations; 4) A Mathematica notebook as well as a Python script for analytically calculating the reduction in linked neutral diversity caused by BGS caused by a functional element.

## RESULTS AND DISCUSSION

### Recovery of nucleotide diversity away from a functional region as predicted under equilibrium

The nucleotide site diversity at neutral sites linked to sites experiencing direct purifying selection, relative to its value in the absence of selection, (*B*), can be obtained by modifying Equation 6 of Nordborg *et al*. (1996), which is of the form:

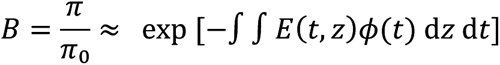

where *π*_0_ is the nucleotide site diversity in the absence of selection and *π* is its value in the presence of BGS. The term *E* in the exponent is a function of the heterozygous selection coefficient (*t* = *hs)* against a deleterious mutation at a selected site and the physical distance (*z*) between the neutral and selected sites. Here, *s* is the reduction in fitness of mutant homozygotes, and *h* is the dominance coefficient; *ϕ*(*t*) is the probability density function for *t*.

For the purpose of the current study, Equation S1a of the Supplementary Information of Campos and Charlesworth (2019) was modified to model a neutral site outside a gene, located *y* basepairs from the end of the functional region. If a selected site is *x* bp from the end of the gene (in the opposite direction), the distance between the two sites is *z* = *x* + *y.* The integral of *E*(*t*, *z*) over *z* is equal to:

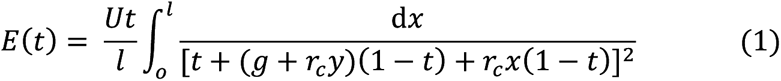

where *U* is the total mutation rate to deleterious alleles over the entire gene, *l* is the length of the gene in basepairs, *g* is the rate of gene conversion, and *r*_c_ is the rate of crossing over per basepair. The crossover map is assumed to be linear, so that the net rate of recombination between the two sites is *g* + *r*_c_*z*, and *z* is assumed to be sufficiently large that the effect of gene conversion is independent of *z*.

By evaluating the integral in Equation 1, we have:

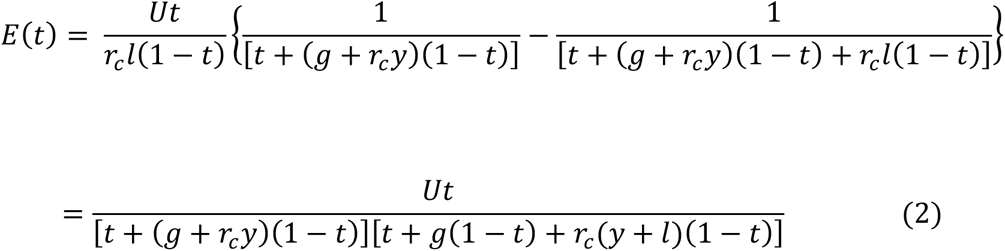

Note that this equation implies that, if *t* is small compared with *y*, BGS effects outside the coding region will be minimal.

We can integrate *E*(*t*) over the distribution of selection coefficients as described in the Appendix. The expectation of *E*(*t*) for a given bin of *t* values is then given by the sum of the following two terms:

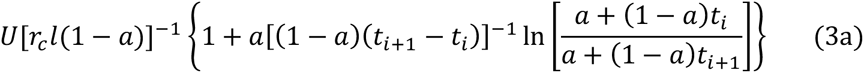

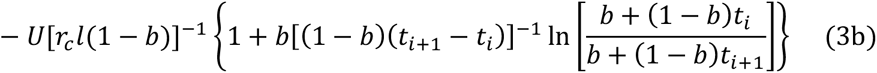

where *a* = *g* + *r_c_y* and *b* = *g* + *r_c_*(*y* + *l*), and the *t*_i_’s correspond to the boundary of the discrete bins. For the case when *b* << 1, the sum of the two components is approximately equal to:

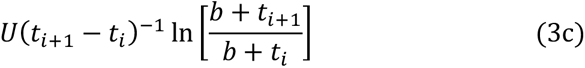

Figure 1b shows the theoretical and simulation results for *r*_c_ = 10^-6^, *l* =1000, *U* = *lµ*, *µ* = 10^-6^, *g* = 0, *t*_0_ = 0, *t*_1_ = 0.00005, *t*_2_ = 0.0005, *t*_3_= 0.005, and *t*_4_ = 0.5. It should be noted that these derivations assume that *N_e_t* >>1, which is violated by the presence of the weakly deleterious DFE class (frequency *f*_1_). Most studies deal with this assumption by ignoring the contribution of mutations with *N*_e_*t* < 5 or 10 (Charlesworth 2013; Elyashiv *et al.* 2016; Torres *et al.* 2019).

As expected from these considerations, when all classes of mutation were included we found a significant discordance between the simulated and theoretically predicted values for the slope of the recovery of diversity as *f*_1_ increases (Figure 2c and 2d, Table S1). On including only mutations with *N*_e_*t* > 2.5 (*i.e.*, *γ* = 2*N*_e_*s* > 5), the diversity patterns are mostly well explained, even when the DFE is highly skewed towards the weakly deleterious class. In fact, it is interesting to note that a combination of high values of *f*_1_ and *f*_2_ can result in BGS effects that extend up to 4 kb, even for very short exons, although the maximum reduction in diversity is around 10-15%, which is consistent with the findings of Charlesworth (2012) and Campos and Charlesworth (2019).

**Figure 2:**
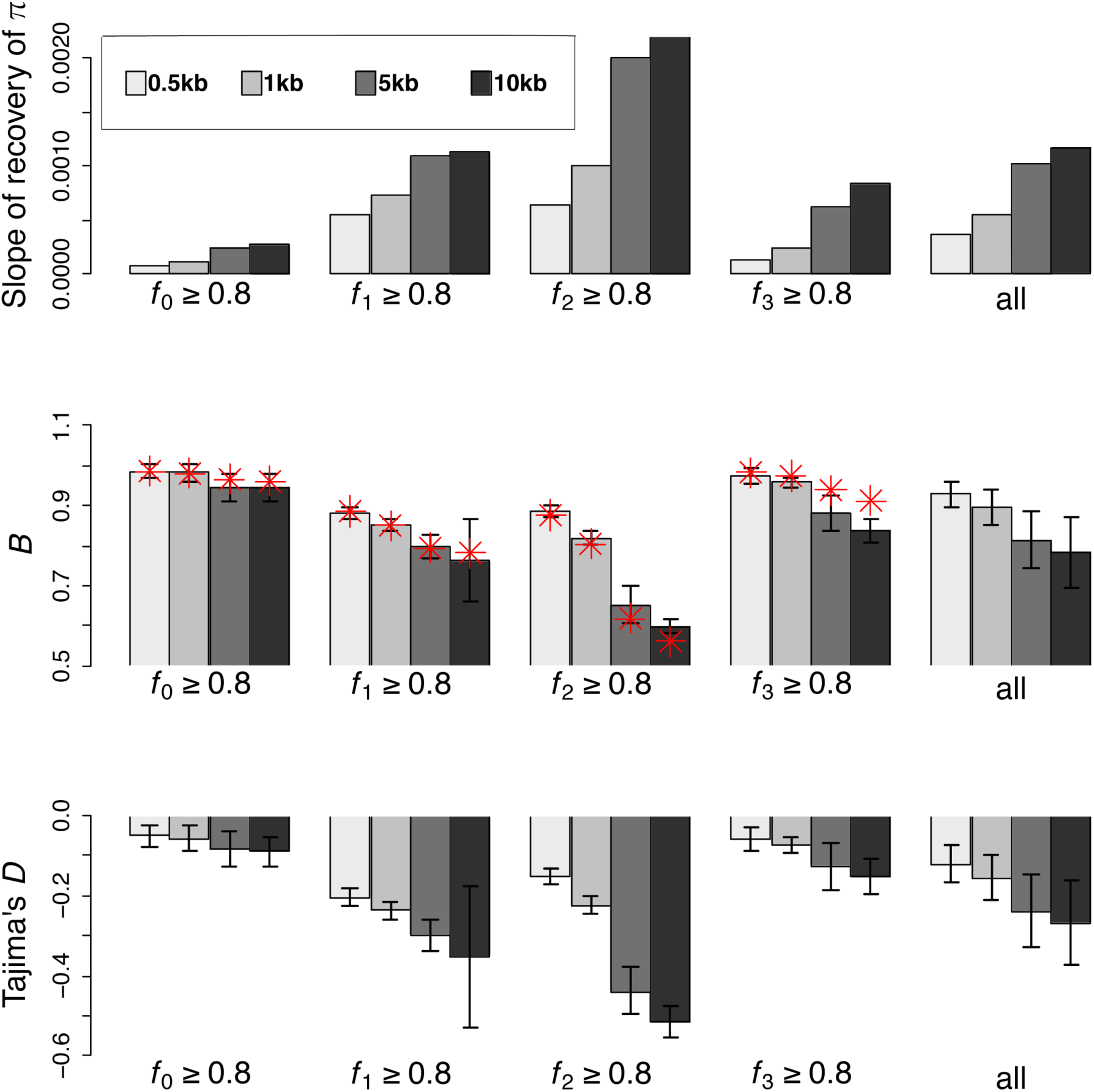
Effects of BGS under demographic equilibrium. (a) The slope of the recovery of nucleotide diversity in 10kb linked neutral regions flanking functional regions, such that π = *slope*\*ln(*distance from functional region*) + *intercept*, (b) nucleotide diversity in 500bp linked neutral regions flanking functional regions relative to neutral expectation (*B*), and (c) Tajima’s *D* for 500bp linked neutral region flanking functional regions. All of the above are shown for various sizes of functional elements (0.5-10 kb) and DFE shapes. The four DFE shapes considered are *f*_i_ ≥ 0.8 for *i* = 0,1,2,3, with more than 80% of mutations residing in DFE class *f*_i_, such that ∑ *f*_j_ ≤ 0.2, where *j*≠*i*. The DFE category “all” represents an average over all possible DFE shapes. The error bars are 2 × standard deviation. Red points show the analytical predictions for *B* with (1) *f*_0_=0.85, *f*_1_=0.05, *f*_2_=0.05, *f*_3_=0.05, (2) *f*_0_=0.05, *f*_1_=0.85, *f*_2_=0.05, *f*_3_=0.05, (3) *f*_0_=0.05, *f*_1_=0.05, *f*_2_=0.85, *f*_3_=0.05, and (4) *f*_0_=0.05, *f*_1_=0.05, *f*_2_=0.05, *f*_3_=0.85.

### Joint effects of demography, the DFE and the number of selected sites on linked neutral variation

While the above results show that the effect of BGS on linked neutral regions can be determined analytically, there are several reasons for investigating BGS effects using simulations. First, the analytical expressions neglect the contribution of very weakly deleterious mutations (*N*_e_*t* < 2.5), and do not predict the site frequency spectrum of variants (the SFS). In addition, they assume demographic equilibrium, which is probably not true of natural populations.

#### Effects of the shape of the DFE and number of selected sites

We first simulated 10kb neutral regions linked to functional regions of varying sizes, 0.5kb, 1kb, 5kb, and 10kb, assuming demographic equilibrium, as shown in Figure 2. By varying the contributions from each bin of selective effects, with frequencies *f*_0_, *f*_1_, *f*_2_ and *f*_3_, it was possible to sample all possible DFE shapes, as described in the Methods section. As expected from Equation 3c, the reduction in diversity is non-linearly proportional to the number of selected sites for a given recombination rate. A larger number of selected sites increases both the total reduction in diversity and the slope of the recovery of diversity away from functional regions (Figure S1). The maximum reduction in diversity in the linked neutral regions (immediately adjacent to the functional region), averaged across all DFE realizations, is approximately 8%, 12%, 24%, and 29% for 0.5kb, 1kb, 5kb, and 10kb selected sites, respectively. Furthermore, for the chosen recombination rate, the median numbers of base pairs necessary to achieve a 50% recovery in diversity are 955, 1035, 1350, and 1650 bp, respectively (Figure 2a).

The reduction of nucleotide diversity at closely linked neutral regions was maximized when the proportion of weakly deleterious mutation (*f*_1_) and moderately deleterious mutations (*f*_2_) was largest (Figure 2b, Table S2). The effect is greatest when purifying selection is weak, allowing mutations to segregate in the population prior to being purged (Campos *et al.* 2017). Although weakly deleterious mutations (*f*_1_) only reduce variation slightly, they generate significant distortions in the SFS (Figure 2c), consistent with previous studies (Charlesworth *et al.* 1995; Charlesworth 2012; Nicolaisen and Desai 2013). Moderately deleterious mutations cause the largest reduction in *π*, the highest rate of recovery of *π* around functional regions, and the largest skew in the SFS towards rare variants. As expected, the proportion of strongly deleterious mutations (*f*_3_) does not greatly affect levels of linked neutral variation, and these mutations skew the SFS only slightly. Furthermore, increasing the number of selected sites results in larger BGS effects for all DFE types, as is to be expected. It should be noted that these generalizations about BGS effects depend on the distance between the neutral and selected sites; in particular, the size of the region affected by deleterious mutations is expected to be an increasing function of the size of their fitness effects. As we were interested in understanding BGS effects caused by all classes of mutations, we focus our further discussion on sites close to the functional boundary, where all classes of mutation are likely to have an impact.

To summarize, at demographic equilibrium, neutral regions linked to functional regions under selection undergo a reduction in diversity and a skew in the site frequency spectrum, both of which depend on the underlying shape of the DFE and the number of directly selected sites (Charlesworth *et al.* 1993, 1995; Charlesworth 2013; Campos and Charlesworth 2019). Importantly for the sake of statistical inference, however, the three classes of deleterious mutation behave differently, suggesting the possibility of distinguishing their relative contributions (as discussed in the next section). Furthermore, these results again demonstrate the potentially important role of BGS in shaping patterns of neutral variation, highlighting the danger posed by ignoring these effects when performing demographic inference (see Ewing and Jensen 2016). Additionally, the dramatic differences in the extent of background selection effects as a function of the number of directly selected sites emphasize the necessity of directly modeling exon sizes in empirical applications.

#### Effects of demography and the shape of the DFE on background selection

We investigated the effects of BGS after recent changes in population size. Populations with the same ancestral population size (*N*_anc_) either experienced 10-fold exponential growth or contraction in the last 4*N*_anc_ generations; BGS effects were compared to populations that remained in equilibrium throughout, for all possible DFE shapes. Both expansion and contraction result in reduced BGS effects (*i.e*., there is an increase in *B* compared to equilibrium), irrespective of the shape of the DFE (Figure 3a, b). This observation suggests that the extent of BGS caused by functional elements is not only determined by the strength of selection, but also by the demographic history of the population. Thus, demographic effects may in principle explain variable inferences among studies of the importance of purifying selection in shaping genome-wide patterns of variation (Cutter and Payseur 2013).

**Figure 3:**
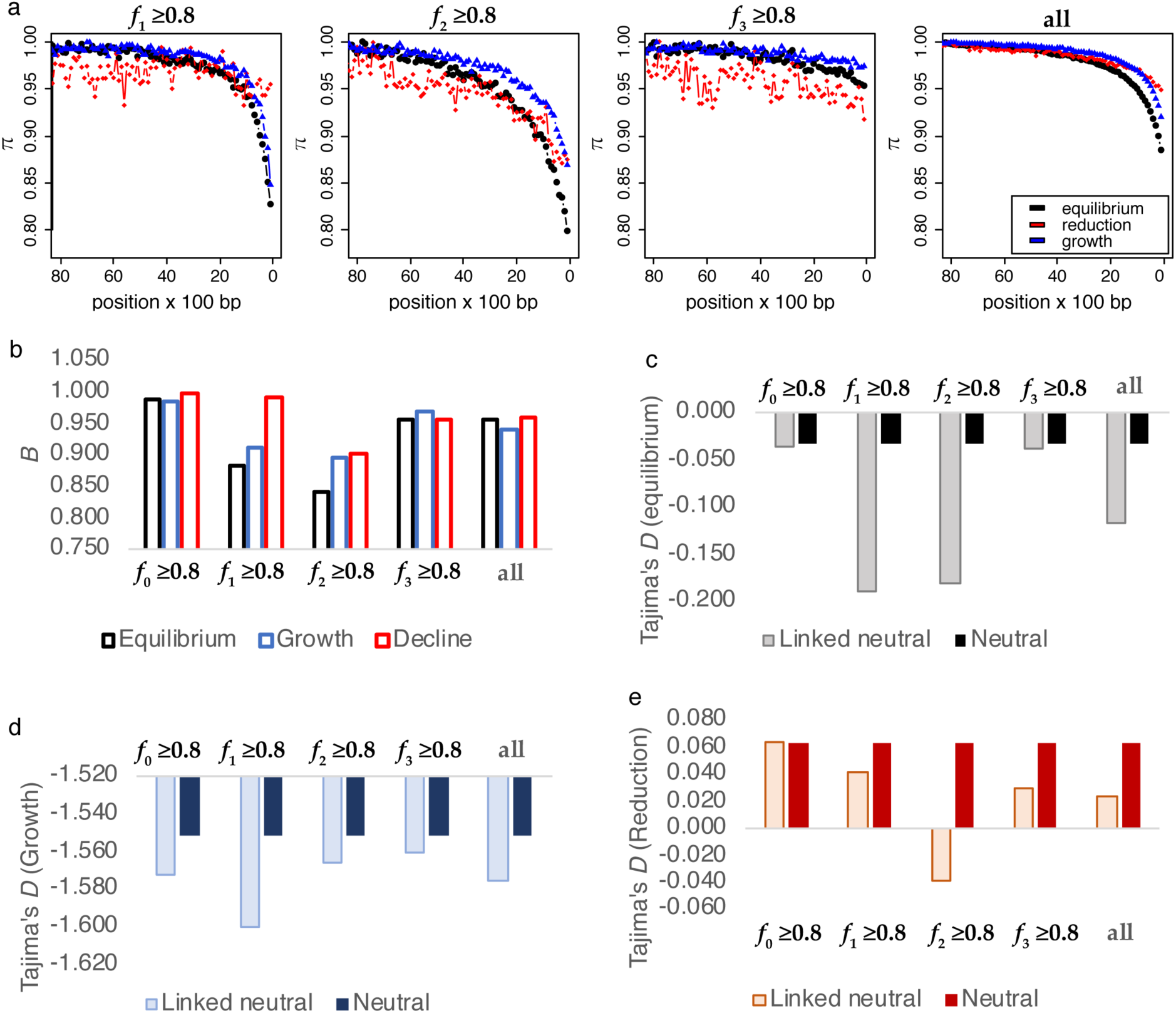
Effects of BGS under non-equilibrium demography. (a) The slope of recovery of nucleotide diversity in linked neutral regions for different DFE shapes under equilibrium demography (black), population expansion (blue), and contraction (red). (b) Nucleotide site diversity relative to neutral expectation (*B*), over 500 bp of linked neutral regions flanking functional regions, for varying DFE shapes and three different demographic models - equilibrium (black), 10-fold exponential expansion (blue), and 10-fold exponential decline (red). (c) Tajima’s *D* for the 500 bp linked neutral region flanking the functional region under equilibrium, (d) after a 10-fold expansion, and (e) after a 10-fold population size reduction. The four DFE shapes considered in all panels are *f*_i_ ≥ 80% for *i* = 0,1,2,3, where more than 80% of mutations reside in DFE class *f*_i_. The DFE category “all” represents an average over all possible DFE shapes. For non-equilibrium demography, γ = 2*N*_anc_*s*, where *N*_anc_ is the ancestral population size.

Interestingly, however, there is still a significant skew in the SFS at linked neutral sites caused by BGS after a population size change (Figure 3c-e). Thus, in more compact genomes, where background selection is pervasive, this suggests that methods which use the SFS to fit demographic models may overestimate growth and either underestimate population contraction or mis-classify contraction as expansion. It is also interesting to note that BGS effects are largest under demographic equilibrium, such that constant population size is likely to be inferred as population growth.

### Inference of the DFE under demographic equilibrium

The next question we investigated was whether the parameters of the DFE can be estimated using the set of summary statistics described in the Methods section. We first determined whether it is possible to distinguish the four different classes of the DFE under demographic equilibrium, using population genomic data and divergence from the closest outgroup species. The simulations involved functional regions of lengths *L* = 0.5kb, 1kb, 5kb and 10kb, with linked neutral regions of 10kb and a discrete DFE as described previously. The ABC approach described in the Methods section was used to quantify our ability to infer the four DFE parameters. The recovery of nucleotide diversity over linked neutral regions was used to calculate the distance in basepairs (*π*_50_) required for diversity to reach 50% of its maximum value (see Methods). The neutral region within *π*_50_ base pairs from the functional region was defined as “Linked”, and the remainder was defined as “Neutral” (Figure 4a). Statistics were calculated for three regions (Functional, Linked, and Neutral) separately and the means and variances across simulation replicates of each statistic were used to infer the four parameters. The simulation replicates correspond to independently evolving loci within a genome. In the following sub-sections, we describe the performance of the method and its robustness to various model violations.

**Figure 4:**
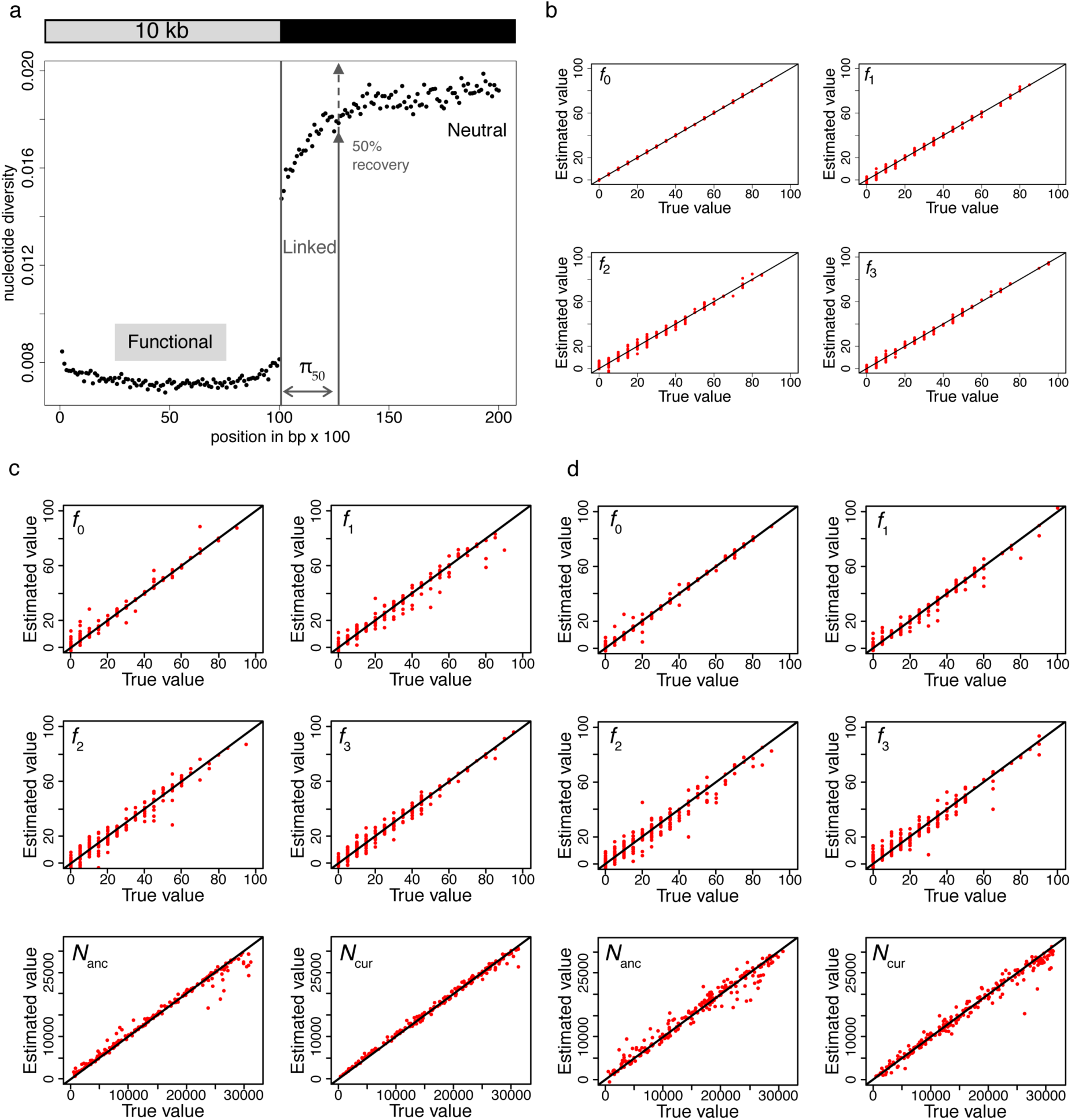
(a) Values of diversity statistics across functional, linked and neutral regions. (b) Accuracy of estimation (cross validation) of the four classes of the DFE using statistics for functional regions only (size 1kb), under equilibrium demography. (c) Joint estimation of population size changes and the DFE using all statistics. (d) Joint estimation of population size changes and the DFE using statistics for functional regions only. The true proportions of mutations in each DFE class and *N*_anc_ and *N*_cur_ are given on the X-axes, while the estimated values are given on the Y-axes. Parameters are indicated on the upper left corner for each plot. Each dot represents one out of 200 different parameter combinations, sampled randomly from the entire set of simulations.

#### Accuracy of inference

All four DFE classes were estimated fairly accurately when using all statistics (Figure S2a). However, under demographic equilibrium, the DFE is inferred much more accurately using statistics from the functional regions alone, thus side-stepping the need for the identification of linked neutral regions (Figure 4b, Figure S2b). In both cases, the accuracy of inference is highest for the neutral class and lowest for moderately deleterious mutations (class 2), and significantly improves when the size of the functional region increases (Figure S2). When using only functional regions to perform inference, the absolute difference between the true value and the estimated value of the neutral class is approximately 0.034, 0.030, 0.017, and 0.010 for functional sizes of 0.5kb, 1kb, 5kb and 10 kb. For 1kb regions, therefore, the method cannot determine whether the neutral class of mutations comprise 30% or 33% of the DFE. For the moderately deleterious class, this error is larger – 0.077, 0.060, 0.028, and 0.019, respectively. These absolute error values are not surprising, as the *f*_i_ in our simulations are multiples of 0.05 out of computational necessity. The accuracy of the estimates can thus be increased by sampling the parameter space more densely.

The accuracy of estimation can also be evaluated using *r*^2^ between the true and estimated values. For instance, for 1kb functional regions, the *r*^2^ values for *f*_0_, *f*_1_, *f*_2_ and *f*_3_ are 0.93, 0.91, 0.89, and 0.87 respectively. It is interesting to note that it is possible to infer the proportion of DFE classes when using statistics from the linked region alone (Figure S3). Although there is an increase in the absolute errors for 10kb regions to 0.103, 0.122, 0.056, and 0.044 in *f*_0_, *f*_1_, *f*_2_ and *f*_3_, respectively, this analysis suggests that, if the population size were known to be at equilibrium, statistics for the linked neutral regions alone could distinguish between the contributions of the four DFE classes.

It should furthermore be noted that this approach does not distinguish between non-synonymous and synonymous mutations. Indeed, no assumption is made regarding which specific bases are neutral, nearly neutral, or deleterious in the coding region. Thus, this method can be used to estimate the DFE for any type of functional region, as well as to assess the non-neutrality of synonymous sites by comparing their frequency in a given coding region with the value of *f*_0_.

#### Effect of mis-specification of exon size and recombination rate

In view of these results, it is important to consider whether accurate estimates depend on correctly specifying precise exon size, or if it would be sufficient to generate priors assuming, for example, a mean exon length for a given genome. To quantify this effect, simulated data sampled from the priors was based on 1kb exons, while the test data were obtained from simulations based on alternative exon sizes. The error in inference of the DFE increases as the difference between exon sizes of the priors and that of the true sizes are increased (Figure S4), with the highest error for the moderately deleterious class 2, although when exon sizes are sufficiently large, mis-specification of exon size does not strongly affect performance. A similar approach was used to determine if the presence of another functional region (also 1kb in size), separated by an intron or intergenic region, would skew inference. As expected, smaller intron sizes result in stronger mis-inference than larger ones, and intronic/ intergenic sizes larger than 4 kb performed essentially as well as those with no nearby functional exon (Figure S5). Moreover, a two-fold difference between assumed and actual recombination rates inflated the errors dramatically (Figures S6 and S7). Informatively, the direction of bias generated differs by DFE class (Figure S7). For example, when true recombination rates are half of those assumed, the inferred weakly deleterious class is greatly inflated. As this class of mutations most strongly skews the linked neutral SFS, this mis-inference presumably arises from an attempt to fit stronger linked effects by inferring a higher proportion of mutations in this class, whereas in reality the increased BGS effects are being generated by fewer recombination events than are assumed.

These results highlight the importance of taking into account the specific exonic-intronic-intergenic structure of a particular genomic region of interest, nearby functional regions and the specific recombination rate. Although any configuration of these details may be directly simulated, an alternative approach is simply to group exons of like size across a genome, and further reduce these to a group that is devoid of neighboring functional regions.

### Joint inference of purifying selection and demography, under non-equilibrium conditions

Based on the above results demonstrating that details of exon sizes and recombination rates are essential for accurate inference, we explicitly modeled both exon sizes and recombination rates when examining our ability to jointly infer demographic changes together with the DFE. As our example involved an African population of *D. melanogaster*, we chose for our simulations single-exon genes that had more than 4kb non-coding regions flanking both sides and whose exon sizes were between 500-2000bp. For this specific set of 94 exons, we simulated functional regions with specified exon sizes linked to 4kb neutral regions and utilized the previously inferred local crossing over rate for each exonic region in question. For every parameter combination, we performed 10 replicates of each of the 94 exon sizes (resulting in a total of 940 replicates per parameter combination), with their respective recombination rates and exon sizes, and summarized the resulting means and variances of the summary statistics.

Models of exponential population size expansion and contraction assumed various ancestral population size (*N*_anc_) and current population size (*N*_cur_), which were both sampled uniformly between 10^5^ – 10^7^, following previous studies (Duchen *et al.* 2013; Arguello *et al.* 2019). As earlier work has inferred the duration of the expansion in Zambian populations to be of the order of *N_e_* generations, this was scaled down and fixed at *N_sim_* = 5000 generations, in order to attempt to infer both historical and current population sizes. Thus, for this framework, we evaluated the estimates of six parameters: *f*_0_*, f*_1_*, f*_2_*, f*_3_*, N*_anc_, and *N*_cur_.

#### Accuracy of joint inference

Encouragingly, the results demonstrated an ability to successfully co-estimate the DFE and both ancestral and current population sizes, using the set of coding and linked non-coding summary statistics described above (Figure 4c). Under non-equilibrium demography, the estimation error for the strongly deleterious class of mutations is larger. The absolute differences between true and estimated values were 0.019, 0.027, 0.033, and 0.034 for the four DFE classes, respectively; the errors in ancestral and current sizes were 10.1% and 7.3% respectively. The *r*^2^ values between the true and estimated values of *f*_0_, *f*_1_, *f*_2_, *f*_3_, *N*_anc_, and *N*_cur_ were 0.97, 0.97, 0.95, 0.95, 0.99, and 0.99, respectively.

The performance of the full 6-parameter estimation procedure is good, without relying on the usual step-wise approach of first utilizing putatively neutral sites to fit a demographic model, and then using this model to estimate DFE parameters. Interestingly, joint estimation is almost as accurate when using statistics from functional regions alone (Figure 4d), although it inflates the errors in the estimates of *f*_2_ and *f*_3_. The absolute differences between the true and estimated values of *f*_0_, *f*_1_, *f*_2_, and *f*_3_ were 0.015, 0.025, 0.054, and 0.049, respectively, while the error in estimates of population sizes increases to 23% and 8% for *N*_anc_ and *N*_cur,_ respectively. Thus the error in ancestral population size is quite large if only functional regions are used to co-estimate all six parameters. Further, unlike the case of demographic equilibrium, statistics in linked regions alone can no longer be used to accurately infer parameters of the DFE (Figure S8).

#### Effect of mis-specification of mutation rate

We evaluated the effect of having incorrect estimates of mutation rate by inferring all six parameters under a scenario in which the assumed mutation rate is half or twice the true value. Under all demographic scenarios, if the assumed mutation rate was half of the true rate, our method correctly estimated *f*_0_, *f*_1_ and *N*_cur_ (Figure S9). However, the strongly deleterious class 3 is consistently under-estimated, the moderately deleterious class 2 is over-estimated, and *N*_anc_ is biased towards a strong population size decline. A comparable magnitude of mis-inference is observed when the assumed mutation rate is twice the true value (Figure S9). Thus, the ABC method described in this study is sensitive to large mismatches between the true and assumed mutation rates, and is thus best suited to organisms in which pedigree and/or mutation accumulation-based estimates are available.

### Statistics important for distinguishing different classes of the DFE and demography

As it is important to understand which statistics are needed for distinguishing between the effects of demography and the different classes of the DFE, two different approaches were used to rank statistics by their importance. First, statistics were simply ranked by their regression coefficient with respect to each parameter separately. Non-linear relationships were taken into account by using Box-Cox transformation, as suggested by Wegmann *et al*. (2009). With stationary population size, most of the top predictors of the fraction of neutral (*f*_0_) and strongly deleterious (*f*_3_) sites are statistics summarizing the functional region (Table S3). The top four statistics for each parameter are displayed in Figure S10. In addition, a modification of the method of Joyce and Marjoram (2008) was also employed to rank statistics (Table S4) for equilibrium demography.

As expected, the statistics that correlate most strongly with the fraction of neutral mutations are the levels of divergence and the fraction of high frequency derived alleles, as summarized by *θ*_H_ (Fu 1995; Fay and Wu 2000) in functional regions. As the weakly deleterious class of mutations generate BGS effects at closely linked sites, statistics for the functional and linked region are most strongly correlated with *f*_1_. This also correlates most with *H*′ in functional regions, a statistic that contrasts the proportion of high frequency derived variants with those of derived variants segregating at intermediate frequency (Fay and Wu 2000). Although this statistic was designed to identify selective sweeps, which tend to increase the proportion of high frequency derived alleles, it is highly predictive of the fraction of weakly deleterious class of mutations in the absence of positive selection. As shown above, larger *f*_1_ generates a stronger skew in the linked neutral SFS towards rare variants and is thus also reflected in the values of Tajima’s *D* in the linked neutral region. Measures of linkage disequilibrium in the functional and linked neutral regions are also correlated with the frequency of the weakly deleterious class of mutations.

Because the moderately deleterious class of mutations generates BGS effects that extend for larger distances than the more weakly selected class, the strongest correlates of this class are generally statistics from the neutral region furthest from the directly selected sites. All the different summaries of the SFS - *θ_W_*, *π*, and *θ*_H_ - correlate with this parameter, as well as the total reduction in linked neutral diversity (given by the intercept of the regression fit of *π* = *slope*\*ln(*distance*) + *intercept*, where *π* is the diversity in linked neutral regions). The strongly deleterious class of mutations is correlated with the number of singletons and *θw*.

A similar analysis was performed on simulations with demographic non-equilibrium. Here, the DFE parameters are significantly correlated only with the statistics for functional regions (Table S5 and S6). As expected intuitively, the statistics most highly correlated with the two demographic parameters are for the neutral linked regions. Ancestral population sizes correlate best with statistics that capture high frequency derived alleles in linked neutral as well as functional regions, as these represent older mutations; current population sizes correlate most with statistics that summarize LD. The same is true when ranked statistics are obtained only from functional regions. Because the class of moderately deleterious mutations and ancestral population sizes are correlated with overlapping sets of statistics, the estimates of these two parameters are partially confounded. As such, LD-based statistics are essential for distinguishing between demography and purifying selection, and in estimating ancestral and current population sizes. In addition, although the variances and means of the statistics are highly correlated, the variances play a more important role in estimating current population sizes.

### Comparison with DFE-alpha

Because there are no other programs that simultaneously co-estimate both demographic and selection parameters, we compared the performance of our method to the step-wise approach of DFE-alpha (Keightley and Eyre-Walker 2007; Eyre-Walker and Keightley 2009; Schneider *et al.* 2011), a program used widely for the inference of the DFE. DFE-alpha assumes that synonymous sites are neutral and uses their SFS to infer changes in population size. Conditional on the inferred demography and under the assumption that the deleterious selection coefficients follow a given distribution (generally gamma), the program infers the shape and scale parameter of the assumed distribution.

We simulated demographic equilibrium, 2-fold population growth and 2-fold population contraction, and inferred the change in population size and the DFE using both ABC and DFE-alpha. Because DFE-alpha uses neutral sites to infer demography, in all cases we simulated a DFE consisting of 30% neutral mutations, which are a proxy for synonymous sites. These simulations were performed exactly as described previously for non-equilibrium conditions. Exons sizes between 500-2000 bp with flanking 4 kb linked neutral regions were simulated with recombination rates specific to the selected 94 exons (and a total of 940 replicates for every parameter combination). DFE-alpha performs slightly better than ABC if the true DFE is indeed gamma distributed (Figure S11) although our method is able to infer the DFE with very similar accuracy.

For a discrete DFE which is skewed towards highly deleterious mutations, DFE-alpha and ABC perform with similar accuracy. However, our method performs better if the DFE is skewed towards slightly deleterious mutations as shown in Figure S12. It is important to note that, for the purpose of this comparison, simulations were run with numbers of directly selected sites between 500–2000 bp, with 30% of mutations being neutral, because neutral mutations are required to estimate demographic parameters with DFE-alpha. Under these conditions, background selection results in a relatively small skew in the neutral SFS (Campos and Charlesworth 2019).

As noted previously, a potential advantage of the methodology proposed here is that, by simultaneously estimating selection and demography, one is not required to make any assumptions about the neutrality of synonymous sites. We evaluated this feature by simulating a scenario where 33% of the assumed neutral sites were actually experiencing weak direct selection. As weak purifying selection generates a larger fraction of rare variants than stronger selection, programs based on neutrality would be likely to falsely infer growth. As expected, DFE-alpha inferred 2-fold growth under demographic equilibrium, and in fact inferred slight growth even for a 2-fold contraction (Figure 5). The resulting DFE over-estimated the fraction of neutral mutations and under-estimated the fraction of weakly deleterious mutations. As noted previously, such mis-inference will increase with the density of selected sites. Our ABC approach, however, accurately estimated the proportion of neutral mutations present in the selected region (Figure 5), illustrating the importance of joint inference.

**Figure 5:**
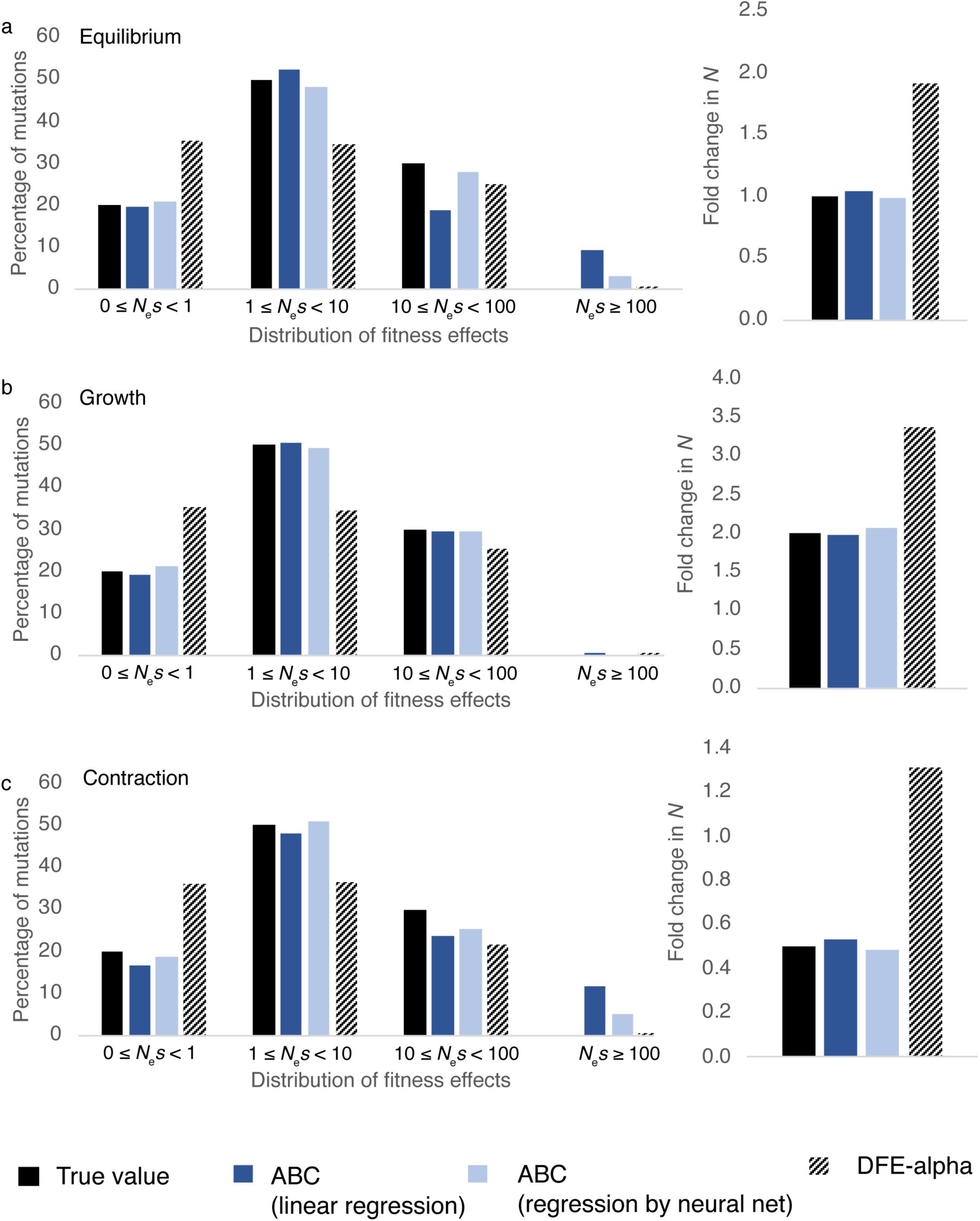
Comparison of the performance of the proposed ABC approach in the current study with DFE-alpha, under (a) demographic equilibrium, (b) exponential growth, and (c) exponential decline. In all cases, 30% of sites were assumed to be synonymous, out of which 33% were weakly selected. Solid black bars are the true simulated values, dark blue bars give the ABC performance using ridge regression, and light blue bars give the ABC performance using linear regression aided by neural nets. Patterned bars show the performance of DFE-alpha. A total of 998,300 sites were analyzed in the functional region for each parameter combination, with approximately 332,767 representing synonymous and 665,533 representing nonsynonymous sites.

### Application to *Drosophila melanogaster*

When simulating genomic regions, the presence of nearby coding regions that are not included in the models can generate additional BGS effects and thus bias inference. We thus restricted our analyses to protein-coding exons in the *D. melanogaster* genome between 500 to 2000 bp in length that are single exon genes, and are flanked on both sides by intergenic regions larger than 4 kb, so that effects of linkage with other nearby functional elements are avoided. It should be noted that any genic structure could readily be chosen for inference by directly simulating the associated details when constructing the priors - we have simply chosen this realization in order to provide an illustrative application.

The recombination rates of both the 5′ and 3′ flanking intergenic regions are highly correlated (Figure S13) and span a considerable magnitude (Figure S14), with a mean rate of 2.78 cM/Mb (*i.e.,* the average recombination rate for these chosen single exon genes is very near the autosomal genome-wide average of 2.32 cM/Mb). We also verified that this set of genes was not unusual with respect to genome-wide coding sequence divergence (Figure S15). Furthermore, because sites in intergenic regions in *D. melanogaster* may also experience direct selection (Halligan and Keightley 2006; Casillas *et al.* 2007), we used phastCons scores to exclude intergenic sites that might potentially be functionally important. All sites with a phastCons score larger than 0.8 were excluded (Siepel *et al.* 2005).

Table 1 provides the observed summary statistics for each region class, where intergenic sites that had a greater than or equal to 80% probability of belonging to a conserved element (*i.e*., with phastCons score ≥ 0.8) were excluded. It should be noted that the absence of a large difference between divergence (*i.e*., number of fixed substitutions specific to *D. melanogaster*) in exonic vs intergenic regions is consistent with previous studies (Table 1 in Andolfatto 2005). In addition, because we have restricted our analyses to sites where the ancestor of *D. melanogaster* could be predicted with high confidence, our analyses may be skewed towards more conserved sites, potentially resulting in lower divergence in intergenic regions. Previous estimates of divergence along the *D. melanogaster* lineage at 4-fold degenerate sites were approximately 0.05-0.06 (Halligan and Keightley 2006; Langley *et al.* 2012; Charlesworth *et al.* 2018), while that in coding regions was 0.023 (Langley *et al.* 2012). Although our estimates are lower than previous estimates, this discrepancy is explained by the larger number of individuals used to subtract polymorphic sites in this study (Table S7). With a sample size of 1 allele (corresponding to pairwise divergence), our estimates of divergence at 4-fold degenerate and in coding regions are 0.05 and 0.023, respectively, consistent with previous studies. In addition, a very similar relation between pairwise divergence and polymorphism-adjusted divergence is found with simulated data (Table S8).

**Table 1:** Statistics calculated for the 94 single-exon genes including their 3′ flanking intergenic sequences, for 76 haploid genomes (devoid of any inversion) from the Zambian population of *D. melanogaster*. Sites with phastCons scores higher than 0.8 were excluded. Functional refers to exons, linked refers to intergenic region (∼ 1kb) adjacent to exons and neutral refers to intergenic regions further away from exons that are adjacent to linked regions (Figure 4a). Derived alleles were identified by polarizing alleles with respect to the ancestral sequence of *D. melanogaster* obtained from ancestral reconstruction over 15 insect species.

|  | mean |  |  | standard deviation |  |  |
| --- | --- | --- | --- | --- | --- | --- |
|  | functional | linked | neutral | functional | linked | neutral |
| $\pi$ | 0.0083 | 0.0106 | 0.0107 | 0.0039 | 0.0042 | 0.0038 |
| $\theta_w$ | 0.0120 | 0.0166 | 0.0162 | 0.0045 | 0.0053 | 0.0049 |
| $\theta_H$ | 0.0088 | 0.0098 | 0.0097 | 0.0054 | 0.0053 | 0.0056 |
| $H'$ | -0.0633 | 0.0871 | 0.1169 | 0.5371 | 0.4118 | 0.3829 |
| Tajima's $D$ | -1.0537 | -1.1469 | -1.1103 | 0.5338 | 0.4874 | 0.4694 |
| Singleton density | 0.0215 | 0.0303 | 0.0307 | 0.0086 | 0.0116 | 0.0117 |
| Haplotype diversity | 0.9711 | 0.9680 | 0.9762 | 0.0452 | 0.0458 | 0.0444 |
| $r^2$ | 0.0328 | 0.0364 | 0.0363 | 0.0109 | 0.0136 | 0.0128 |
| $D$ | 0.0005 | 0.0005 | 0.0006 | 0.0009 | 0.0010 | 0.0012 |
| Branch-specific divergence | 0.01378 | 0.0156 | 0.0159 | 0.0075 | 0.0077 | 0.0071 |

Interestingly, although previous studies have inferred approximately 2-4-fold growth in the Zambian population of *D. melanogaster* (Ragsdale and Gutenkunst 2017; Kapopoulou *et al.* 2018), we infer only a 1.2-fold growth, with an ancestral *N_e_* of 1,225,393 and current *N_e_* of 1,357,760. Our estimates of ancestral *N_e_* are comparable to those inferred by previous studies of African populations of *D. melanogaster* (Li and Stephan 2006; Laurent *et al.* 2011; Duchen *et al.* 2013; Arguello *et al.* 2019; Figure 6). As shown in Figure 6, we infer a much larger proportion of mildly deleterious mutations and a smaller proportion of highly deleterious mutations than in previous studies (Keightley and Eyre-Walker 2007; Huber *et al.* 2017), with *f*_0_ = 24.7%, *f*_1_ = 49.4%, *f*_2_ = 3.9%, and *f*_3_ = 21.9%, but this reflects the fact that our procedure includes synonymous sites among the total. Because we have inferred the DFE for a select class of single exon genes, which have slightly higher than average divergence (Figure S15), it is possible that these exons are experiencing weaker purifying selection than the genome-wide mean. Furthermore, because we have obtained the DFE of both coding sequences and UTR regions, 4-fold degenerate and UTR sites comprise 12% and 29% of the total, respectively. Previous studies have estimated 6-10% of all mutations at non-synonymous sites to be effectively neutral. Thus, assuming that all 4-fold degenerate sites are neutral, ∼40% of UTR regions are neutral (Andolfatto 2005; Campos *et al.* 2017), and ∼6-10% of nonsynonymous mutations are neutral, we expect *f*_0_ to be ∼27-30%. Encouragingly, we infer *f*_1_ =25%. This observation implies that the majority of synonymous sites are not experiencing direct selection, consistent with previous results for *D. melanogaster* (Jackson *et al.* 2017). Further, although we infer a larger proportion of weakly deleterious mutations than previous studies from the distribution of γ =2*N*_e_*s*, the distribution of *s* is quite comparable (Table S9) to that inferred by Huber *et al*. (2017).

**Figure 6:**
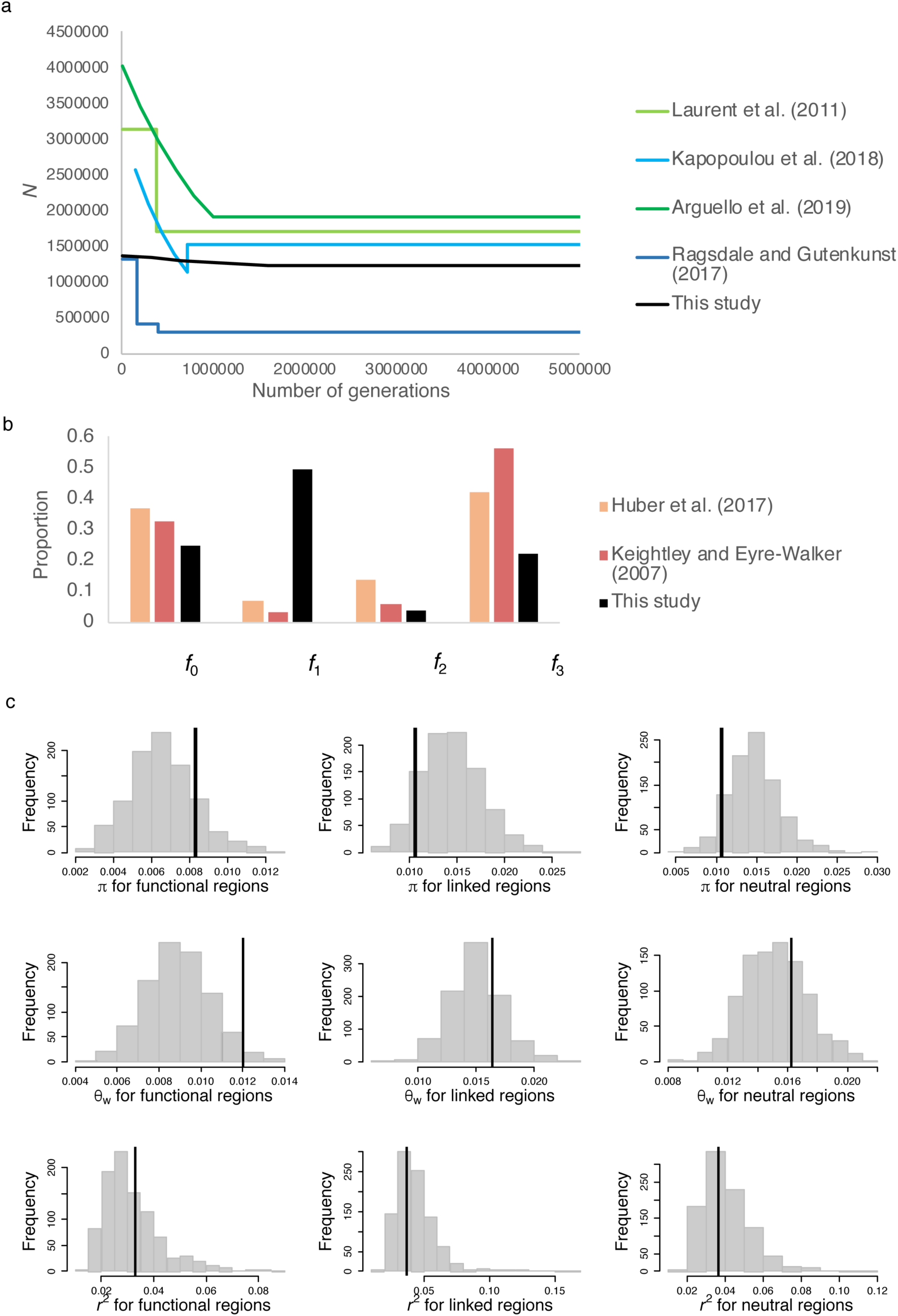
Joint inference of demography and purifying selection in the Zambian population of *D. melanogaster*. (a) Demographic model inferred in previous studies of the Zambian population (blue lines), the Zimbabwe population (green lines), and the current study (black lines). (b) The DFE for deleterious mutations in coding regions (including synonymous and non-synonymous sites) as inferred by previous studies of other populations (colored bars) and at exonic sites of single-exon genes as inferred in the current study (black bars). The X-axis is for *f*_0_: 0≤ 2*N*_e_*s* <1, *f*_1_: 1≤ 2*N*_e_*s* <10, *f*_2_: 10≤ 2*N*_e_*s* <100, and *f*_3_: 100≤ 2*N*_e_*s* <10000. For the previous studies, the DFE shown in this figure includes the fraction of synonymous sites in the neutral *f*_0_ class. (c) Distribution of key summary statistics (*π*, *θ*_W_, *r*^2^) for functional, linked and neutral regions when simulating 100 replicates of 94 exons each using the inferred parameters. The vertical lines represent values of the statistics obtained from 76 individuals of *D. melanogaster* from Zambia, after excluding non-coding sites with phastCons score ≥ 0.8.

In order to test whether our inferred parameters explain the observed *D. melanogaster* data, we simulated 10 replicates of each of the 94 exons using the parameter estimates, and evaluated whether the means of the observed *D. melanogaster* values are in the 5% tails of the simulated distribution of statistics. Our parameter estimates result in a very good fit to empirical *D. melanogaster* population data (Figure 6, Figure S16) for all three categories – functional (*i.e*., exonic), linked (*i.e*., non-coding region adjacent to exons) and neutral (*i.e.,* non-coding region adjacent to the linked region), except for Tajima’s *D* (linked region *p* = 0.011, neutral region *p* = 0.010) and divergence (linked region *p* = 0.029, neutral region *p*=0.0) in intergenic regions – although these statistics are well fitted in functional regions.

As both positive selection in exons and purifying selection in non-coding regions could partially drive these patterns, we investigated both of these model violations. Non-coding regions flanking 2kb of the selected exons (which were used to perform inference) were found to have 777 sites that had phastCons scores greater than or equal to 0.8, with a mean and median length of 25 and 15 bp, respectively. We therefore simulated conserved elements in non-coding regions that were 20bp in length, uniformly distributed, and which made up 40% of the flanking neutral sites (*i.e*., 800 sites in total). Conserved elements were simulated with either weak (*f*_1_ =1), moderate (*f*_2_ =1) and strong (*f*_3_ =1) purifying selection. Upon masking these sites, as was done in our *Drosophila* data analysis, there was no observed difference in the distribution of all statistics (Figure S17), suggesting that background selection caused by small conserved elements does not significantly affect our inference, and in fact does not alter the fit of our inferred model to the data. Interestingly, without masking sites – that is, by allowing sites that experience direct weak purifying selection to remain in the flanking sequences – our model is much better able to explain the lower Tajima’s *D* and divergence in intergenic regions (Figure S18). It thus appears likely that weak purifying selection on sequences in intergenic regions could contribute to the discrepancies between observed and expected.

Next, we simulated positive selection with selection coefficient *s_b_* for beneficial mutations under 4 different scenarios - representing rare and strong (1% of all mutations in exonic regions are beneficial with 2*N_e_s_b_*= 1000), common and strong (5% of mutations in exonic regions are beneficial with 2*N_e_s_b_* = 1000), common and weak (5% of mutations in exonic regions are beneficial with 2*N_e_s_b_* = 10) and rare and weak (1% of mutations in exonic regions are beneficial with 2*N_e_s_b_* = 10) selection. We find that, although strong positive selection, whether common or rare, better explains the lower Tajima’s *D* values in intergenic regions, it also drastically alters the distribution of most other statistics, resulting in an overall much poorer fit (Figure S19, S20). For instance, common and strong positive selection reduces *θ*_H_ by an order of magnitude relative to our fitted model, and drastically increases the variance while decreasing the mean of haplotype diversity. In contrast to strong positive selection, weakly positively selected mutations do not alter the distribution of Tajima’s *D* in intergenic regions, but slightly increase *θ*_H_ in functional regions, which improves the fit to the observed data (Figure S21, S22). In addition, all cases of positive selection significantly increase divergence in functional regions. For comparison, we also simulated the two scenarios of positive selection used by Lange and Pool (2018) - 0.2% of all mutations are beneficial with 2*N*_e_*s_b_* =60, and 0.00013% of all mutations are beneficial with 2*N*_e_*s_b_* =10000. As the frequency of positively selected alleles is lower in these scenarios, there was no observed difference between the distribution of statistics resulting from including or excluding positive selection (Figure S23, S24). Thus, if the frequency of strongly positively selected mutations is much lower than 1%, as was proposed by Campos *et al.* (2017), our estimates of both demography and DFE shape should be unbiased, and beneficial fixations would be virtually undetectable. Future studies will investigate the ability of our approach to quantify the properties of beneficial mutations.

## CONCLUSION

Independent of specific views on the roles of adaptive vs. non-adaptive explanations for observed levels and patterns of DNA sequence variation and divergence, it has been widely accepted that natural populations are not at demographic equilibrium, but are often characterized by fluctuating population sizes and other demographic perturbations. Additionally, a rich empirical and experimental literature has demonstrated the pervasive importance of purifying selection in eliminating the constant input of deleterious variants. It has also been found that ignoring direct effects of purifying selection and its impact on linked sites can strongly bias demographic inference (Ewing and Jensen 2016), and that ignoring demographic effects biases estimates of parameters of selection (Jensen *et al.* 2005; Thornton and Jensen 2007; Crisci *et al.* 2012, 2013). Yet, despite agreement that these processes are certain to be occurring constantly in populations and shaping patterns of variation and evolution, the construction of a statistical approach capable of simultaneously estimating parameters of the concerned processes has been difficult. Here we provide one such approach, for which we demonstrate an ability to co-estimate the parameters of a generalized DFE along with those underlying the population history.

By fitting a four-parameter DFE model that includes weak, intermediate and strong purifying selection, as well as neutrally evolving sites, this approach avoids two common, and potentially perilous, assumptions: 1) synonymous sites are not assumed to be neutral, consistent with a growing body of literature (Chamary and Hurst 2005; Lynch 2007; Zeng and Charlesworth 2010a; Lawrie *et al.* 2013; Choi and Aquadro 2016; Jackson *et al.* 2017), and 2) the DFE is not assumed to follow a specific parameterized distribution, such as the widely-used gamma distribution.

Our results demonstrate that it is possible to jointly infer the deleterious DFE and past demographic changes using an ABC framework, by including various summary statistics capturing aspects of the SFS, linkage disequilibrium and divergence, compared between coding and flanking non-coding sequences. Ancestral population sizes and the frequency of the most deleterious classes of the DFE are estimated with relatively low accuracy, whereas the current population sizes and the neutral mutation class are estimated with high accuracy. In addition, we demonstrated that, if synonymous sites are indeed experiencing substantial purifying selection, existing programs such as DFE-alpha will over-estimate recent growth and under-estimate the proportion of mildly deleterious mutations. Importantly, the approach proposed here performs equally well regardless of whether synonymous sites are neutral or selected. However, our approach continues to assume the neutrality of flanking non-coding regions, though putatively conserved sites were masked; the impact of this masking on inference was thoroughly assessed via simulation.

Because we make no assumptions about which sites in the functional region of interest are neutral, it is in principle possible to estimate the DFE for any functional element using this methodology. The results further suggest that the accurate co-estimation of these parameters is possible using only functional regions. Such an approach may be extremely useful in genomes for which it is difficult to characterize putatively neutral sites, as well as for compact genomes in which non-coding regions may be limited. However, we have only tested relatively simple demographic models, and future studies evaluating our ability to jointly estimate more complex population histories would be of value.

This approach can in principle be applied to any organism and functional class of interest, although power analyses suggest the utility of prior knowledge of the boundaries of functional regions as well as mutation and recombination rates. Here we have provided an illustrative example with *D. melanogaster*. The results suggest that the Zambian population of this species has been largely stable in size, and that exonic regions have a large proportion of mildly deleterious mutations. Although this result might seem surprising, the DFE inferred by the current method provides the distribution of selective effects over all sites, including synonymous sites and sites in UTRs. Hence, in comparing the DFE estimated in the current study with previous estimates of the neutral class of mutations, it appears unnecessary to invoke widespread selection on synonymous sites in *D. melanogaster.* This result is consistent with most previous studies (Akashi 1995; Jackson *et al.* 2017), and our estimate of the strength of purifying selection acting on synonymous sites in the Zambian population is in line with earlier estimates for African populations (Zeng and Charlesworth 2010a; Jackson *et al.* 2017).

In addition to the proposed inference framework, we have derived an analytical expression for the reduction in variation caused by background selection at neutral sites outside functional regions for the case of a discrete DFE, making it feasible to obtain analytical predictions for any chosen DFE. Not only does a discrete DFE provide flexibility in inference, it may also be a more realistic representation of the true DFE (Kousathanas and Keightley 2013; Bank *et al.* 2014b). Although gamma distributions represent a reasonably good fit to the DFE inferred from genome-wide studies (Eyre-Walker and Keightley 2007), the DFE will be mis-inferred if the true distribution is multimodal (Kousathanas and Keightley 2013), as has been widely observed (*e.g*., in yeast (Bank *et al.* 2014a), viruses (Sanjuán 2010), and *E.coli* (Jacquier *et al.* 2013)). In addition, the best-fitting parameterized continuous distribution appears to be extremely specific to the particular dataset being tested, and most alternative distributions fit the data nearly as well as the best-fitting distribution (Huber *et al.* 2017; Kim *et al.* 2017). The discrete DFE proposed here thus reduces the number of necessary assumptions, and has been shown to perform well in the plausible scenario in which common assumptions are indeed violated (*e.g*., if the true DFE is not gamma-distributed). Both the analytical results under demographic equilibrium and simulations under demographic non-equilibrium show that the number of selected sites and the specific shape of the DFE (for instance, the frequency of mildly and moderately deleterious mutations) both decrease linked neutral variation around functional regions more than previously appreciated, and skew the SFS even when there is no reduction in diversity. Such variation in exon lengths and DFE shapes across a genome can increase the variance of statistics in linked neutral regions, which could contribute to false positives when detecting positive selection using outlier approaches.

There are at least three important caveats, which will be the subject of future study. The first concerns the estimates of ancestral and current effective population sizes. As the effective population size varies across the genome in a fashion correlated with local recombination rates (Becher *et al.* 2020), the estimates provided here ought to be viewed as a mean across the loci in question. While we have improved upon the common assumption of a singular genome-wide value by directly modeling each locus-specific recombination rate when performing inferences, the general importance of this effect in demographic modeling remains in need of further study. The second caveat concerns biases in inference using ABC-based methods under model violations, such as a mis-specification of the mutation or recombination rate. As the method is sensitive to such violations, it will be best applied to organisms in which these parameters have been experimentally measured. Moreover, we have not included other types of mutations such as insertions / deletions, gene duplications and TE insertions, all of which will increase the deleterious mutation rate and thus the effects of BGS (Comeron 2014).

The third caveat concerns inferences about selection. This study represents a proof-of-concept in demonstrating that the simultaneous inference of demography and the DFE is feasible, thereby avoiding common assumptions underlying a step-wise inference approach. While this interplay of genetic drift and purifying selection is in fact alone sufficient to fit all features of the data (consistent with previous claims: Comeron 2014, 2017; Harris *et al.* 2018; Jensen *et al.* 2019), this is not the same as claiming that positive selection is not also occurring. As our simulation results demonstrate, the presence of rare, weakly beneficial mutations is consistent with the data, though the inclusion of these parameters does not result in an improved fit. The question is less about presence/absence, than it is about statistical identifiability. Conversely, the addition of a strongly beneficial mutational class was found to be inconsistent with observed data. In order to investigate this further, future work will evaluate the ability to co-estimate a beneficial class of fitness effects within this framework. It should also be noted that the example chosen to highlight our approach focuses on only a subset of genes in the *D. melanogaster* genome, and the observed DFE in this class is not necessarily universal across all coding regions in the population under consideration. In fact, the means of the scaled selection coefficients of deleterious mutations have been shown to be negatively correlated with divergence at nonsynonymous sites (Campos *et al.* 2017). Importantly, however, a general inference approach that incorporates these two dominant processes will be a valuable tool in future genomic scans, and our appropriate null is anticipated to greatly reduce the notoriously high false-positive rates associated with the identification of positively selected loci. The ability of our approach to reject the hypothesis of frequent hard selective sweeps involving strongly selected beneficial mutations is encouraging, as hypothesis rejection is often scientifically more robust than model fitting.

## ACKNOWLEDGEMENTS

We would like to thank Rebecca Harris for discussions related to this project. This research was conducted using resources provided by Research Computing at Arizona State University (http://www.researchcomputing.asu.edu) and the Open Science Grid, which is supported by the National Science Foundation and the U.S. Department of Energy’s Office of Science. We especially thank Lauren Michael and Christina Koch from the Open Science Grid for their efforts to provide technical assistance. This work was funded by National Institutes of Health grant R01GM135899 to JDJ.

## APPENDIX

## Derivation of the analytical expression for the reduction in diversity due to background selection

We assume a discrete DFE with four bins, such that *t* is uniformly distributed within each bin. The expectation of *E*(*t*) within each bin is proportional to its integral with respect to *t*. The overall expectation of *E*(*t*) is given by:

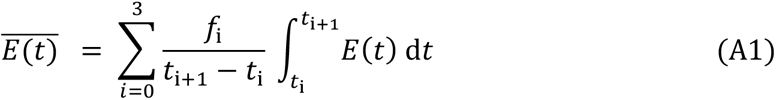

where the *t*_i_ correspond to the boundaries of the discrete bins. These are such that 0 ≤ 2*N_e_s* ≤ 1, 1 ≤ 2*N_e_s* ≤ 10, 10 ≤ 2*N_e_s* ≤ 100 and 100 ≤ 2*N_e_s* ≤ 10000, respectively. In our case, these correspond to *t*_0_ = 0, *t*_1_ = 0.00005, *t*_2_ = 0.0005, *t*_3_= 0.005, and *t*_4_ = 0.5. While this mirrors the DFE considered here, a similar procedure can be used for any set of bins for a given DFE.

In order to determine *E*(*t*) from the first line of Equation 2 of the main text, we write *a* = *g* + *r_c_y* and *b* = *g* + *r_c_*(*y* + *l*), and note that:

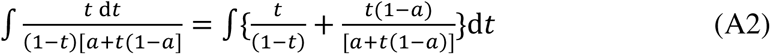

The second integral on the right-hand side of this equation can be evaluated by substituting *u* = *a* + *t*(1 – *a*) for *t*, so that d*t* = d*u*/(1 – *a*). This gives:

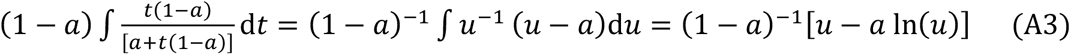

With this change in variable, the normalizing factor for the probability density function is now (*u*_1_ – *u*_0_)^−1^= (1 – *a*) ^−1^(*t*_1_ – *t*_0_)^−1^. The contribution of this component to the expectation of *E*(*t*) yields Equation 3a of the main text.

A similar expression can be written for the integral of – *t*/[(1 – *t*)[*b* + *t*(1– *b*)] in the first line of Equation 2. When adding this to the integral of *t*/[(1 – *t*)[*a* + *t*(1– *a*)], the integrals involving 1/(1 – *t*) cancel out, so that this term simply contributes Equation 3b to the expectation of *E*(*t*).

## SUPPLEMENTARY FIGURES AND TABLES

**Table S1:** Predictions of diversity in linked neutral regions for two different DFE realizations, as predicted by Equations 3a and 3b. The expected diversity with no selection is 0.02.

| DFE | Length of coding region: | Distance from selected site --> |  |  |
| --- | --- | --- | --- | --- |
|  |  | 100 bp | 10 kb | 100 kb |
| $f_0 = 0; f_1 = 0, f_2 = 50, f_3 = 50$ | 1 kb | 0.0174857 | 0.019805 | 0.0199807 |
|  | 5 kb | 0.0150363 | 0.0192131 | 0.0199054 |
|  | 10 kb | 0.0141392 | 0.0187112 | 0.0198152 |
| $f_0 = 0; f_1 = 0; f_2 = 80; f_3 = 20$ | 1 kb | 0.0160713 | 0.0197472 | 0.0199892 |
|  | 5 kb | 0.0124679 | 0.0190189 | 0.0199474 |
|  | 10 kb | 0.0113456 | 0.0184565 | 0.0198979 |

**Table S2:** Reduction in neutral and linked neutral diversity calculated analytically as a function of the DFE - as illustrated by considering DFE realizations in which one class is largely over-represented, for different exon lengths (0.5kb, 1kb, 5kb, and 10kb). The expected diversity under neutrality is 0.02.

| | $f_0 \geq 80\%$ | $f_1 \geq 80\%$ | $f_2 \geq 80\%$ | $f_3 \geq 80\%$ |
| --- | --- | --- | --- | --- |
| **Neutral Diversity (0.5 kb)** | 0.01995 | 0.01982 | 0.01958 | 0.01975 |
| **Linked neutral diversity (0.5kb)** | 0.01976 | 0.01809 | 0.01795 | 0.01891 |
| **Slope of recovery (0.5kb)** | 0.00007 | 0.00054 | 0.00063 | 0.00013 |
| **Neutral Diversity (1 kb)** | 0.01992 | 0.01972 | 0.01926 | 0.01951 |
| **Linked neutral diversity (1kb)** | 0.01968 | 0.01756 | 0.01674 | 0.01906 |
| **Slope of recovery (1kb)** | 0.00010 | 0.00072 | 0.00100 | 0.00024 |
| **Neutral Diversity (5 kb)** | 0.01972 | 0.01934 | 0.01760 | 0.01814 |
| **Linked neutral diversity (5 kb)** | 0.01915 | 0.01634 | 0.01318 | 0.01714 |
| **Slope of recovery (5kb)** | 0.00024 | 0.00109 | 0.00200 | 0.00061 |
| **Neutral Diversity (10kb)** | 0.01960 | 0.01909 | 0.01673 | 0.01709 |
| **Linked neutral diversity (10kb)** | 0.01903 | 0.01615 | 0.01208 | 0.01579 |
| **Slope of recovery (10kb)** | 0.00027 | 0.00112 | 0.00220 | 0.00083 |

**Table S3:**
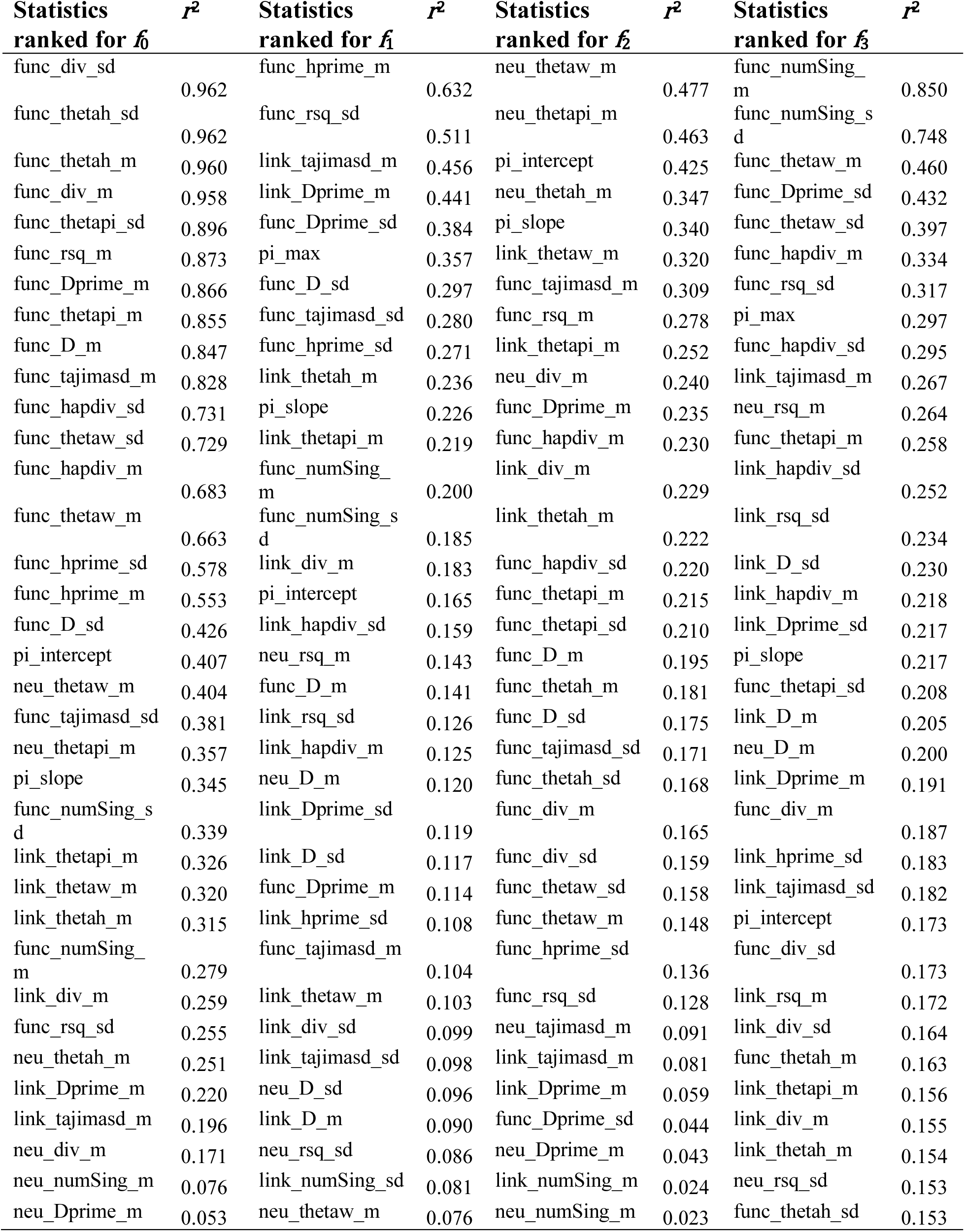
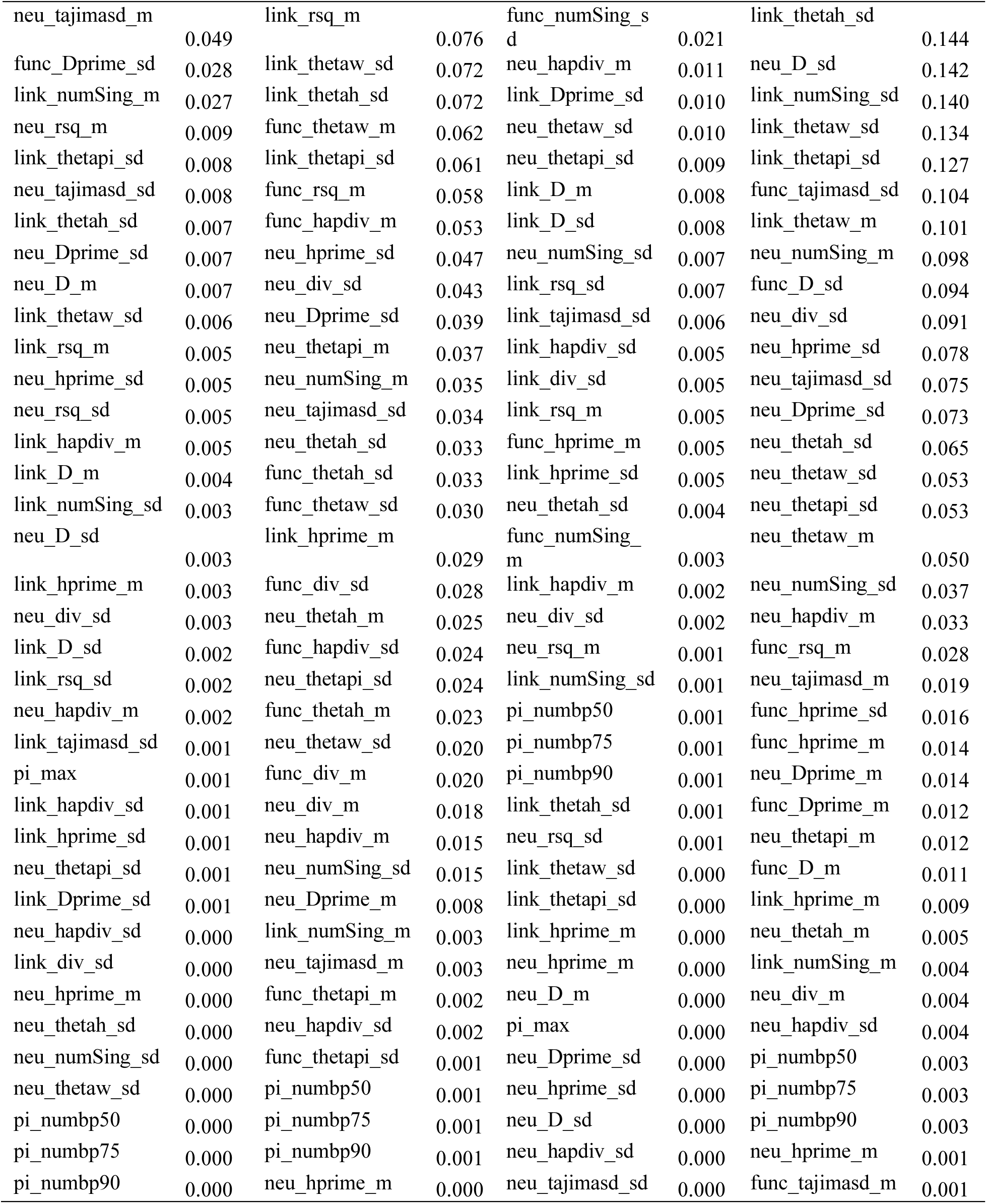
Statistics ranked by their importance in predicting the DFE classes under equilibrium using the correlation coefficients between the statistics and parameters.

**Table S4:**
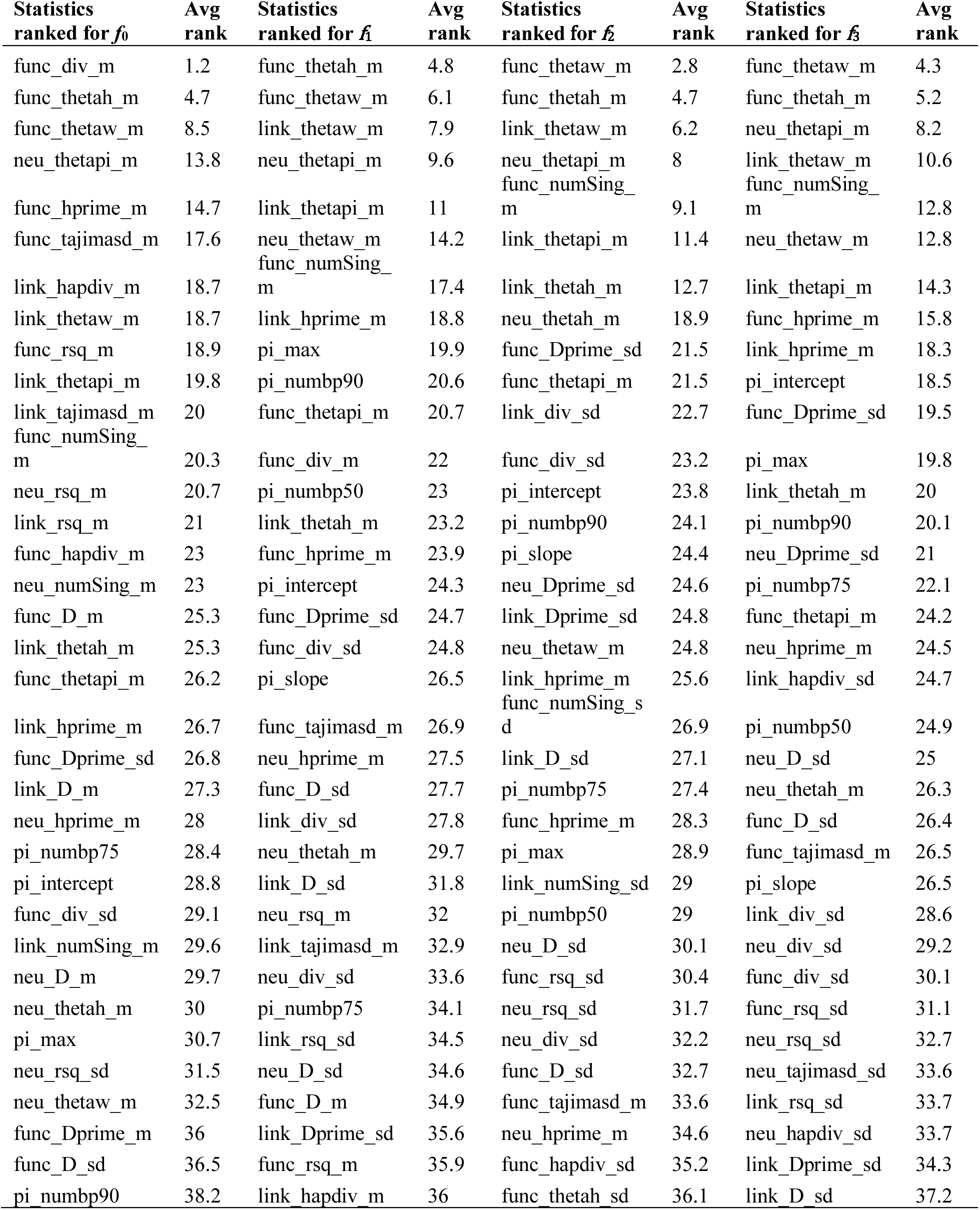
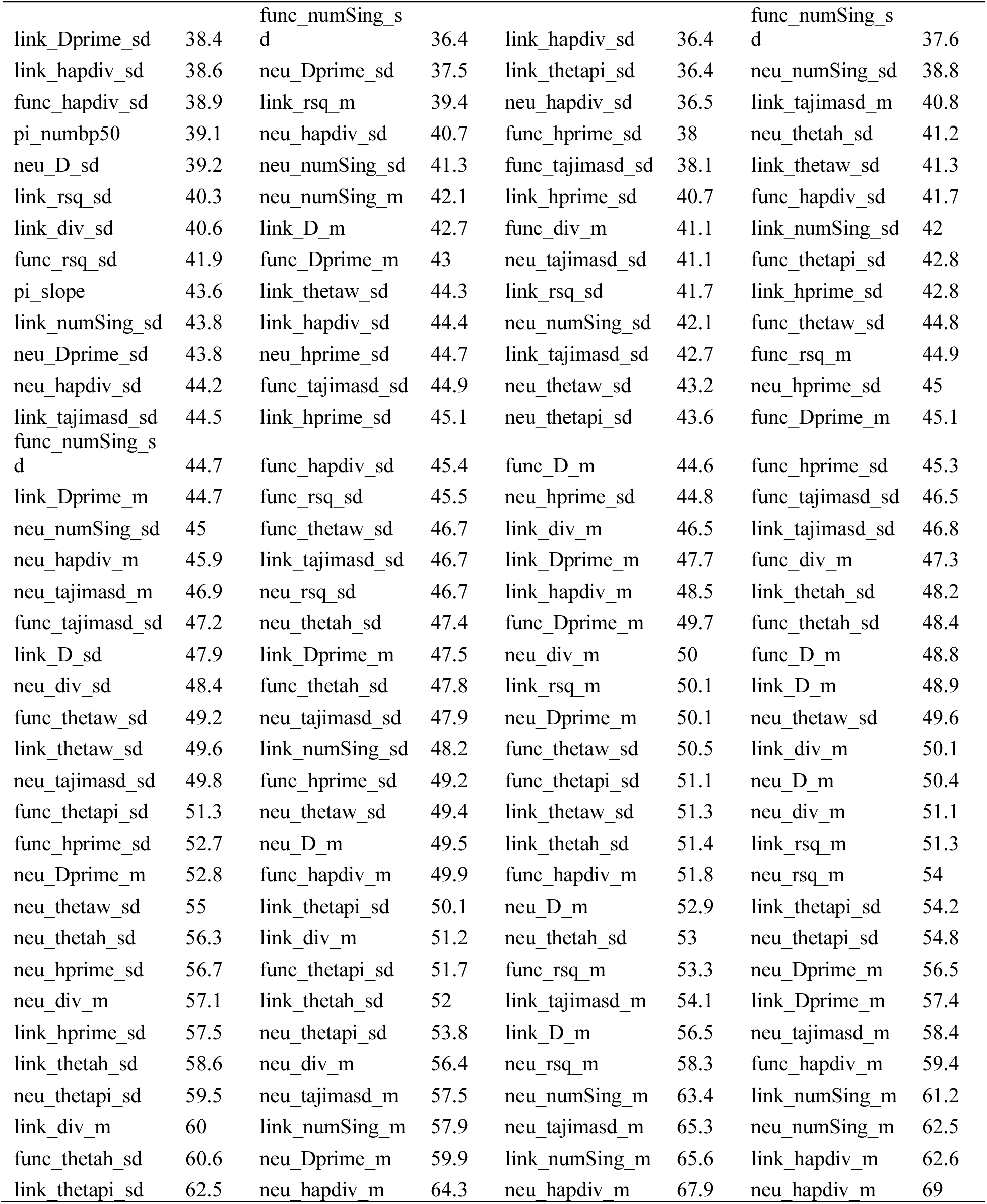
Statistics ranked by their importance in predicting the DFE classes under equilibrium using a modified algorithm of Joyce and Marjoram (2008) and by averaging the ranking across 10 replicates for each parameter separately.

**Table S5:**
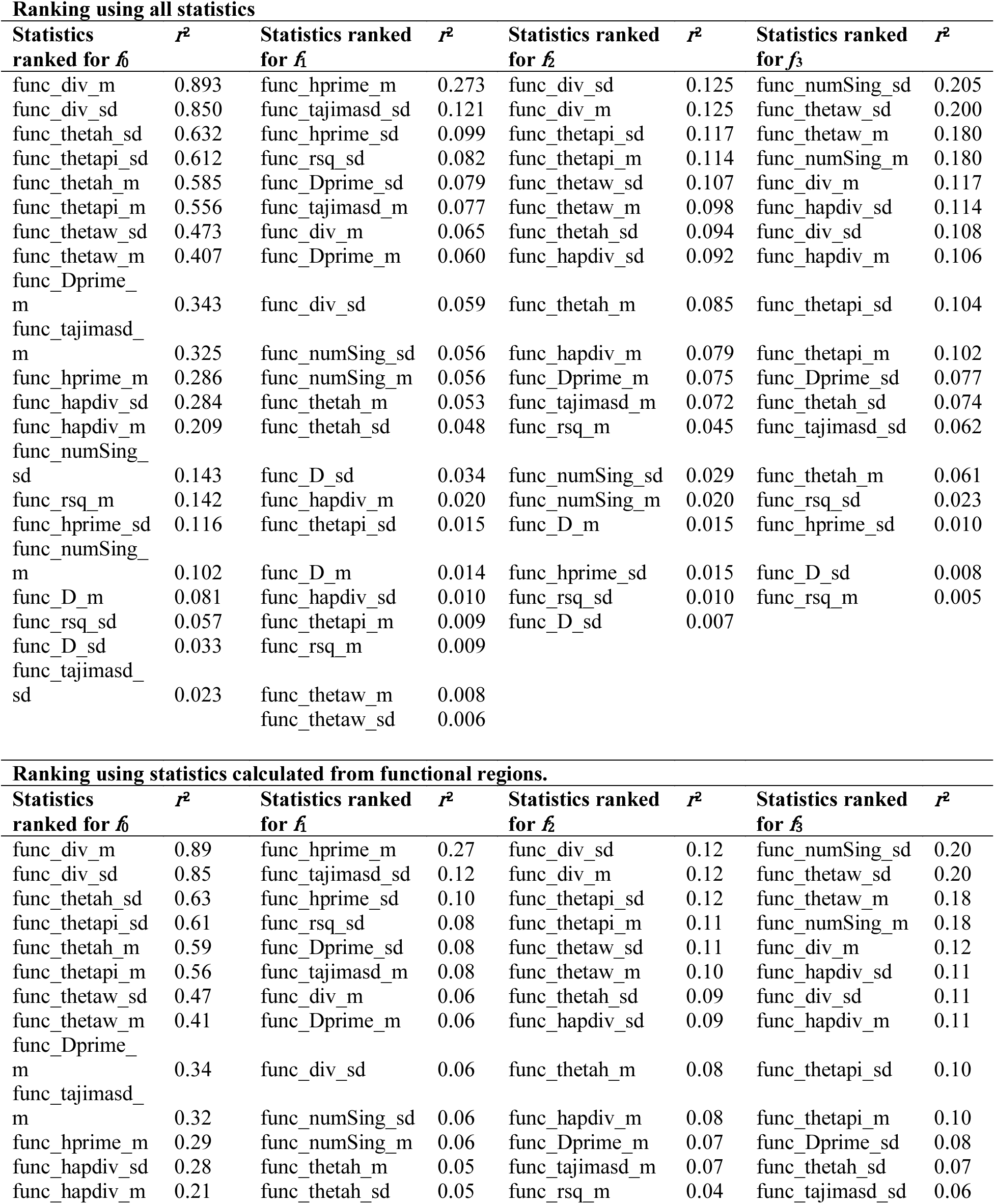
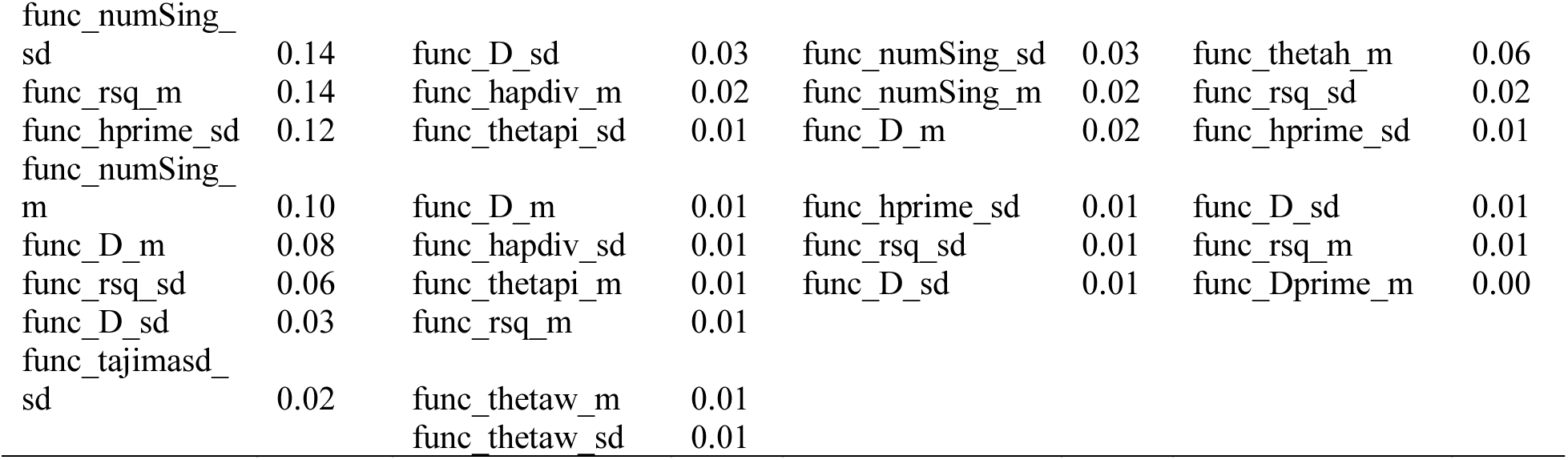
Ranking of statistics under demographic non-equilibrium. Statistics significantly correlated with parameters of the DFE when statistics from all regions are used and when only functional statistics are used for ranking. Significance was evaluated with *p* < 0.05 with Bonferonni corrections.

**Table S6:**
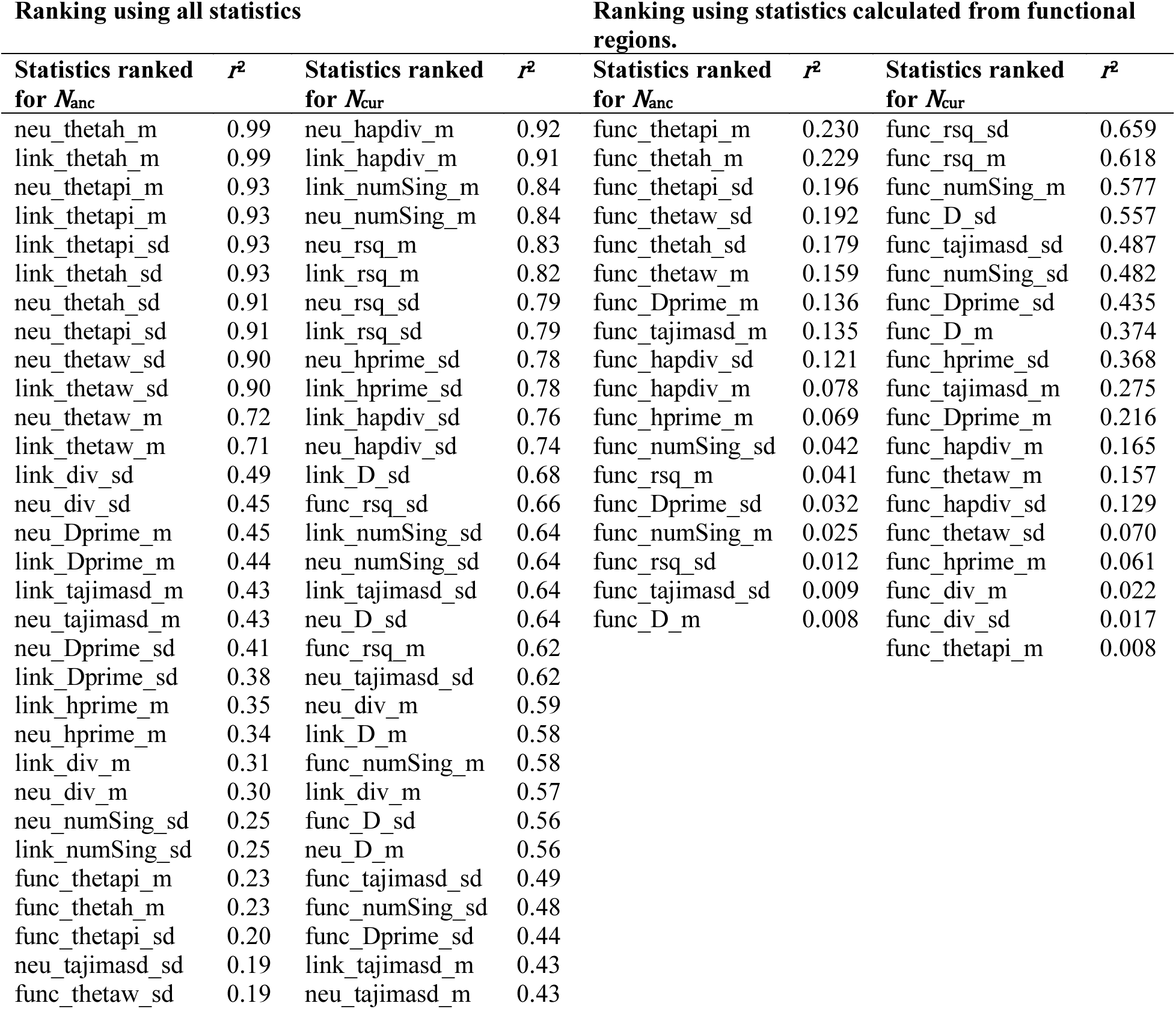
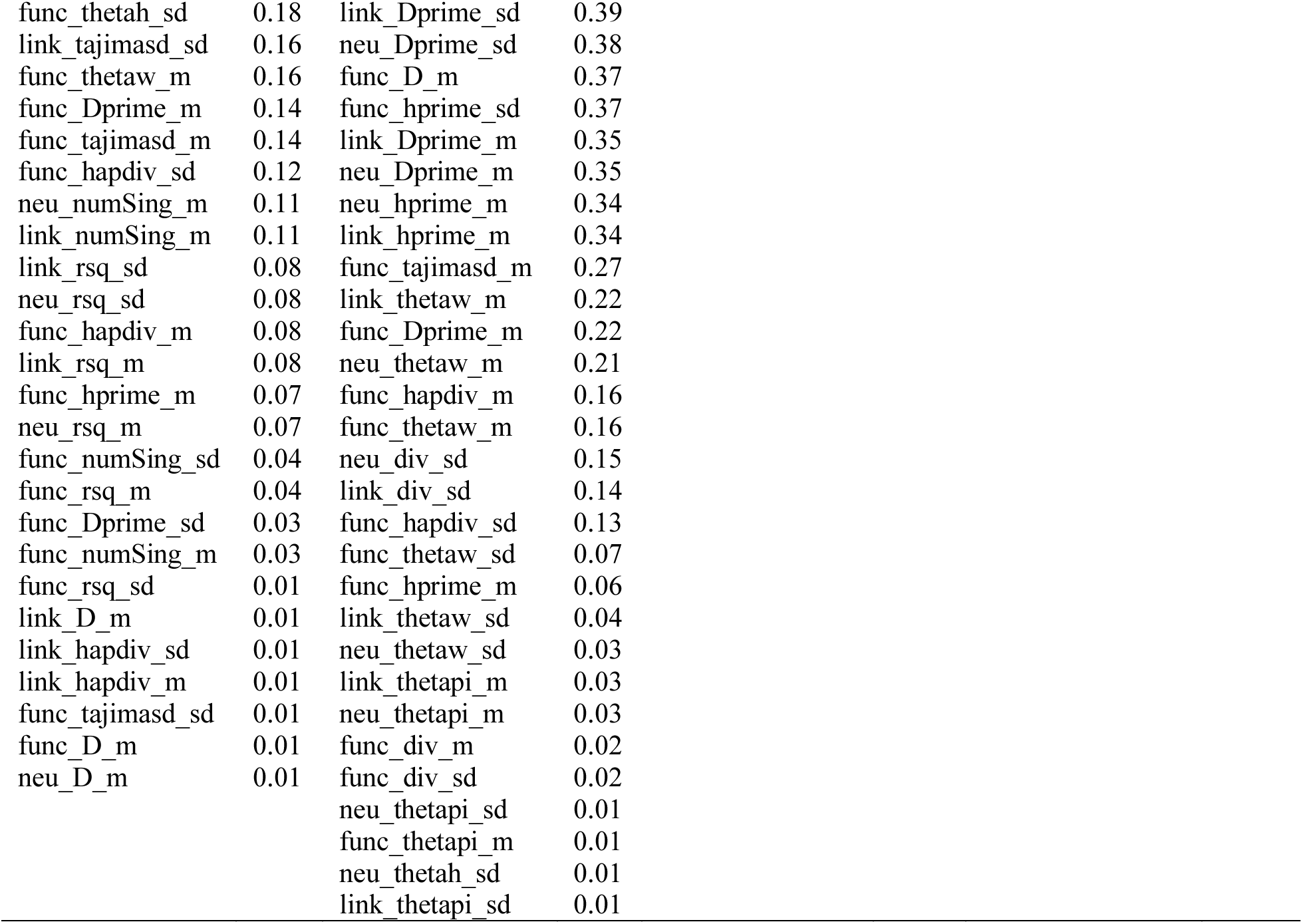
Ranking of statistics when distinguishing between demography and purifying selection. Statistics significantly correlated with parameters of demography when statistics from all regions are used, and when only functional statistics are used for ranking. Significance was evaluated with *p* < 0.05 with Bonferonni correction.

**Table S7:** The mean numbers of fixed differences per site (*i.e*., polymorphism-adjusted divergence) for different site types in *D. melanogaster*, where different numbers of individuals from the Zambia population were used to identify the set of polymorphic sites.

|  | **Sample size:** |  |  |  |  |  |
| --- | --- | --- | --- | --- | --- | --- |
|  | **1** | **2** | **5** | **15** | **30** | **76** |
| **exon** | 0.0238 | 0.0198 | 0.0170 | 0.0160 | 0.0159 | 0.0153 |
| **coding** | 0.0228 | 0.0182 | 0.0157 | 0.0146 | 0.0141 | 0.0135 |
| **4-fold** |  |  |  |  |  |  |
| **degenerate** | 0.0497 | 0.0423 | 0.0349 | 0.0316 | 0.0311 | 0.0300 |
| **0-fold** |  |  |  |  |  |  |
| **degenerate** | 0.0182 | 0.0123 | 0.0108 | 0.0102 | 0.0098 | 0.0094 |

**Table S8:**
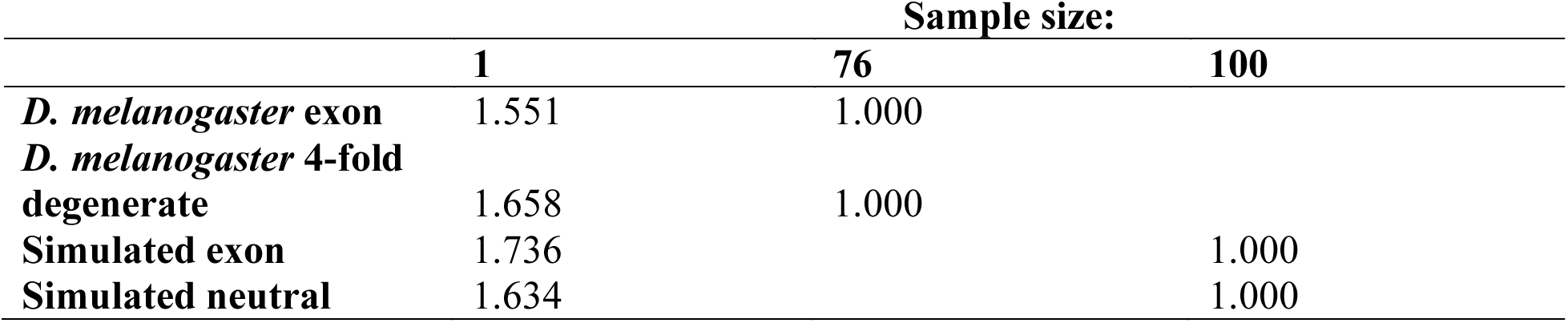
The increase in divergence values obtained when calculating pairwise divergence (corresponding to a sample size of 1) relative to alternate sample sizes (which exclude polymorphic sites from divergence).

|  | **Sample size:** |  |  |
| --- | --- | --- | --- |
|  | **1** | **76** | **100** |
| ***D. melanogaster* exon** | 1.551 | 1.000 |  |
| ***D. melanogaster* 4-fold degenerate** | 1.658 | 1.000 |  |
| **Simulated exon** | 1.736 |  | 1.000 |
| **Simulated neutral** | 1.634 |  | 1.000 |

**Table S9:** Inference of the DFE in 94 exons of *D. melanogaster*. Our inference is only comparable to that of Huber *et al*. (2017) for two classes of *s* – less than and greater than 10^-4^.

| **\|s\|** | **0-10<sup>-4</sup>** | **≥10<sup>-4</sup>** |
| --- | --- | --- |
| **Huber *et al.* 2017 - Gamma distribution** | 0.767 | 0.233 |
| **Huber *et al.* 2017 - Lognormal distribution** | 0.667 | 0.333 |
| **This study** | 0.781 | 0.219 |

**Figure S1:**
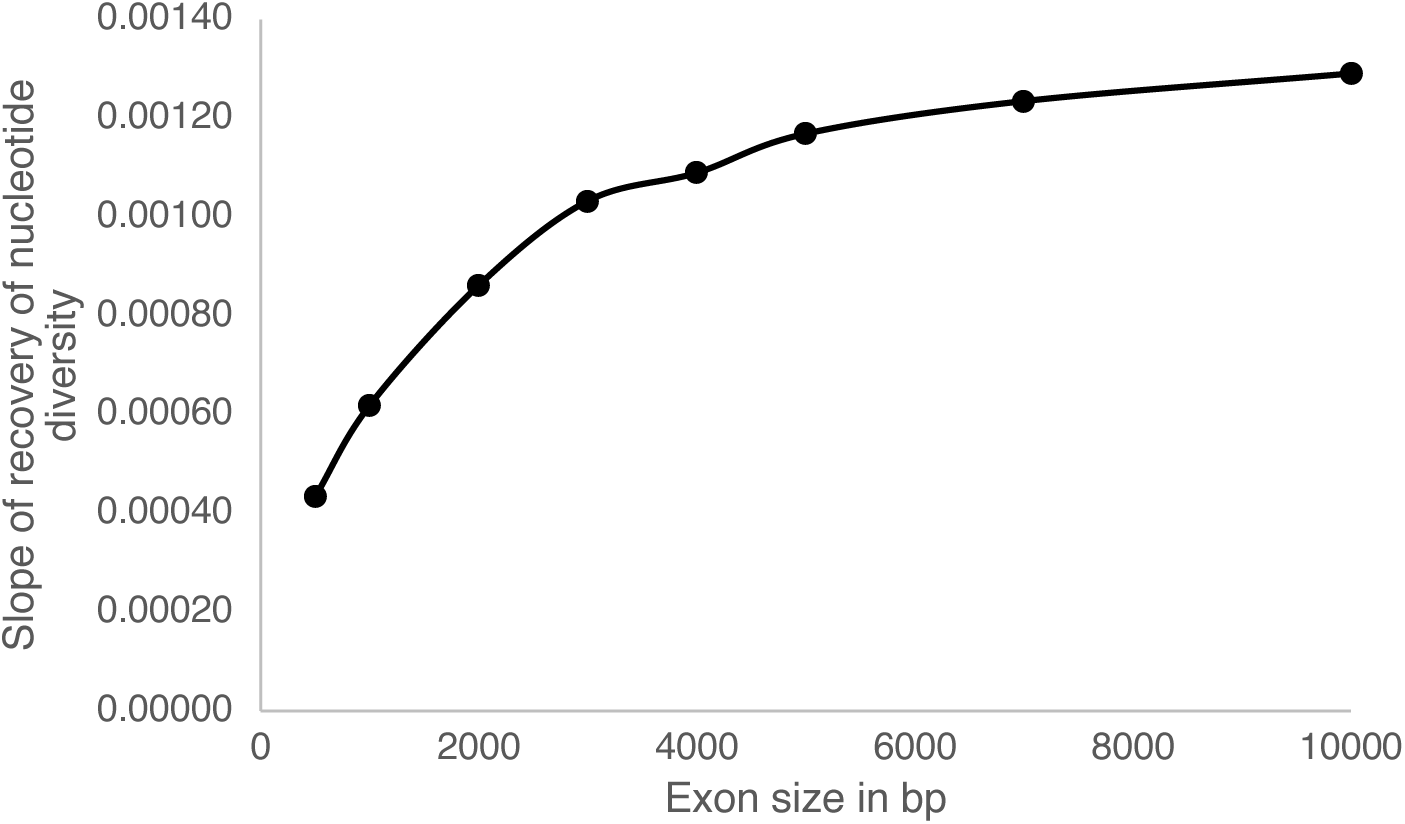
Increase in the slope of recovery of diversity near functional regions of varying sizes as observed via simulations. Larger values of the slope represent a steeper recovery, concordant with larger reduction in diversity observed in the non-coding region.

**Figure S2:**
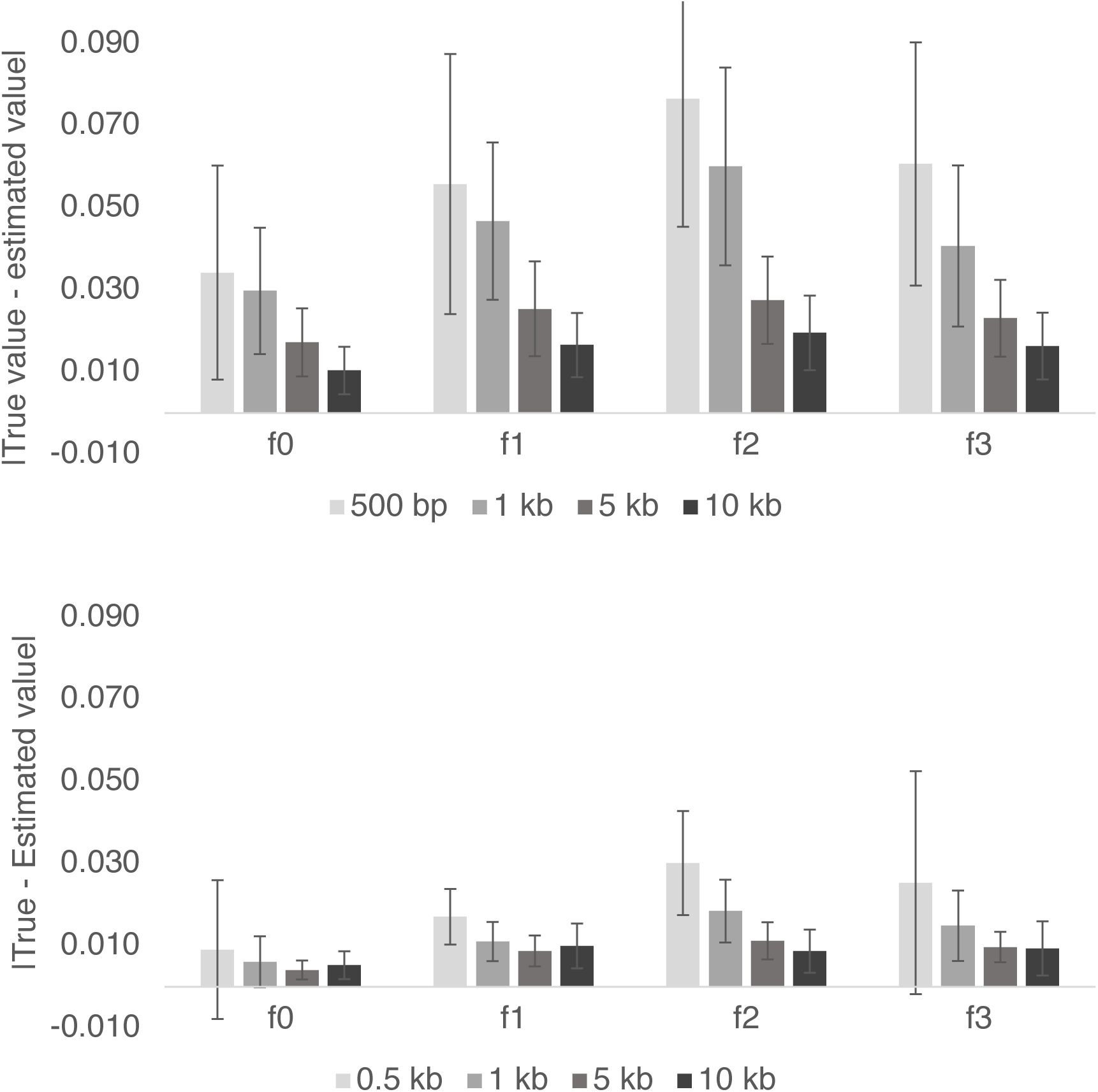
Absolute difference between true and inferred value of parameters characterizing the DFE for lengths of 0.5 kb, 1 kb, 5 kb, and 10 kb of functional regions. The upper panel displays the error in inference when using all statistics, while the lower uses only functional regions.

**Figure S3:**
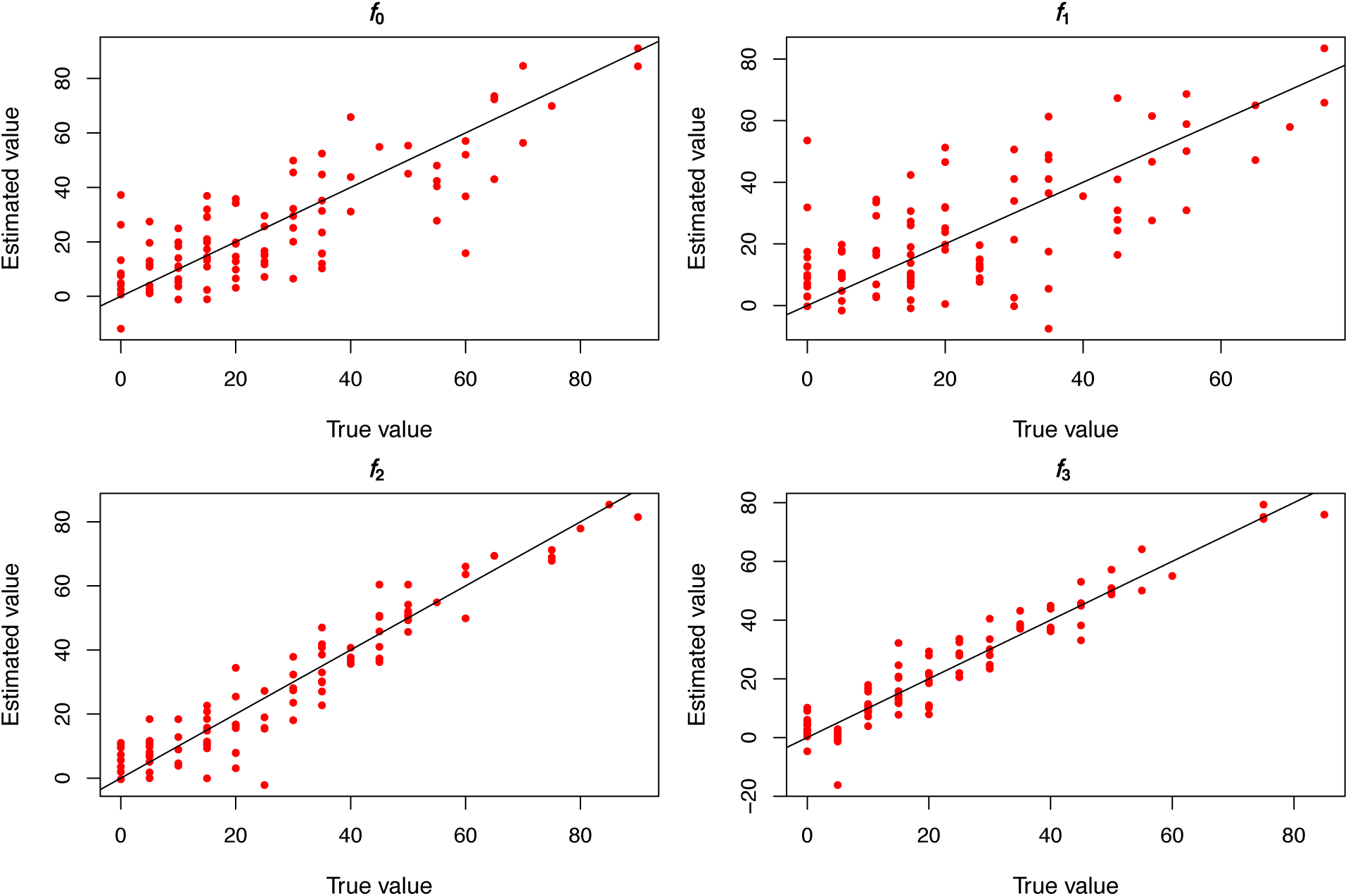
Inference of the DFE under demographic equilibrium using only statistics in the linked neutral regions. The length of the functional region is 10kb.

**Figure S4:**
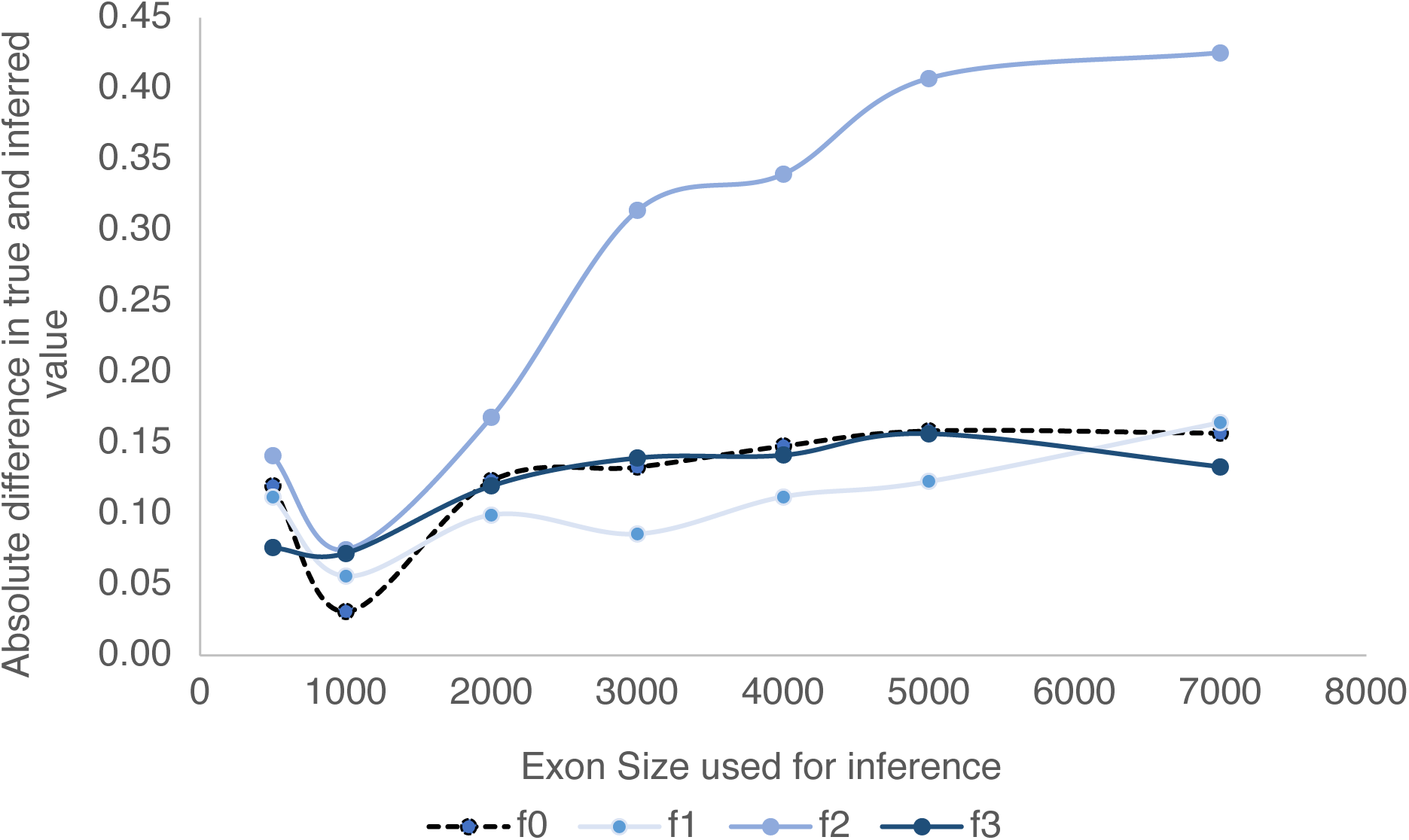
Decrease in accuracy of inference for different DFE classes as the exon size assumed for inference is mis-specified. In this figure, the assumed exon size was 1kb, and the X-axis gives the true exon size.

**Figure S5:**
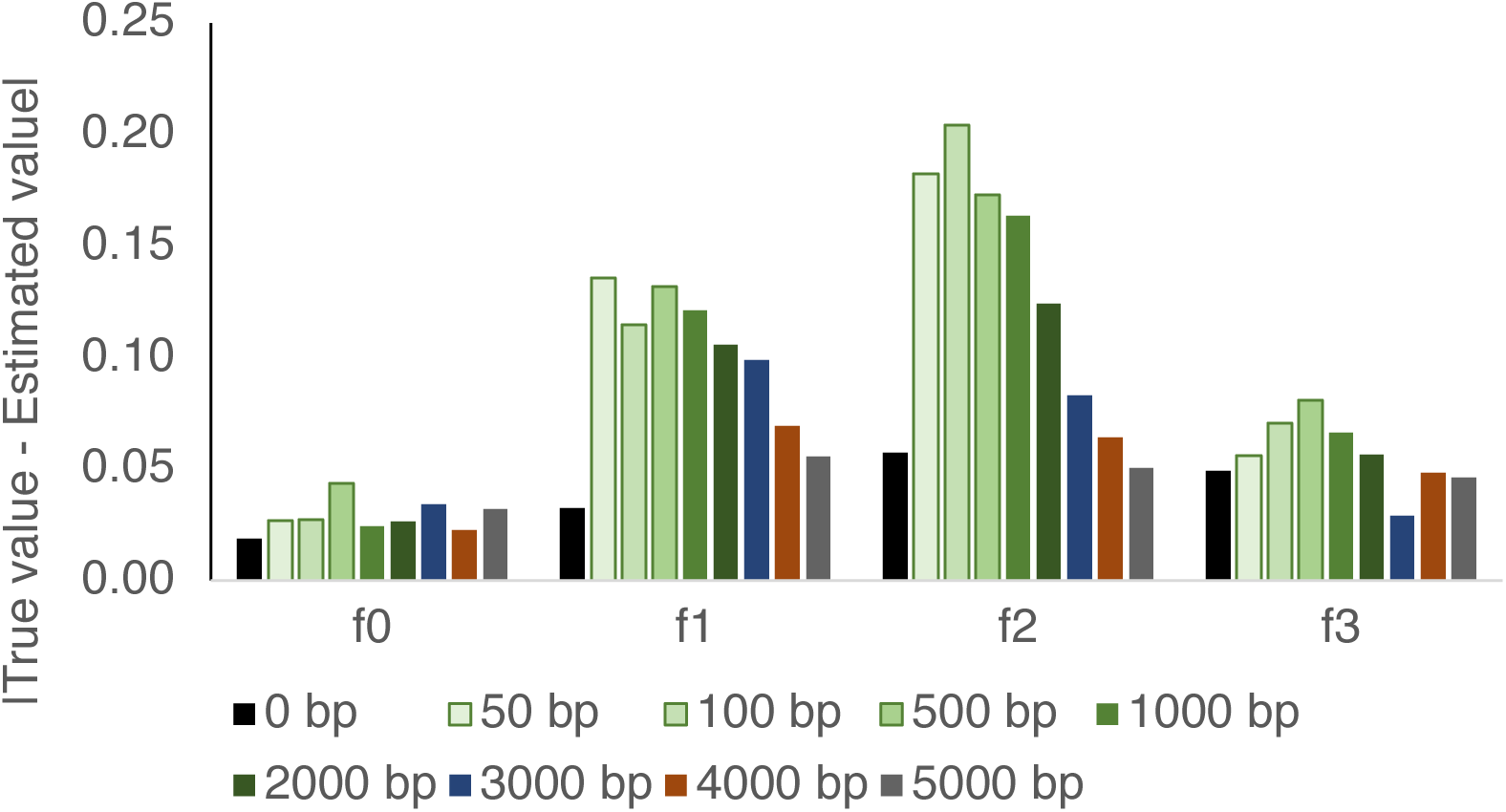
Mis-inference of the DFE in the presence of an additional unaccounted for 1 kb functional region near the target 1kb exon used for inference. The intronic / intergenic distance between the two exons varies from 50-5000 bp, as shown by different colored bars. “0 bp” represents the negative control in which there is no additional 1 kb exon present.

**Figure S6:**
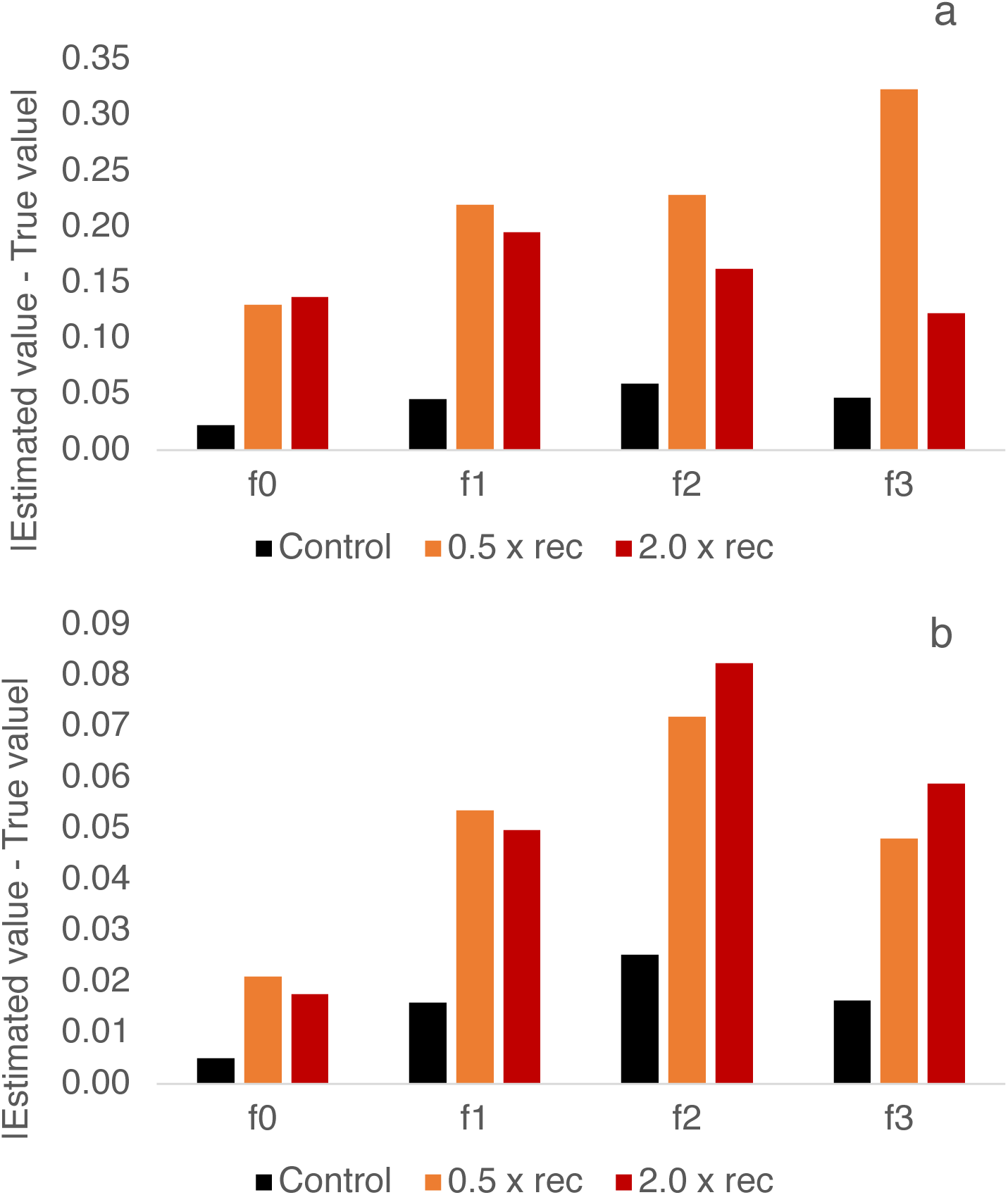
Absolute differences between the true and estimated value of the DFE class, when the true recombination rate is half of that assumed for inference (orange) and when the true value is twice that assumed for inference (red), using a) all statistics and b) statistics only pertaining to the functional region.

**Figure S7:**
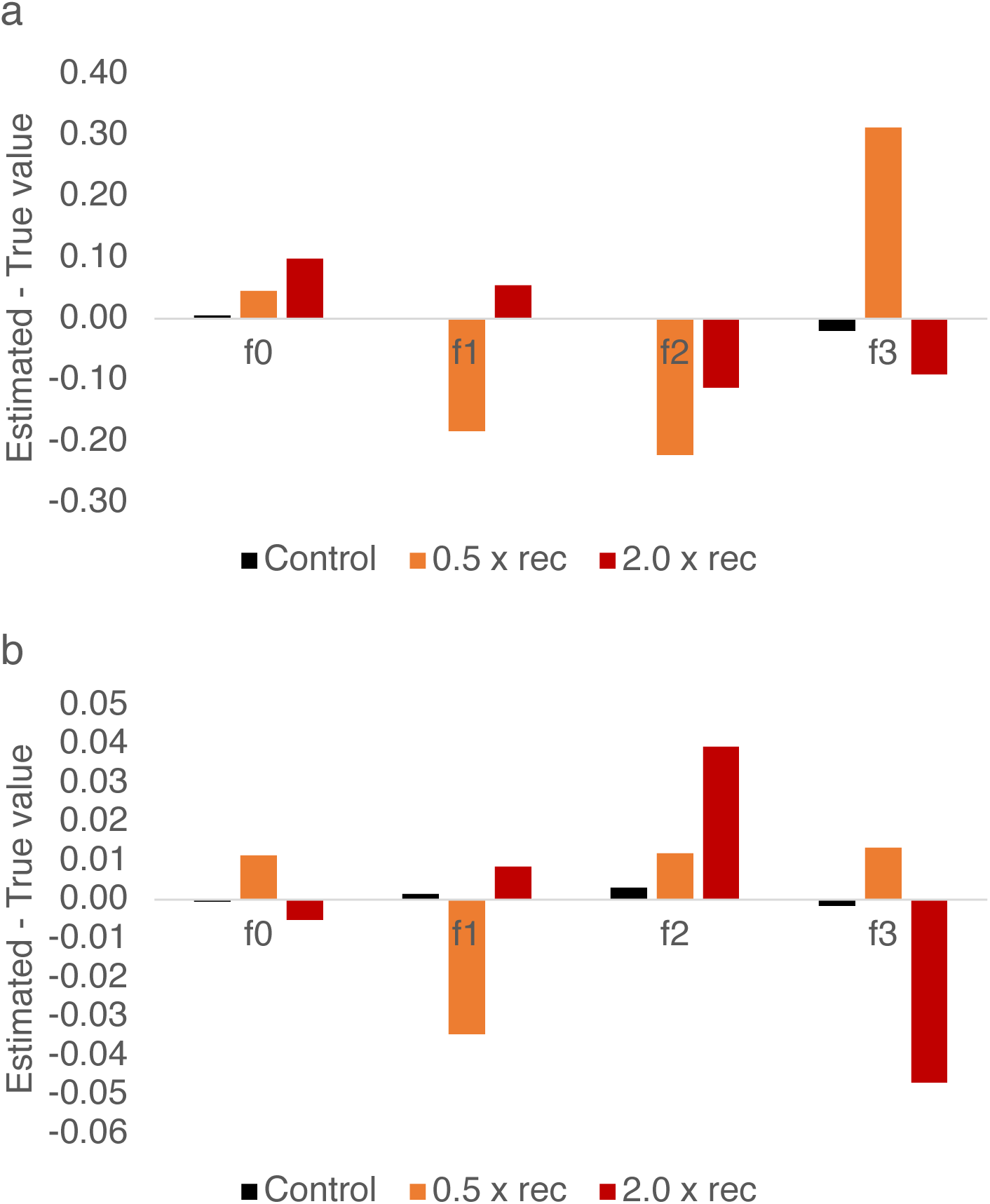
Following Supp Figure 6, the direction of bias in inference of the DFE classes upon mis-specification of the recombination rate is shown, using a) all statistics and b) statistics only pertaining to the functional region.

**Figure S8:**
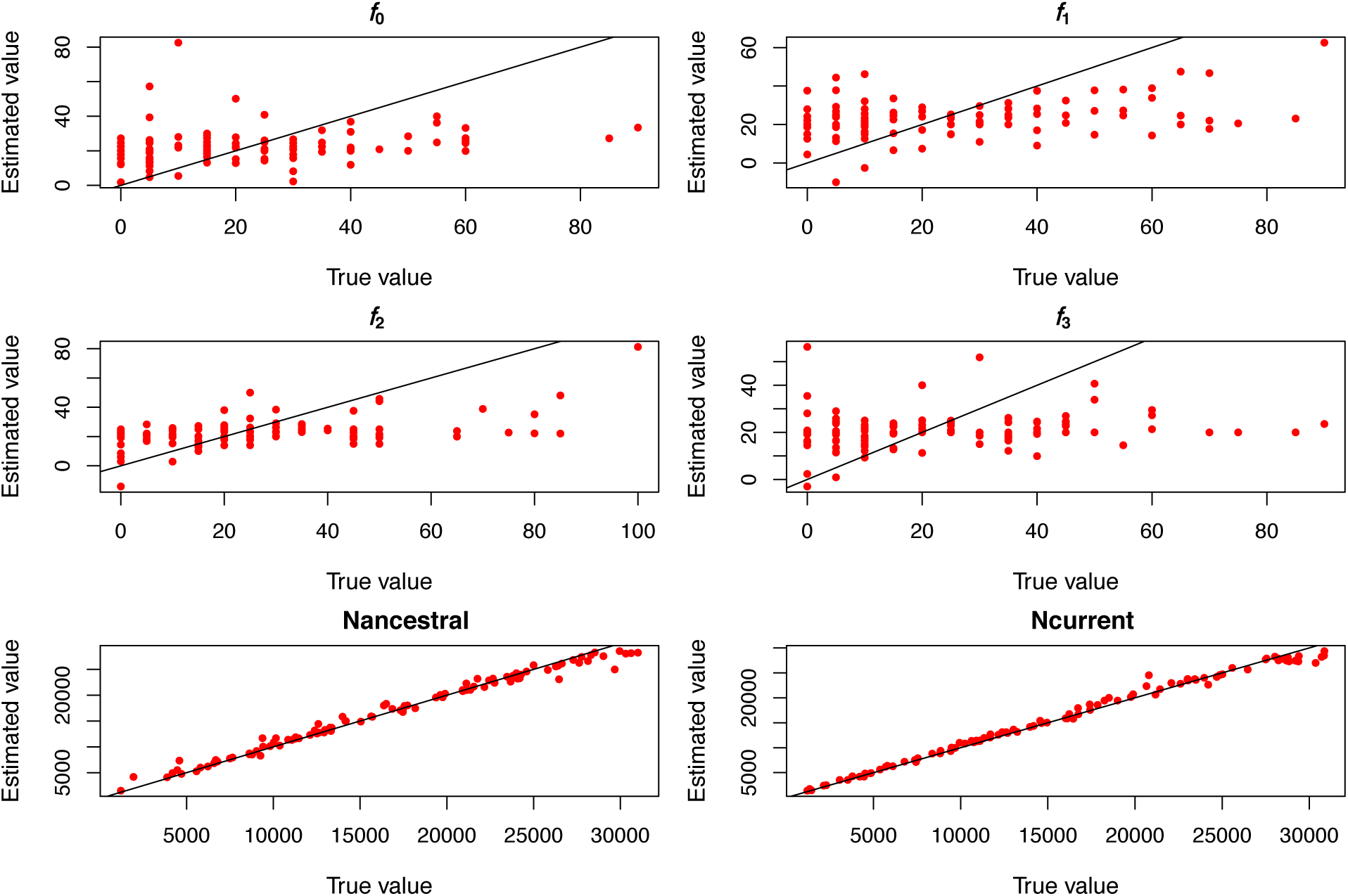
Joint inference of the DFE and demography using statistics only in linked regions (22 summary statistics).

**Figure S9:**
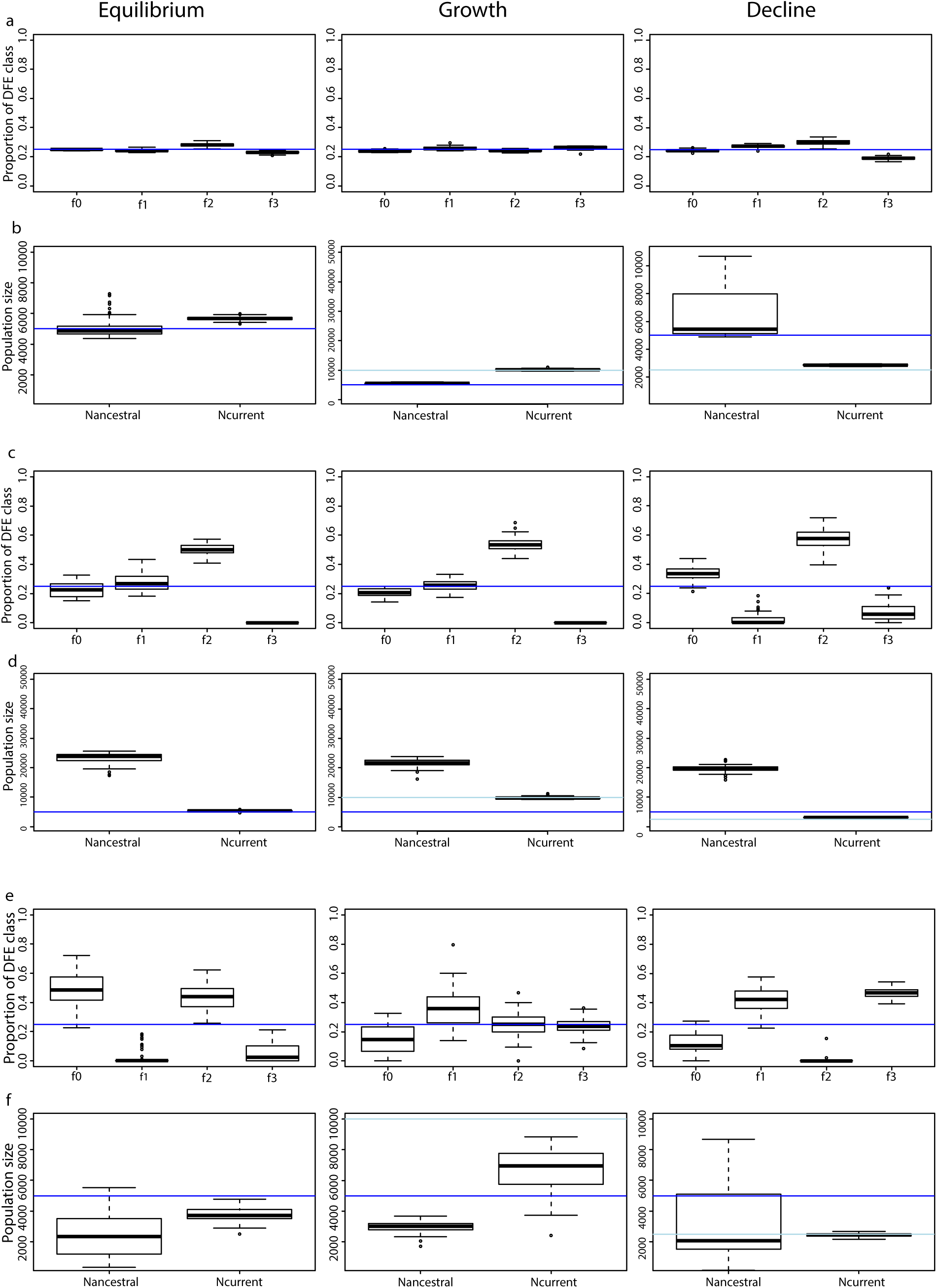
Effect of mis-specification of mutation rate on joint inference of the DFE and demography. (a) Inference of the DFE under equilibrium, 2-fold growth and 2-fold decline, when mutation rate is the same as assumed. Blue line shows the true value. (b) Inference of ancestral and current population sizes under equilibrium, 2-fold growth and 2-fold decline, when mutation rate is the same as assumed. Darker blue and lighter blue lines show the true values of ancestral and current population sizes respectively. (c) Inference of the DFE under equilibrium, 2-fold growth and 2-fold decline when the true mutation rate is twice that assumed. (d) Inference of ancestral and current population sizes under equilibrium, 2-fold growth and 2-fold decline when the true mutation rate is twice that assumed. (e) Inference of the DFE under equilibrium, 2-fold growth and 2-fold decline when the true mutation rate is half of that assumed. (f) Inference of ancestral and current population sizes under equilibrium, 2-fold growth and 2-fold decline when the true mutation rate is half of that assumed.

**Figure S10:**
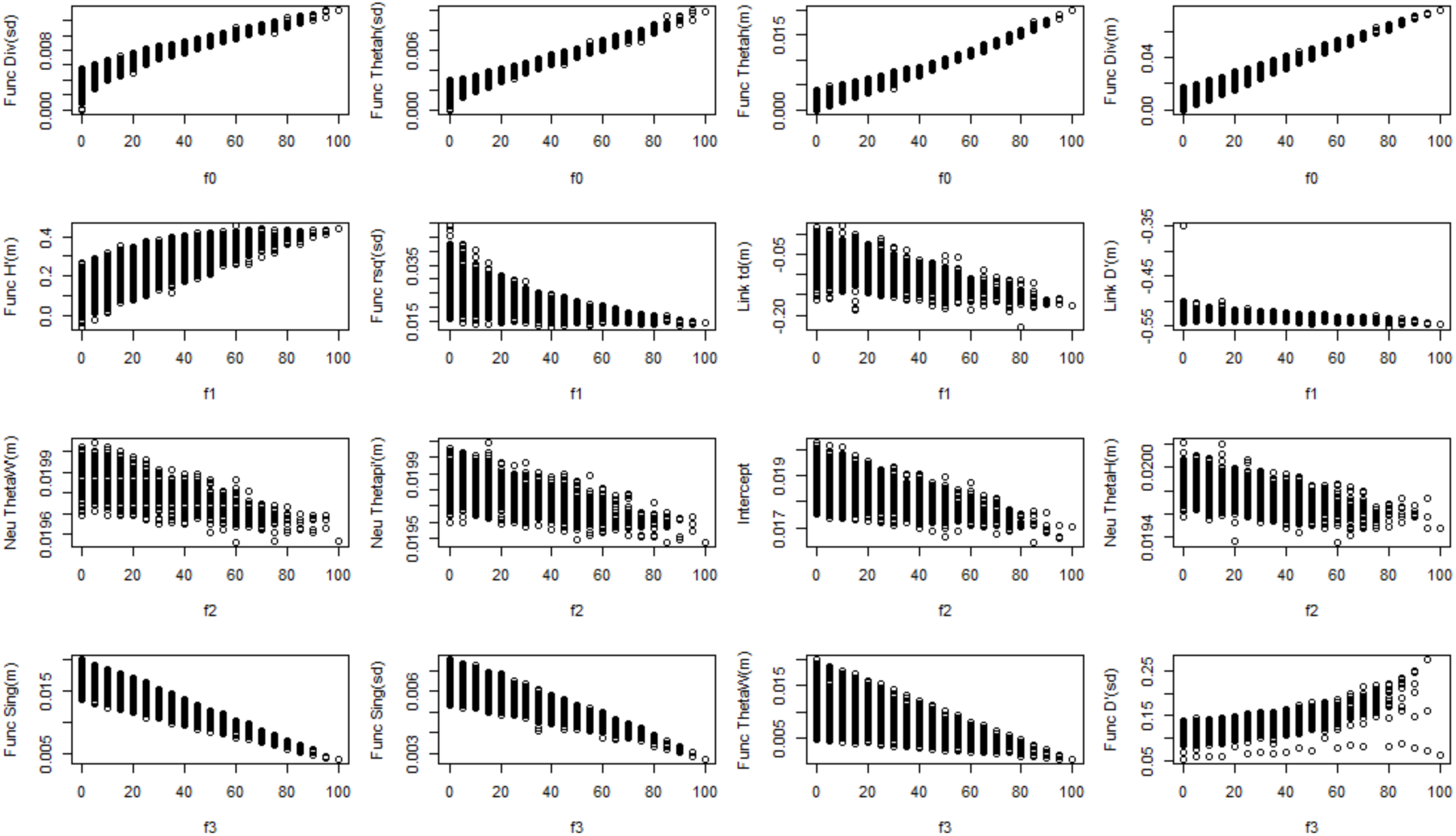
Correlations of the top 4 statistics with parameters characterizing the DFE under demographic equilibrium. “Func” corresponds to the functional region, “Link” to the immediately linked region and “Neu” to the less linked region.

**Figure S11:**
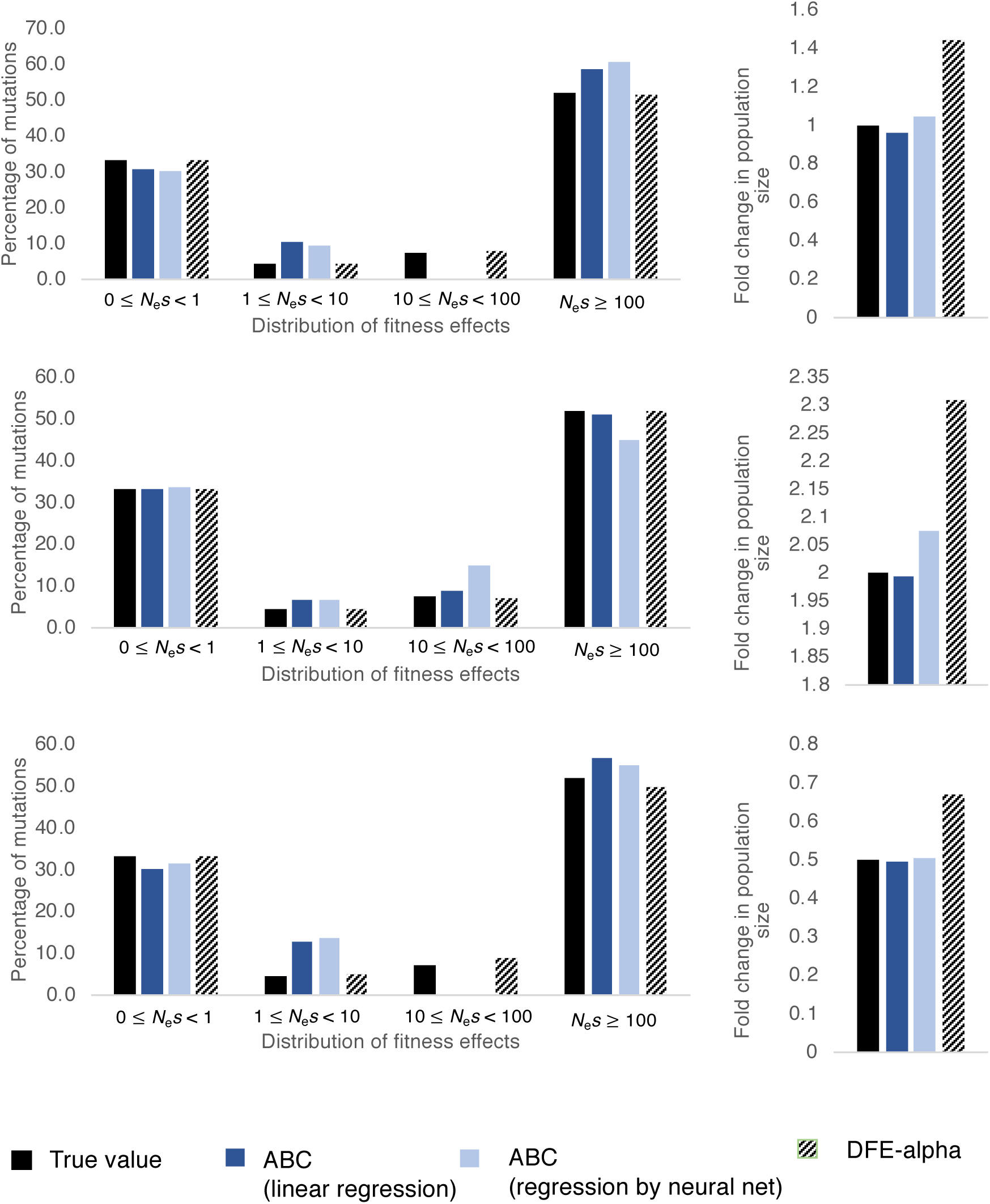
Inference of demography and the DFE by the approach proposed here and DFE-alpha, when the true shape of the DFE is gamma distributed, for equilibrium (top panel), growth (middle panel), and decline (bottom panel). Solid black bars show the true value simulated, dark blue bars show our ABC performance using ridge regression, light blue bars show the ABC performance using linear regression aided by neural net. Patterned bars show the performance of DFE-alpha.

**Figure S12:**
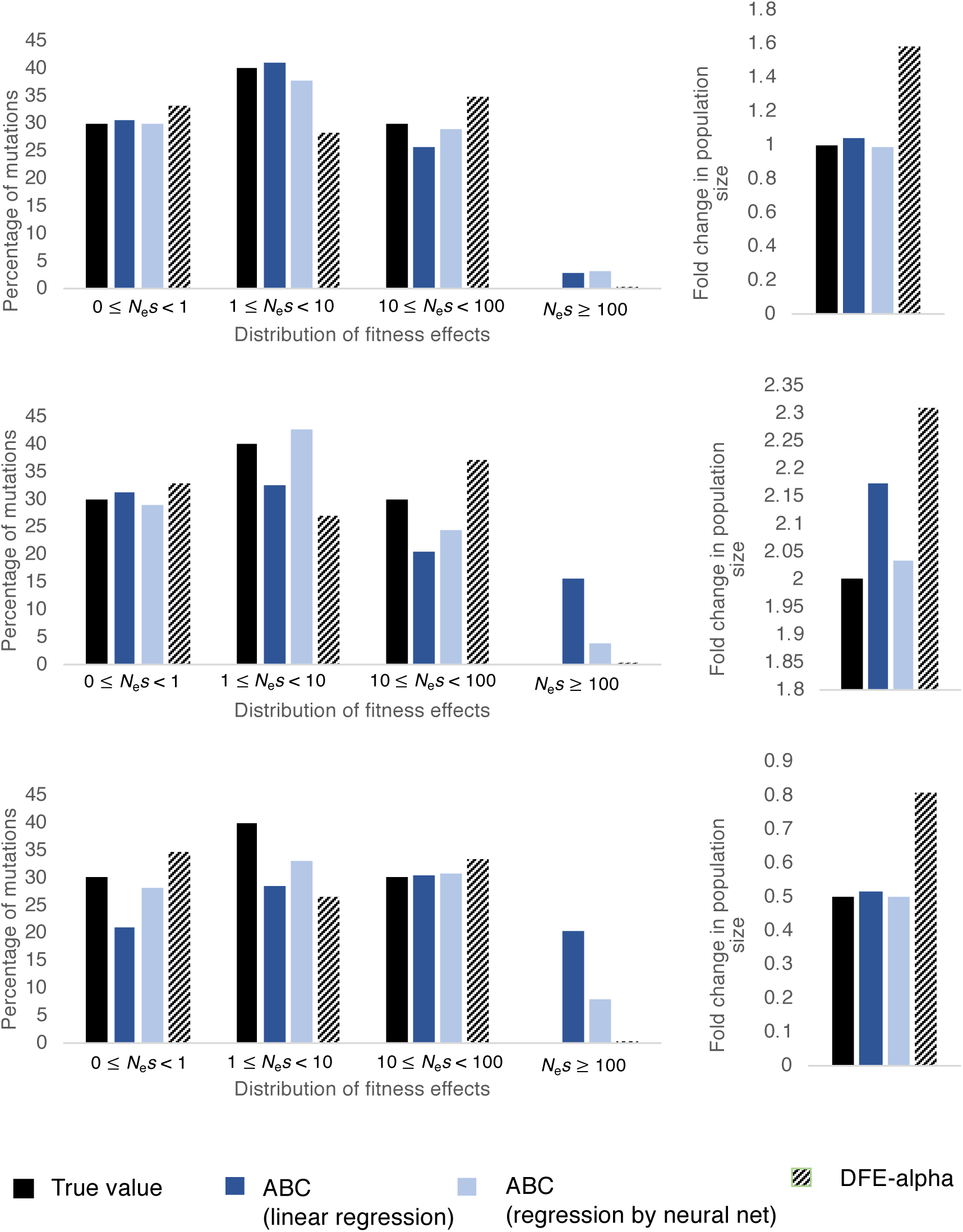
Inference of demography and the DFE when the true shape of the DFE is discrete and skewed towards slightly deleterious class of mutations, for demographic equilibrium (top panel), population size growth (middle panel), and population size decline (bottom panel). Solid black bars show the true value simulated, dark blue bars show the ABC performance using ridge regression, light blue bars show the ABC performance using linear regression aided by neural net. Patterned bars show the performance of DFE-alpha.

**Figure S13:**
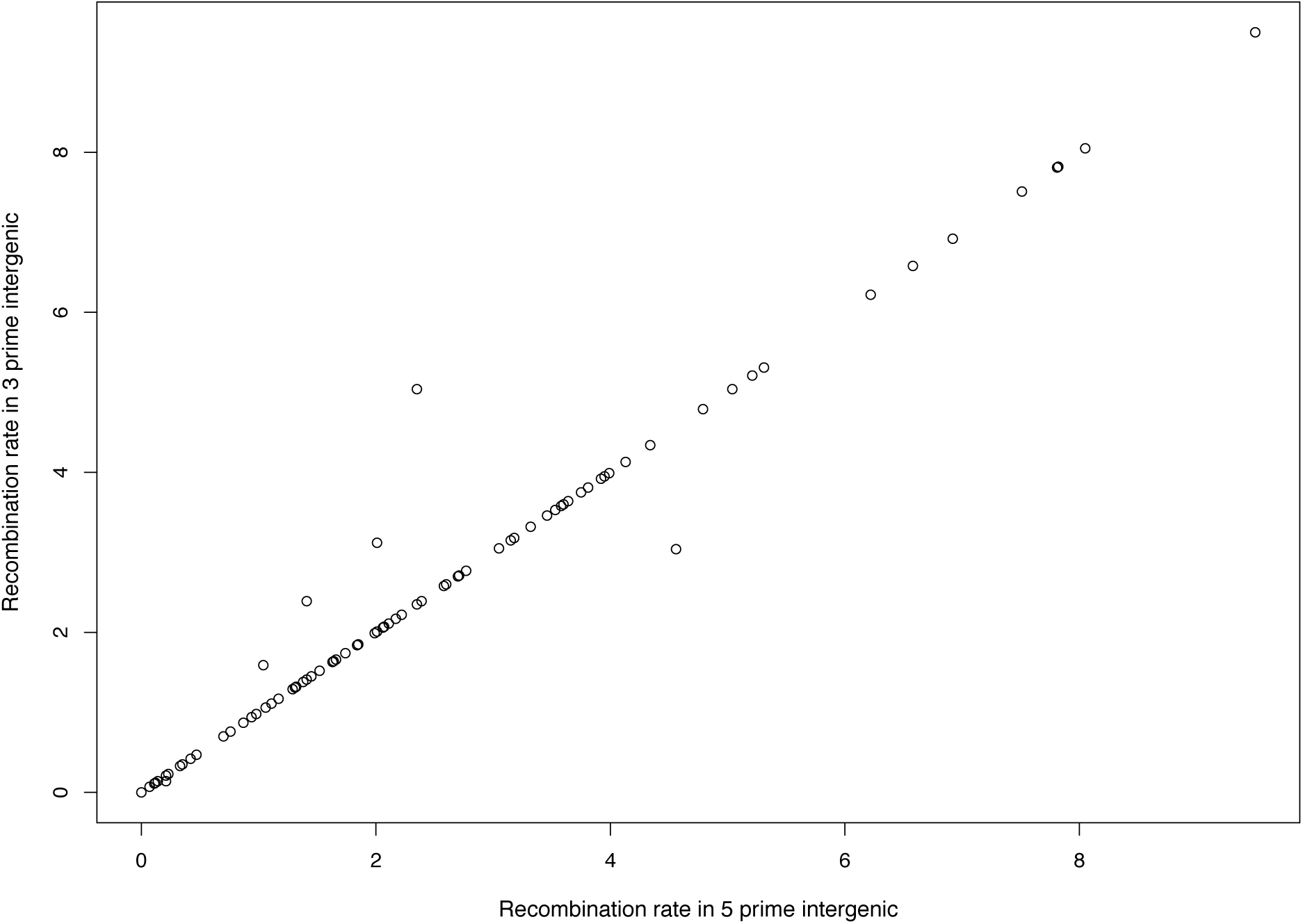
Correlation of recombination rates in 5′ flanking intergenic regions with those in 3′ flanking intergenic regions over all 94 exons chosen for analysis in *D. melanogaster*.

**Figure S14:**
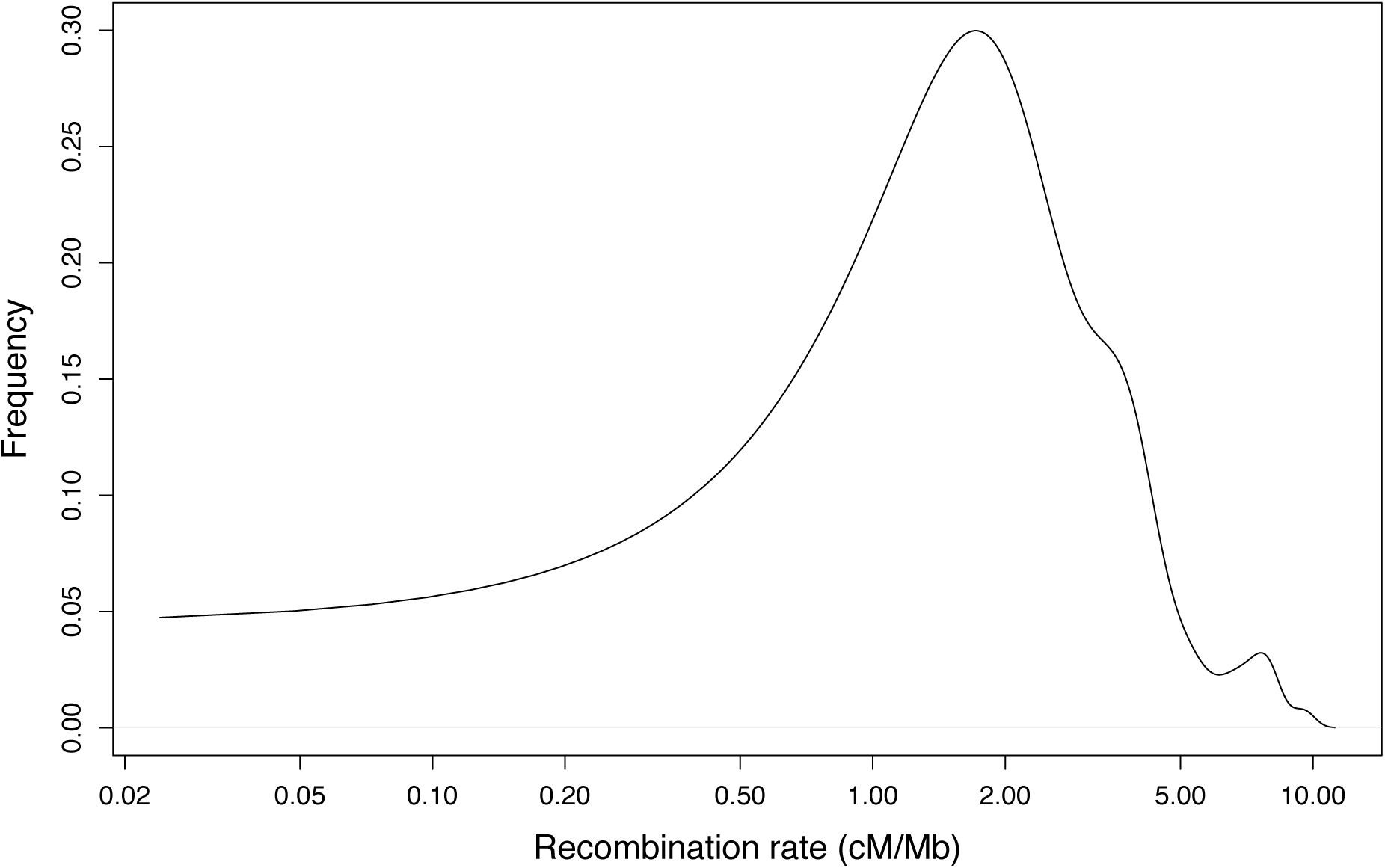
Distribution of the rate of recombination in cM/Mb for all 94 exons selected for analysis in *D. melanogaster*. Note that the x-axis is on a logarithmic scale.

**Figure S15:**
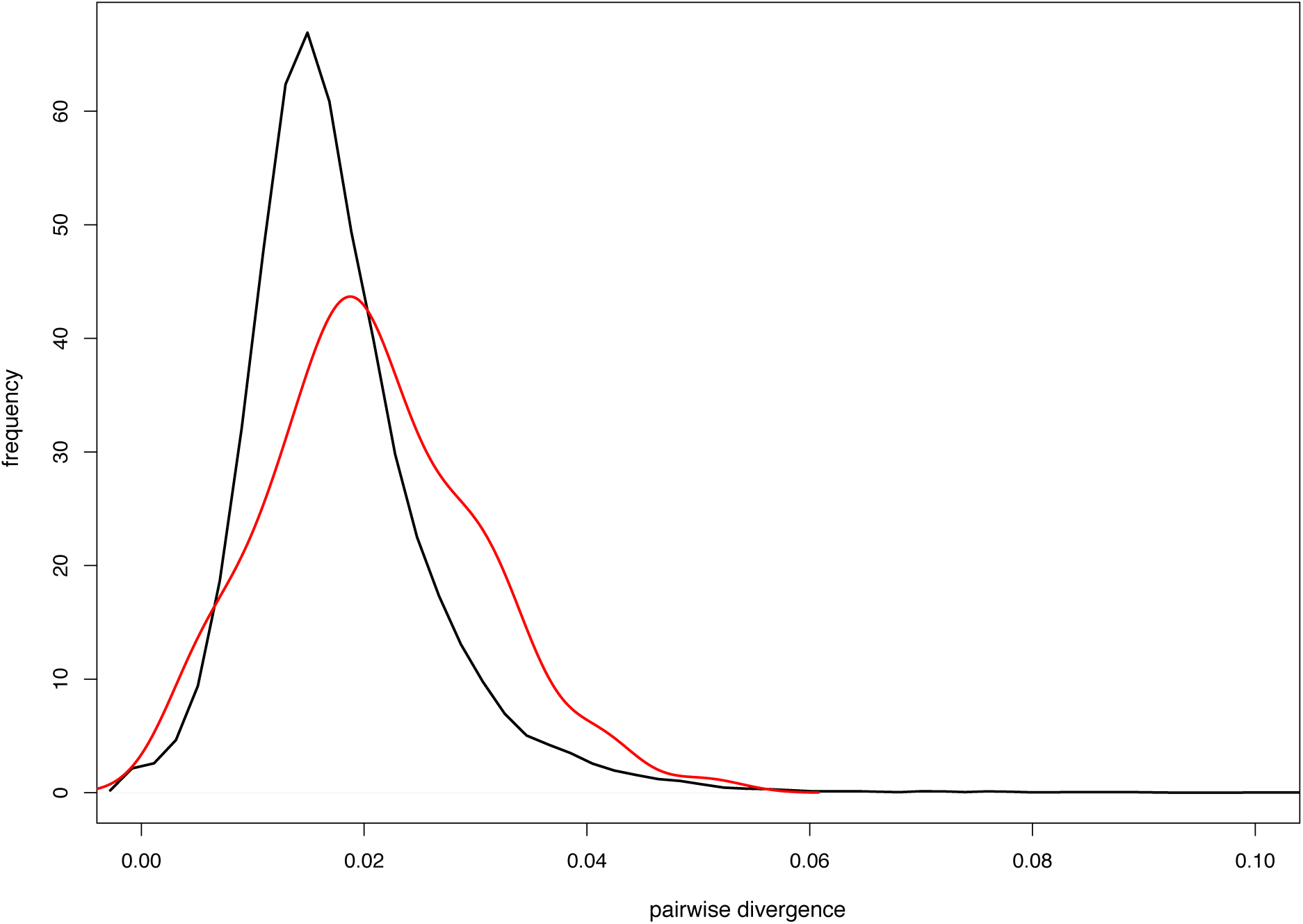
Distribution of divergence per site of single-exon genes that have flanking intergenic regions larger than 4 kb (in red), and for all genes (in black), from *D. melanogaster*.

**Figure S16:**
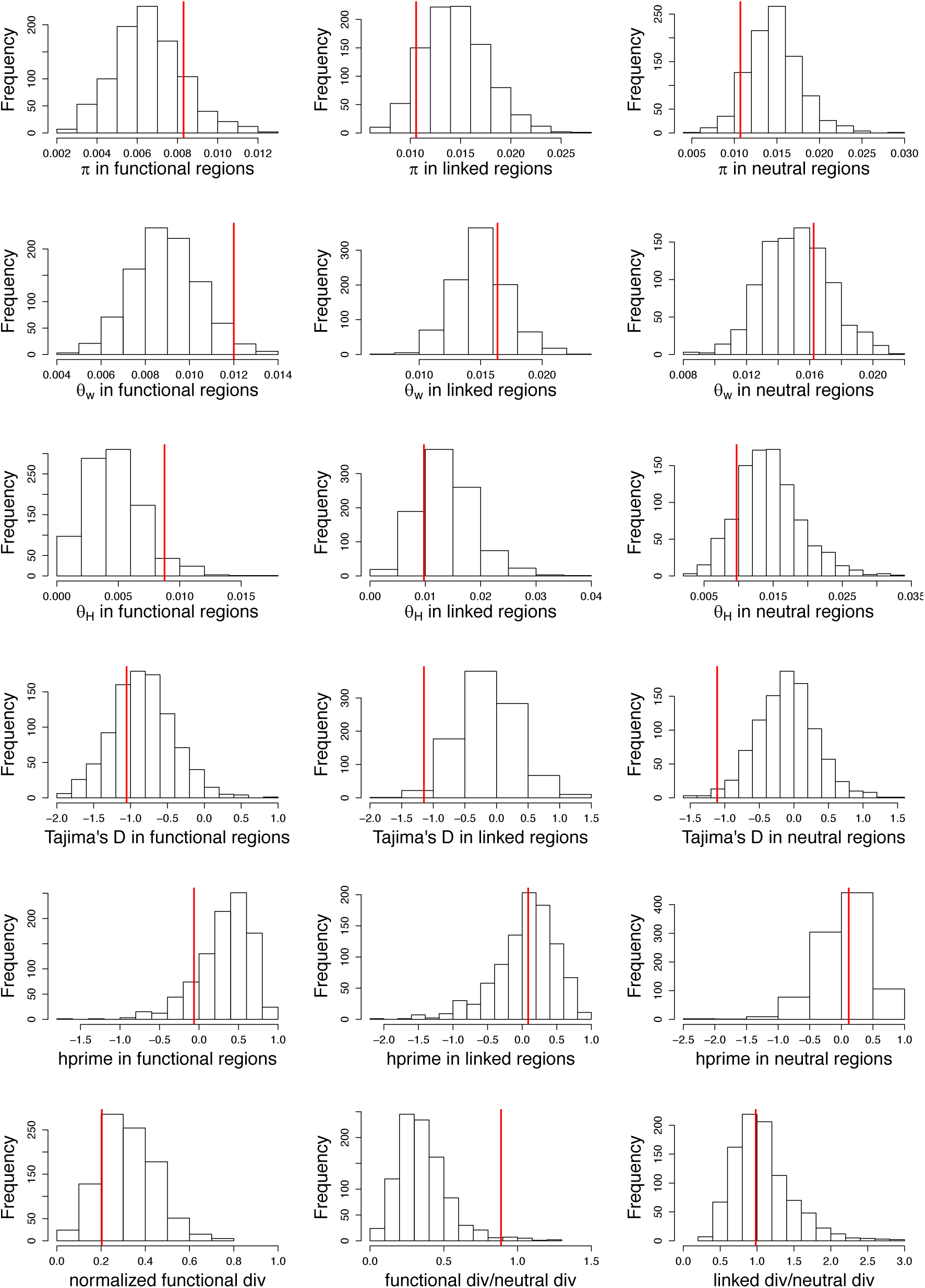
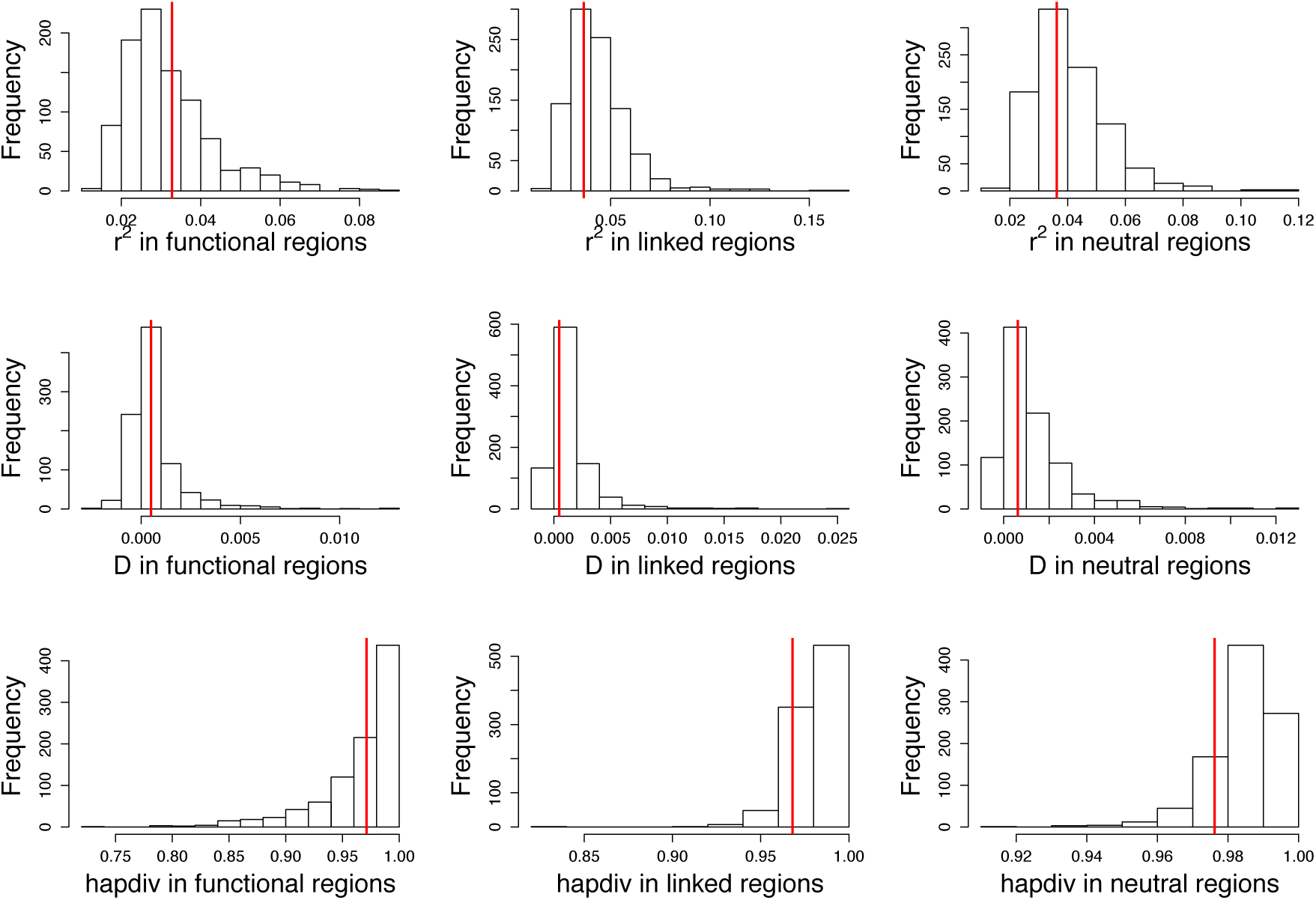
Distributions of summary statistics calculated from 94 exons simulated with 100 replicates each using our inferred model (*i.e*., *f*_0_ = 0.25, *f*_1_=0.49, *f*_2_=0.04, *f*_3_=0.22, *N*_anc_= 1,225,393, *N*_cur_ = 1,357,760). Red lines indicate the value observed in 76 individuals of *D. melanogaster* from Zambia, after excluding sites with phastCons score **≥** 0.8.

**Figure S17:**
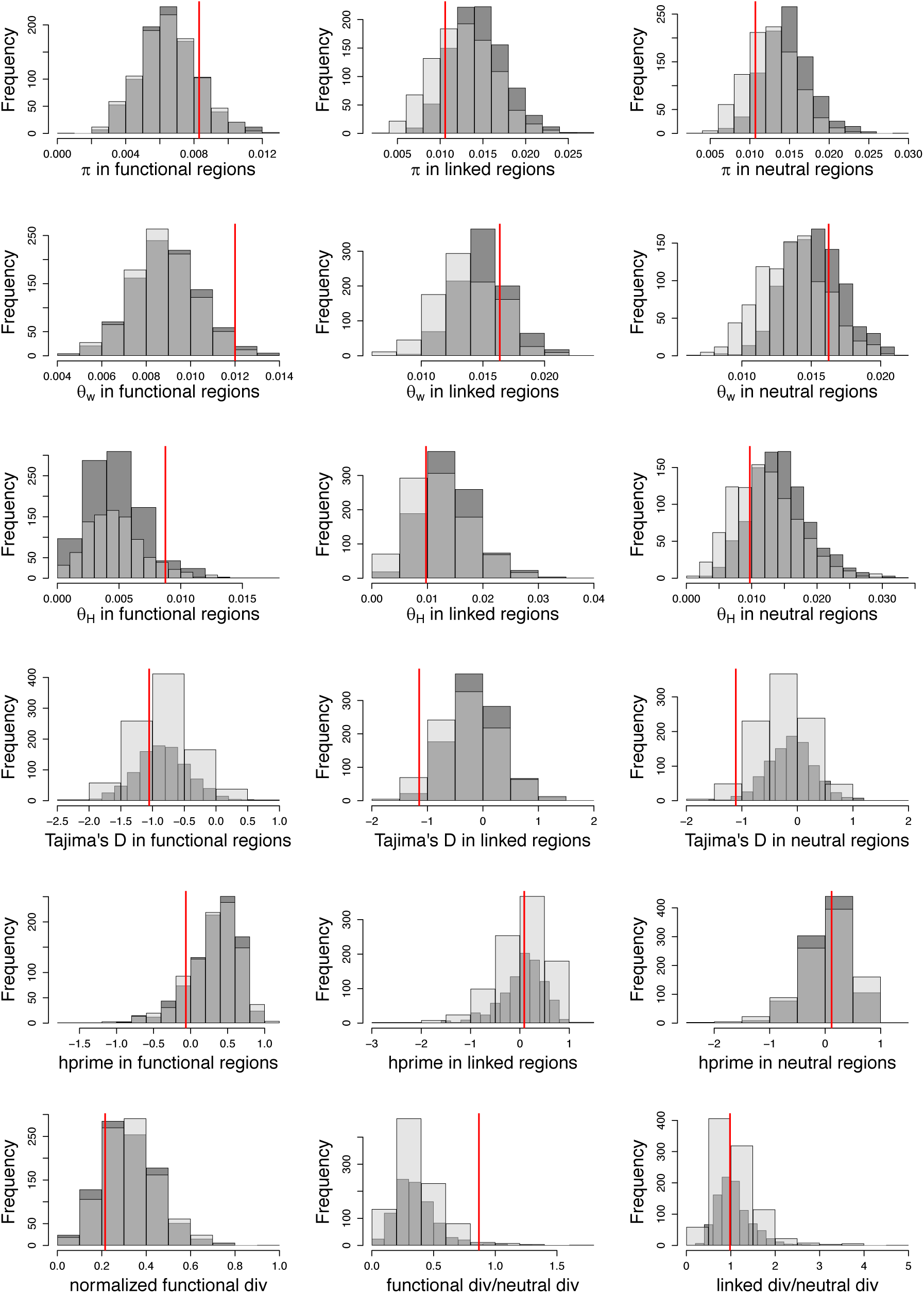
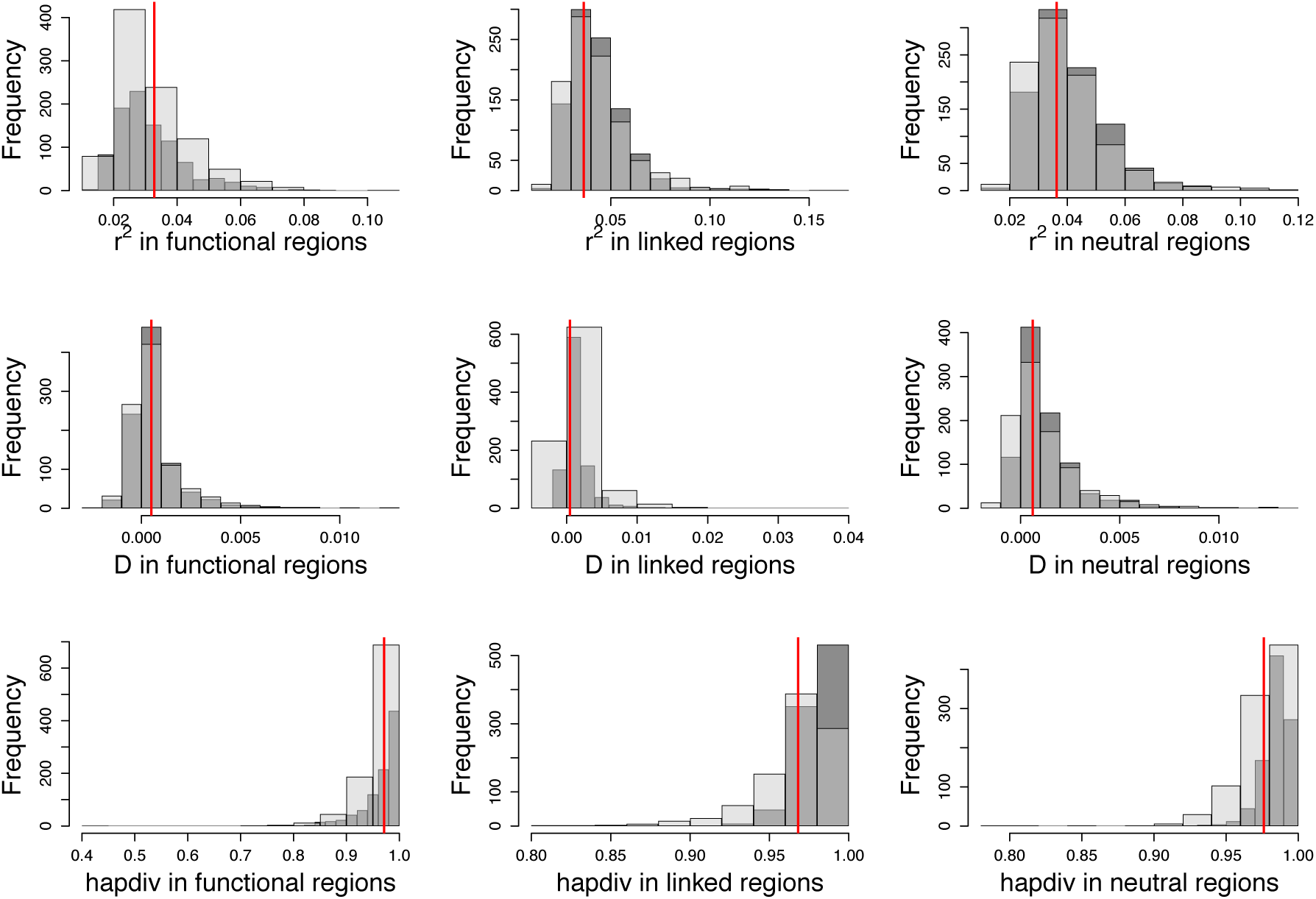
Distribution of summary statistics calculated from 94 exons simulated with 100 replicates each using our inferred model (*i.e*., *f*_0_ = 0.25, *f*_1_=0.49, *f*_2_=0.04, *f*_3_=0.22, *N*_anc_= 1,225,393, *N*_cur_ = 1,357,760). In this case, conserved elements that represent 40% of non-coding regions were simulated as experiencing purifying selection with the class of mutations that result in the strongest BGS effects (−100<2*N*_e_*s*<-10). and these sites were masked while calculating statistics. Red line indicates the value observed in 76 individuals of *D. melanogaster* from Zambia, after excluding sites with phastCons score **≥** 0.8. Dark grey bars represent no selection on non-coding regions and light grey bars represent simulations with selection on non-coding regions.

**Figure S18:**
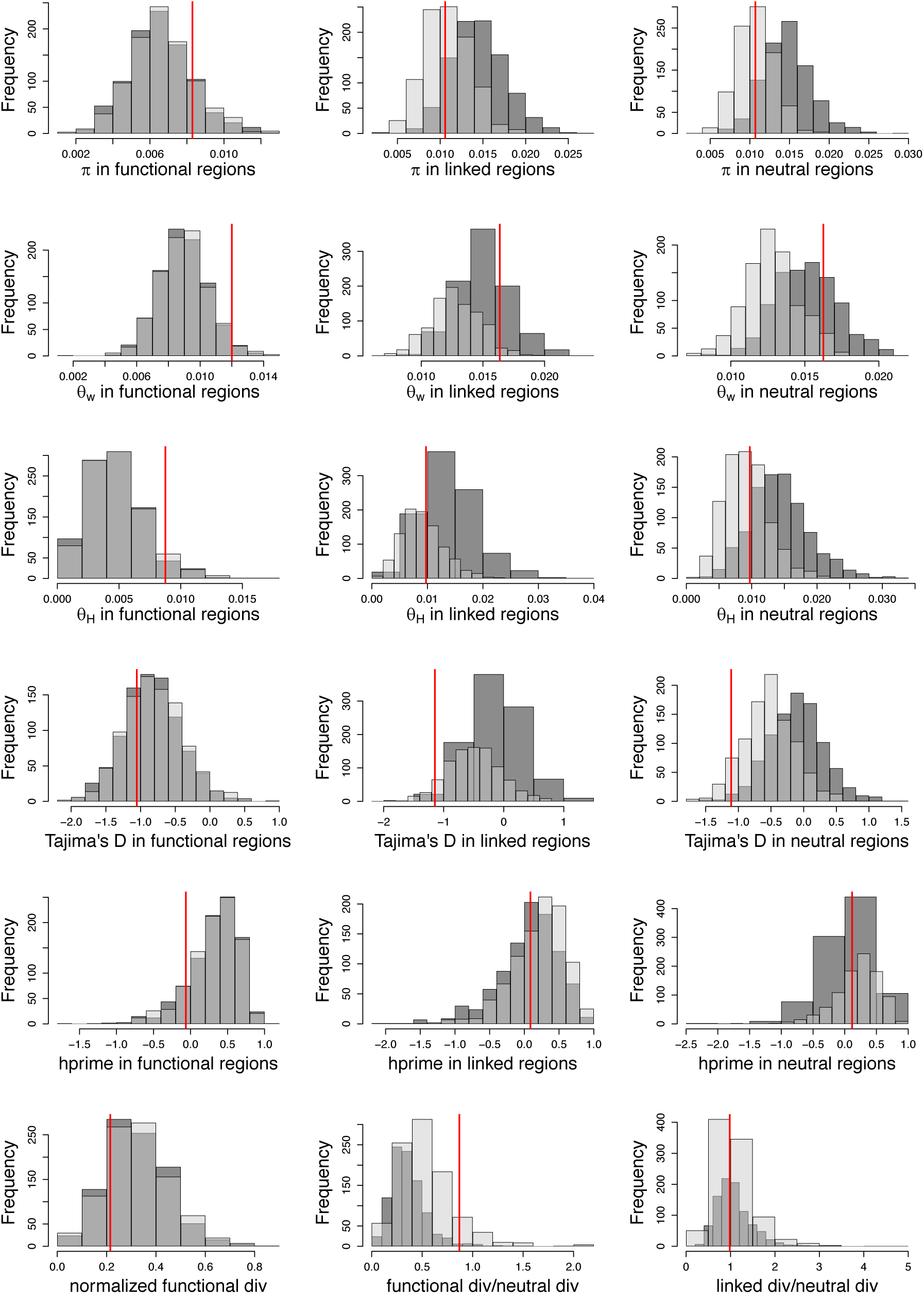
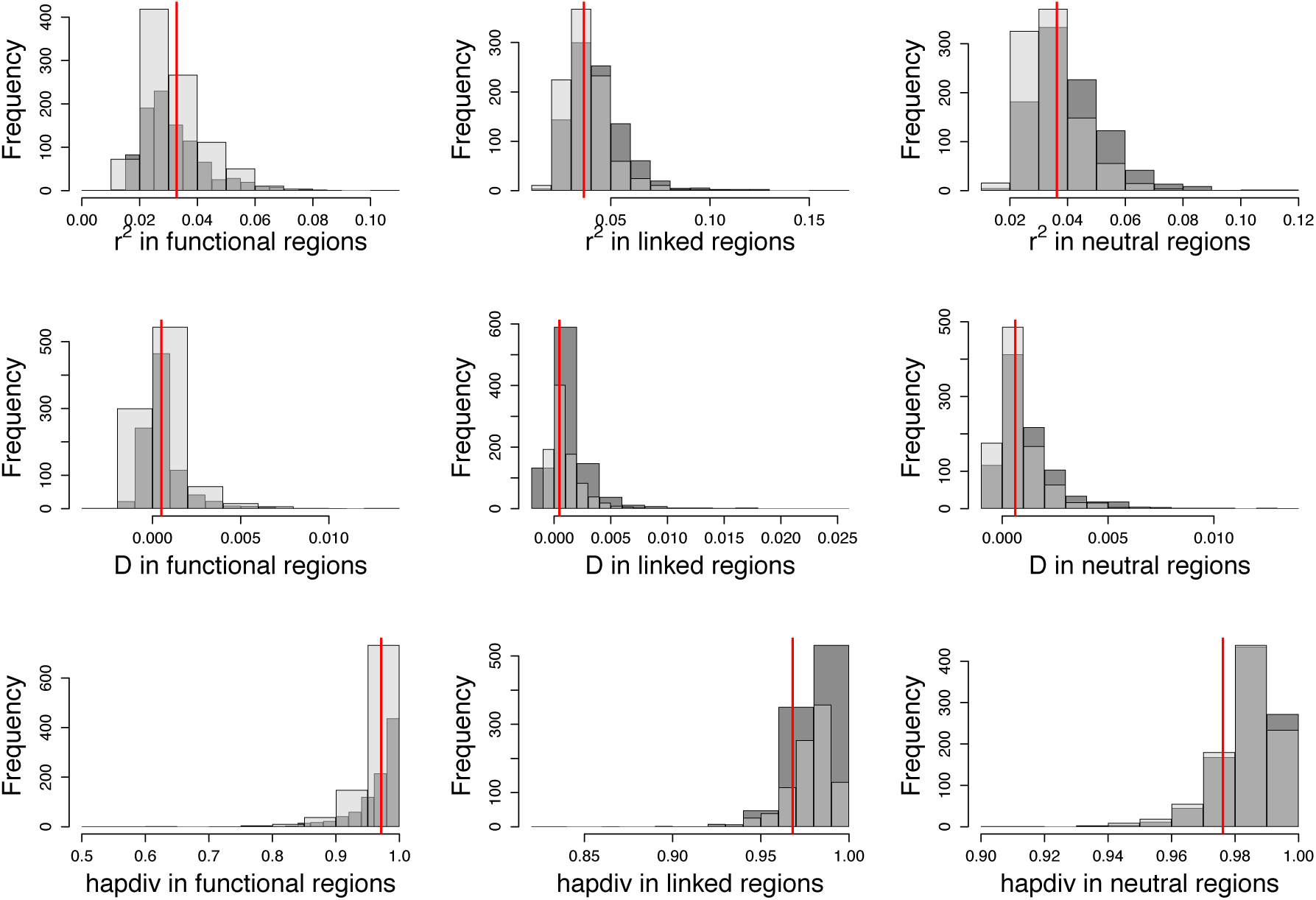
Distribution of summary statistics calculated from 94 exons simulated with 100 replicates each using our inferred model (*i.e*., *f*_0_ = 0.25, *f*_1_=0.49, *f*_2_=0.04, *f*_3_=0.22, *N*_anc_= 1,225,393, *N*_cur_ = 1,357,760). In this case, conserved elements that represent 40% of non-coding regions were simulated as experiencing weak purifying selection (−10<2*N*_e_*s*<-1) and these sites were included while calculating statistics. Red line indicates the value observed in 76 individuals of *D. melanogaster* from Zambia, after excluding sites with phastCons score **≥** 0.8. Dark grey bars represent no selection on non-coding regions and light grey bars represent simulations with selection on non-coding regions.

**Figure S19:**
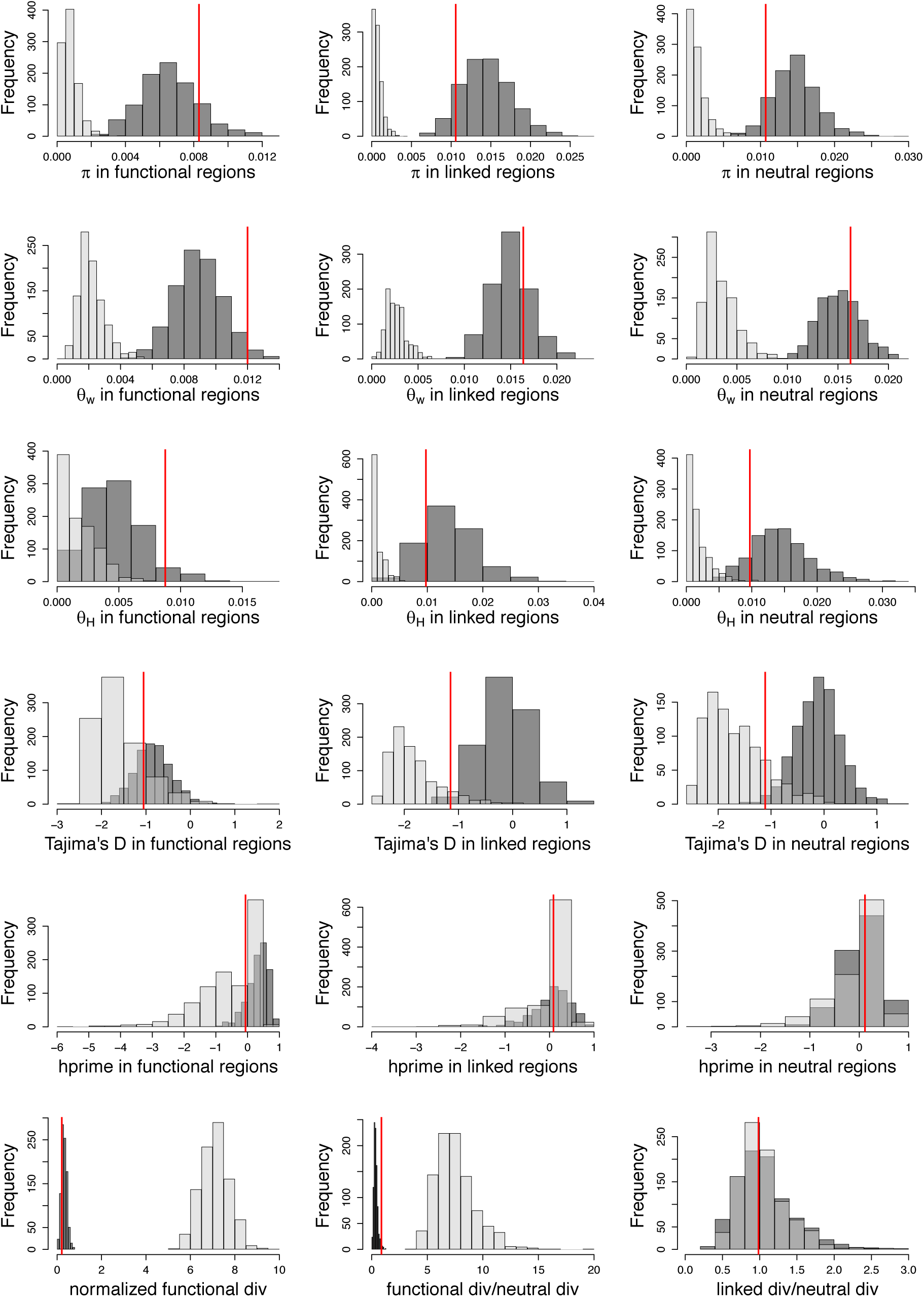
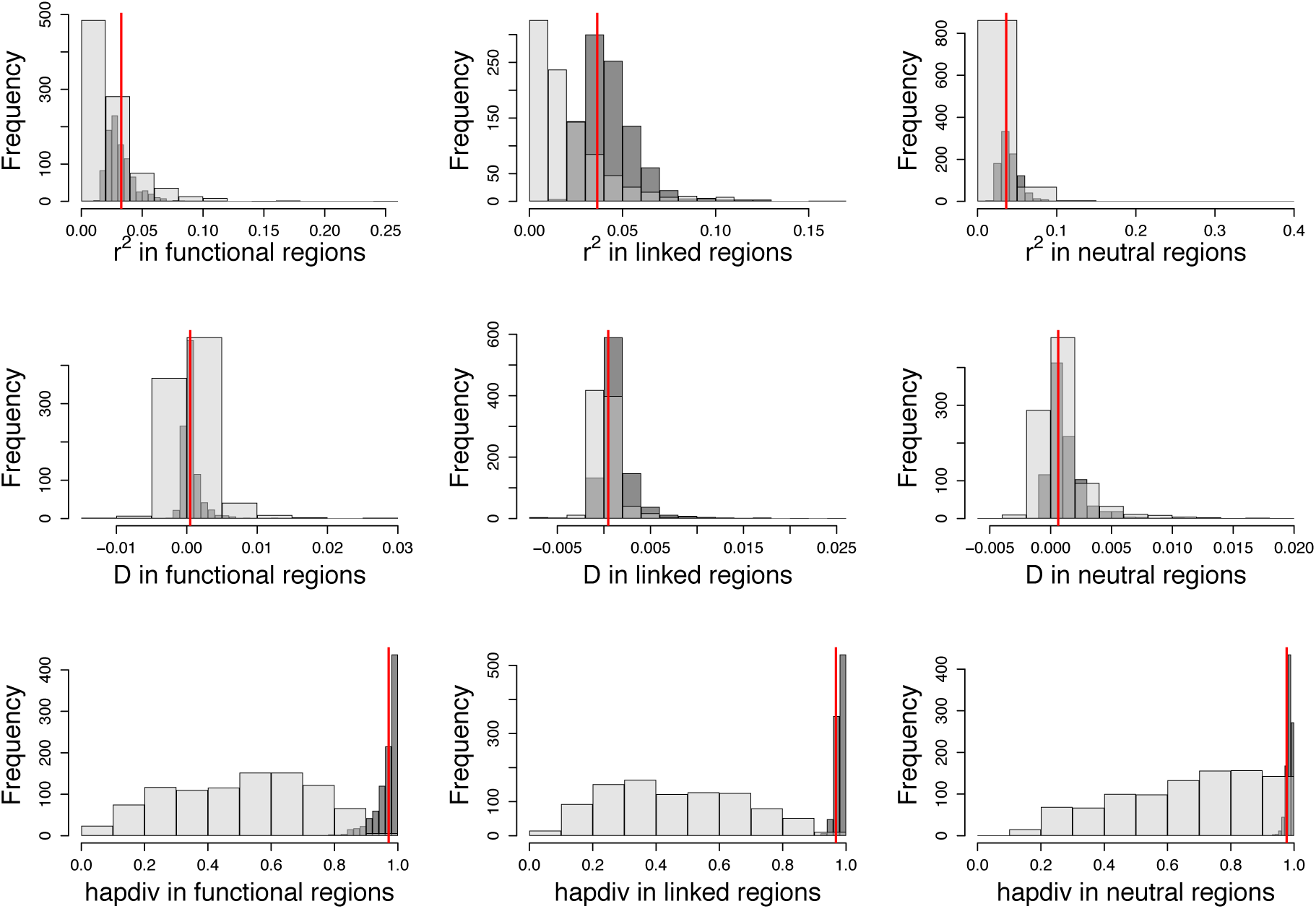
Distribution of summary statistics calculated from 94 exons simulated with 100 replicates each using our inferred model (*i.e*., *f*_0_ = 0.25, *f*_1_=0.49, *f*_2_=0.04, *f*_3_=0.22, *N*_anc_= 1,225,393, *N*_cur_= 1,357,760). Functional regions were simulated as experiencing rare (1%) and strong positive selection (2*N*_anc_*s* = 1000). Red lines indicate the value observed in 76 individuals of *D. melanogaster* from Zambia, after excluding sites with phastCons score **≥** 0.8. Dark grey bars represent no positive selection and light grey bars represent simulations with positive selection in functional regions.

**Figure S20:**
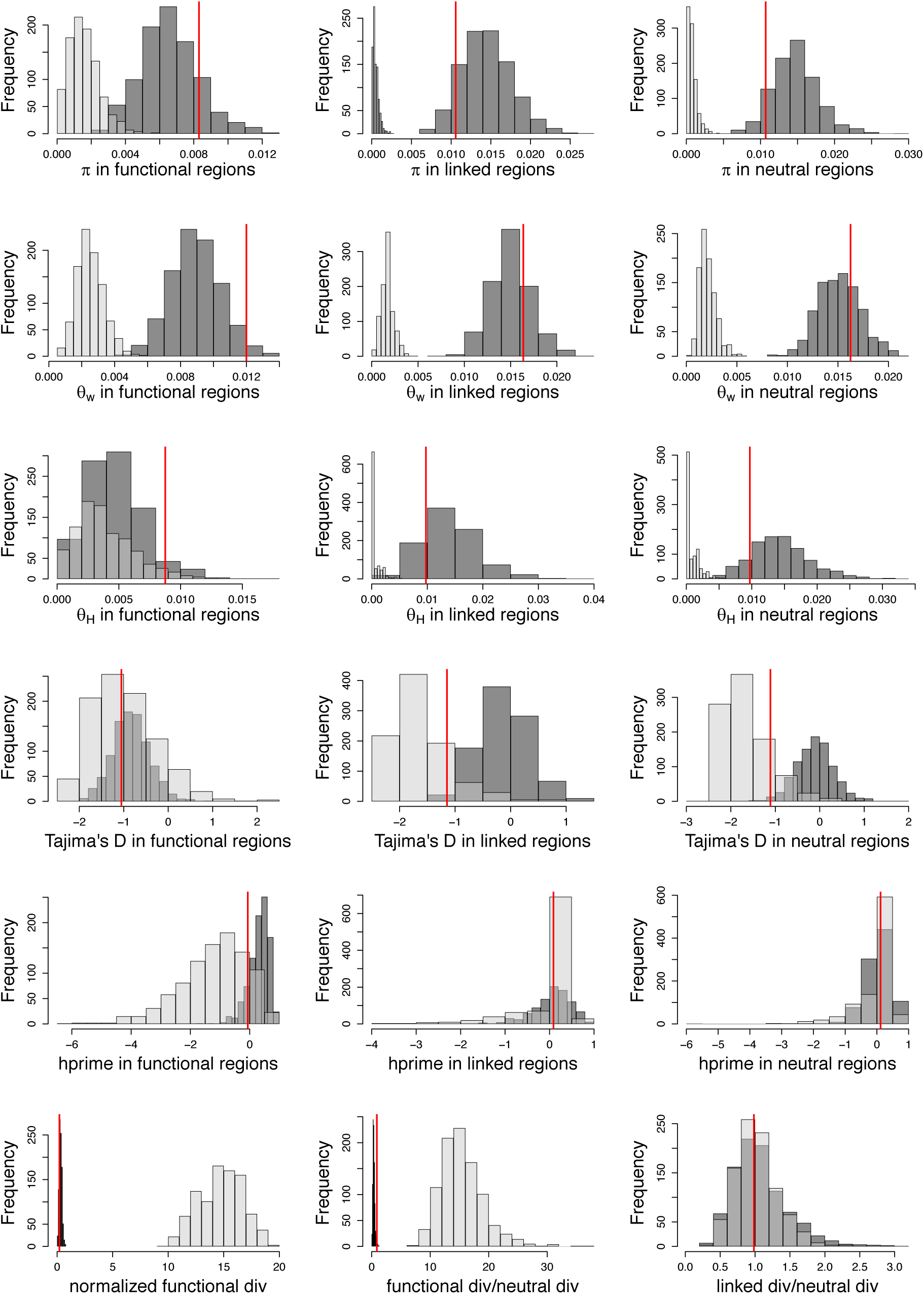
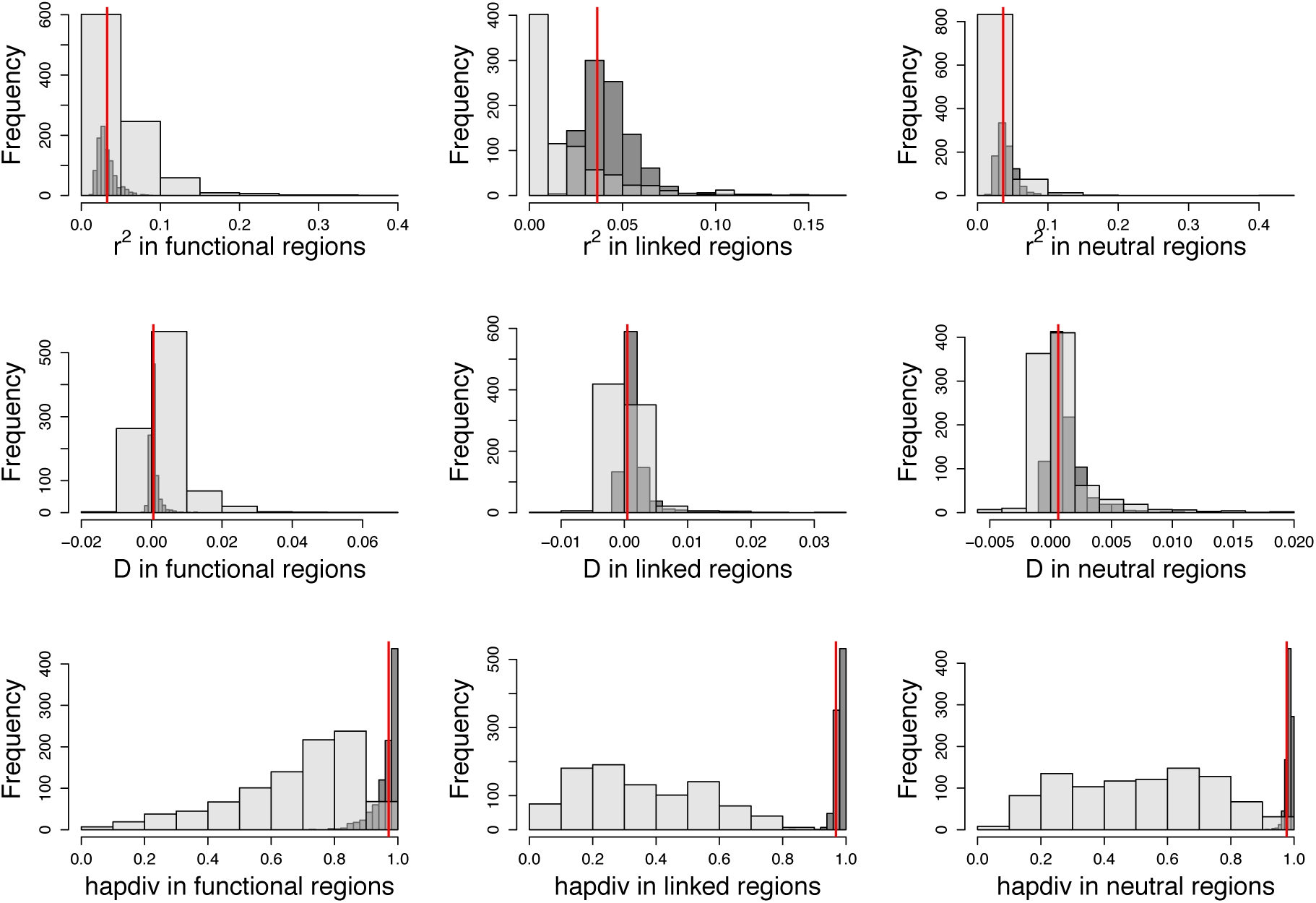
Distribution of summary statistics calculated from 94 exons simulated with 100 replicates each using our inferred model (*i.e*., *f*_0_ = 0.25, *f*_1_=0.49, *f*_2_=0.04, *f*_3_=0.22, *N*_anc_= 1,225,393, *N*_cur_= 1,357,760). Functional regions were simulated to experience common (5%) and strong positive selection (2*N*_anc_*s* = 1000). Red line indicates the value observed in 76 individuals of *D. melanogaster* from Zambia, after excluding sites with phastCons score **≥** 0.8. Dark grey bars represent no positive selection and light grey bars represent simulations with positive selection in functional regions.

**Figure S21:**
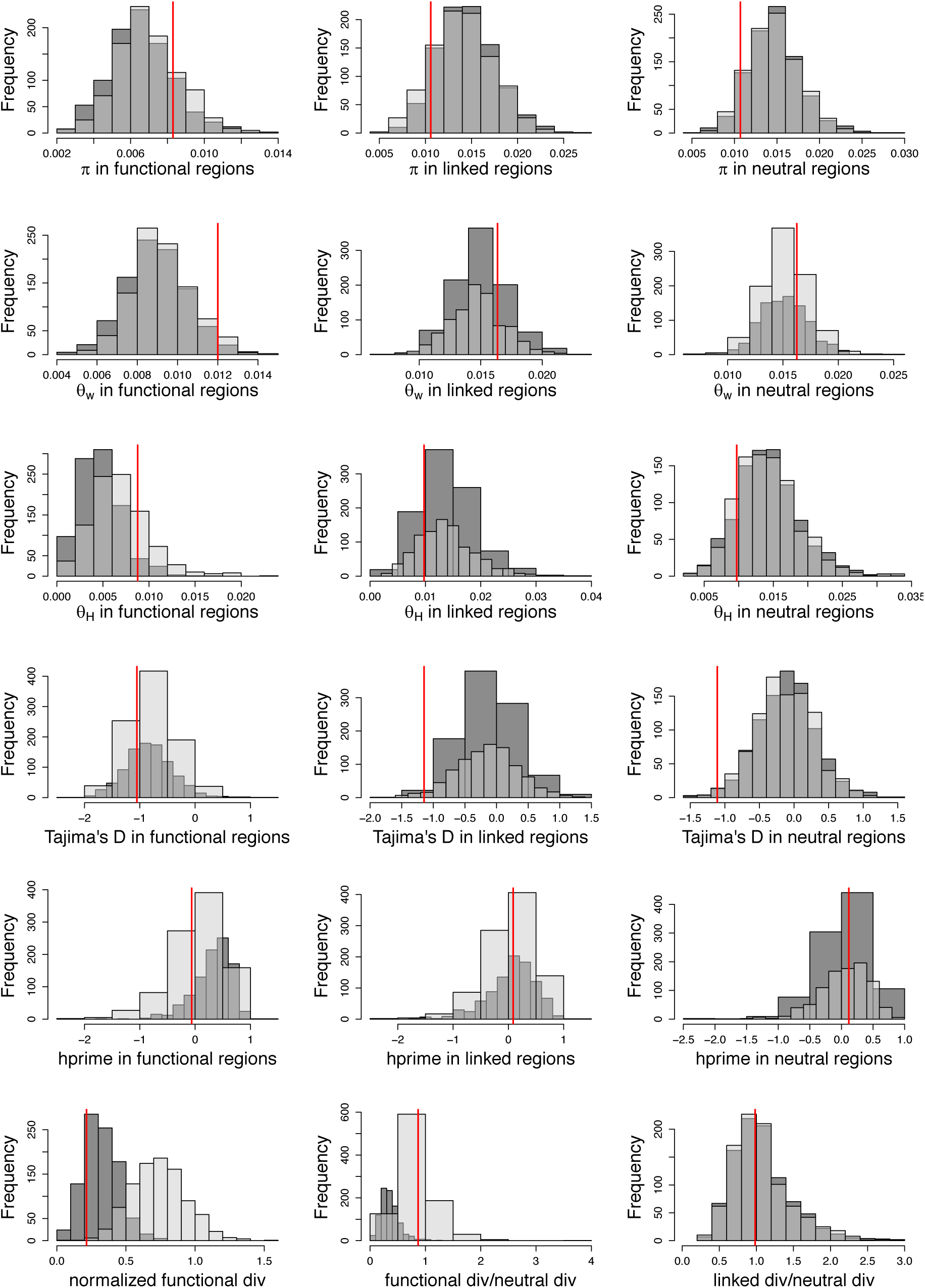
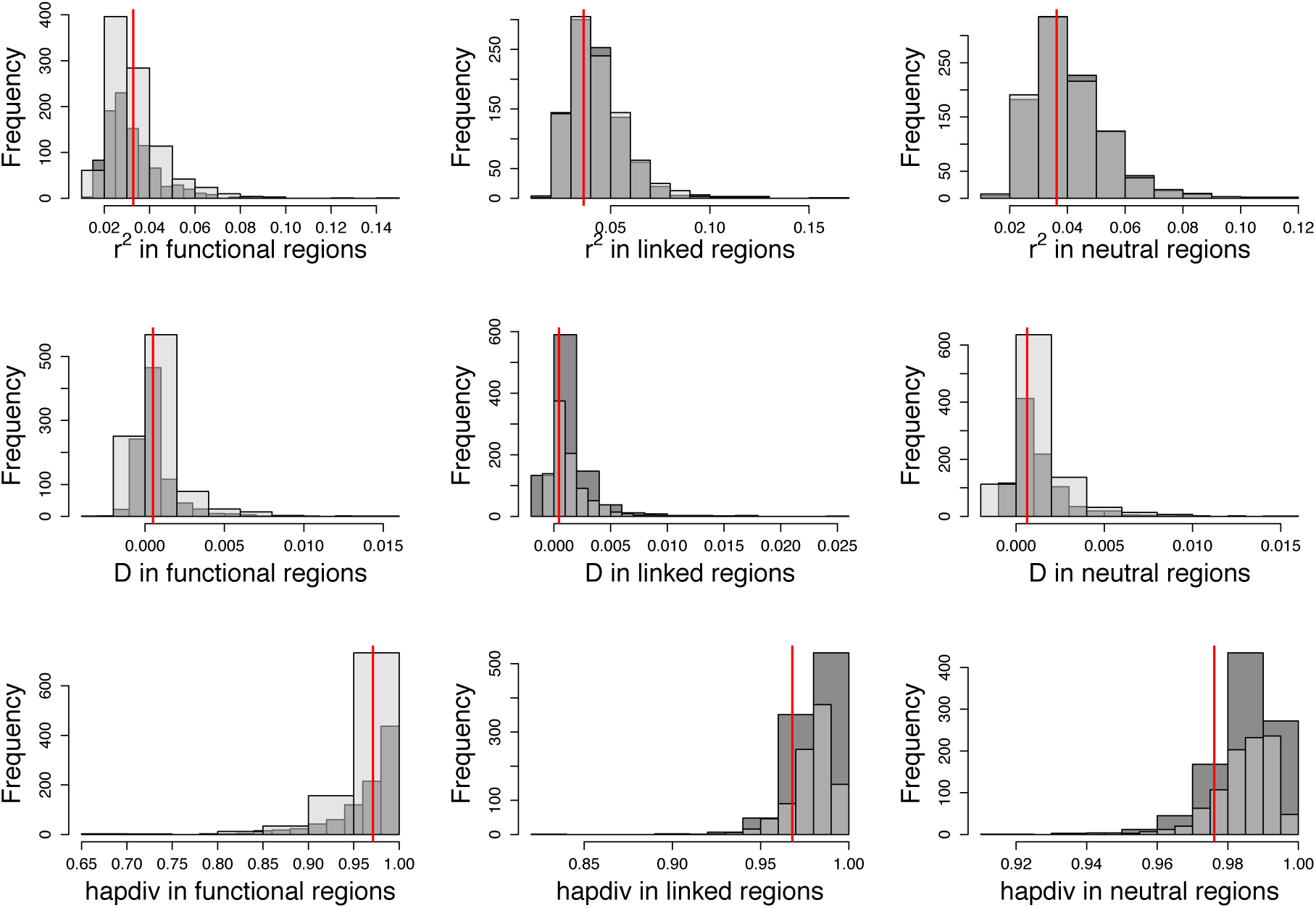
Distribution of summary statistics calculated from 94 exons simulated with 100 replicates each using our inferred model (*i.e*., *f*_0_ = 0.25, *f*_1_=0.49, *f*_2_=0.04, *f*_3_=0.22, *N*_anc_= 1,225,393, *N*_cur_ = 1,357,760). Functional regions were simulated to experience common (5%) and weak positive selection (2*N*_anc_*s* = 10). Red lines indicate the value observed in 76 individuals of *Drosophila melanogaster* from Zambia, after excluding sites with phastCons score **≥** 0.8. Dark grey bars represent no positive selection and light grey bars represent simulations with positive selection in functional regions.

**Figure S22:**
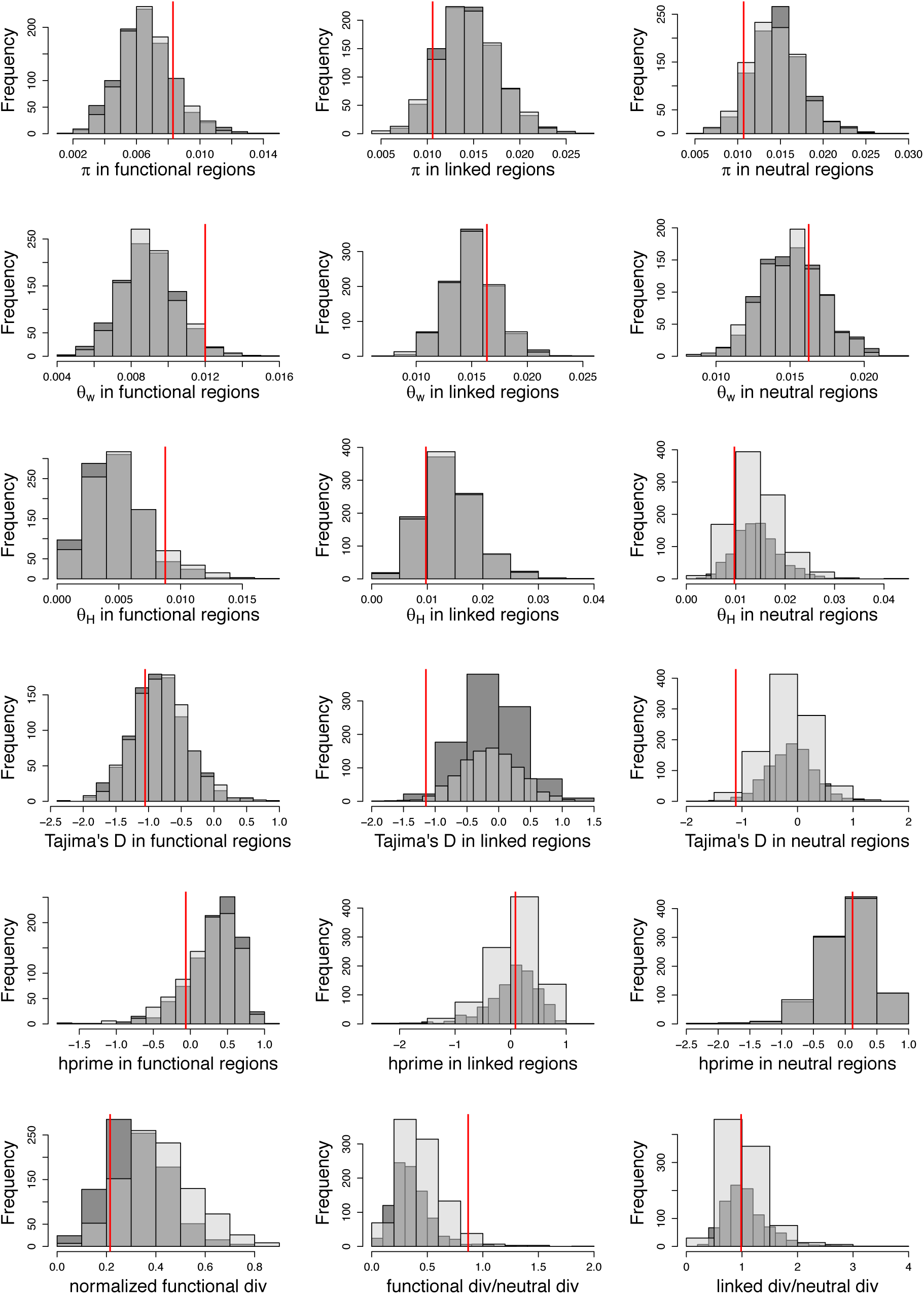
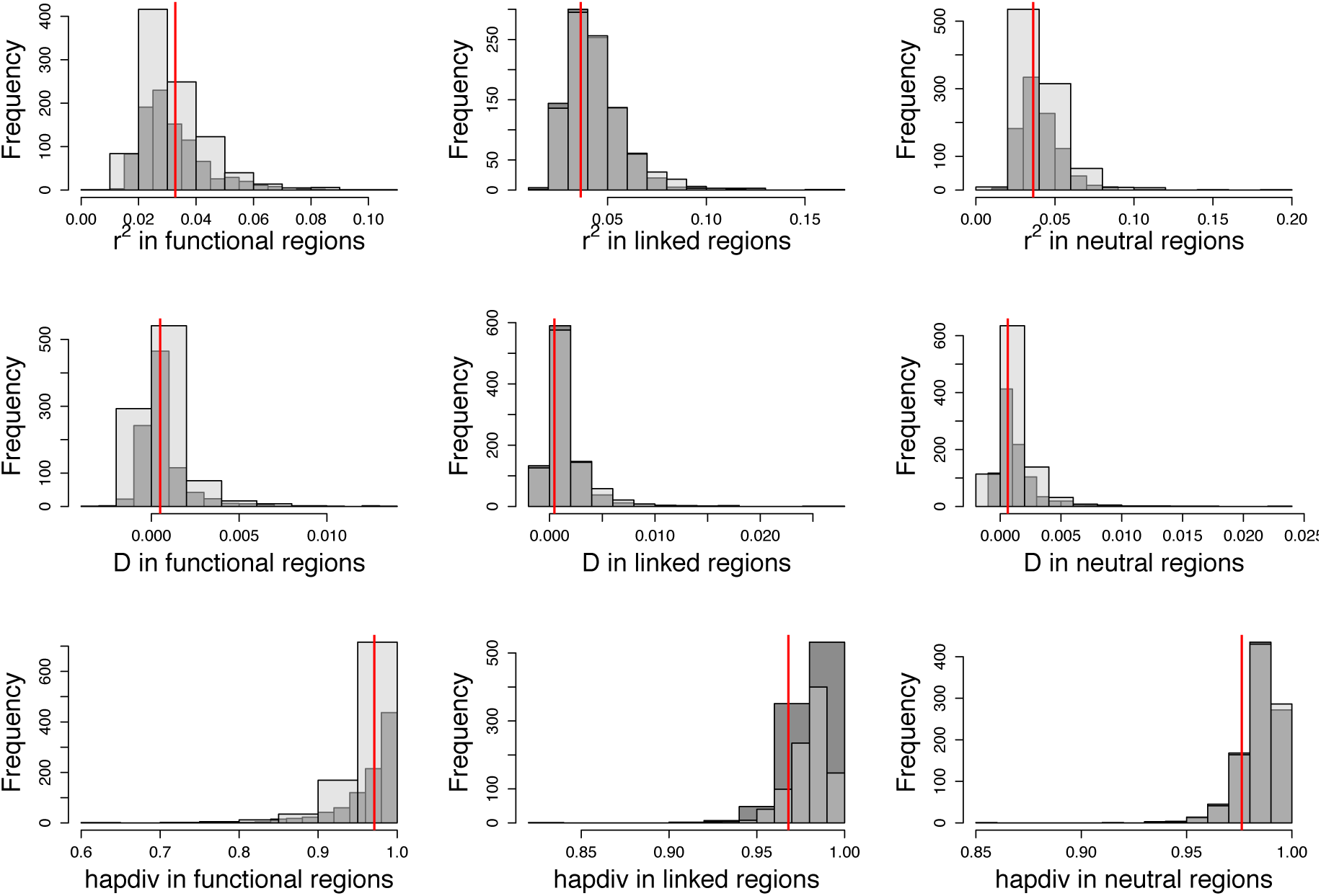
Distribution of summary statistics calculated from 94 exons simulated with 100 replicates each using our inferred model (*i.e*., *f*_0_ = 0.25, *f*_1_=0.49, *f*_2_=0.04, *f*_3_=0.22, *N*_anc_= 1,225,393, *N*_cur_ = 1,357,760). Functional regions were simulated as experiencing rare (1%) and weak positive selection (2*N*_anc_*s* = 10). Red lines indicate the value observed in 76 individuals of *D. melanogaster* from Zambia, after excluding sites with phastCons score **≥** 0.8. Dark grey bars represent no positive selection and light grey bars represent simulations with positive selection in functional regions.

**Figure S23:**
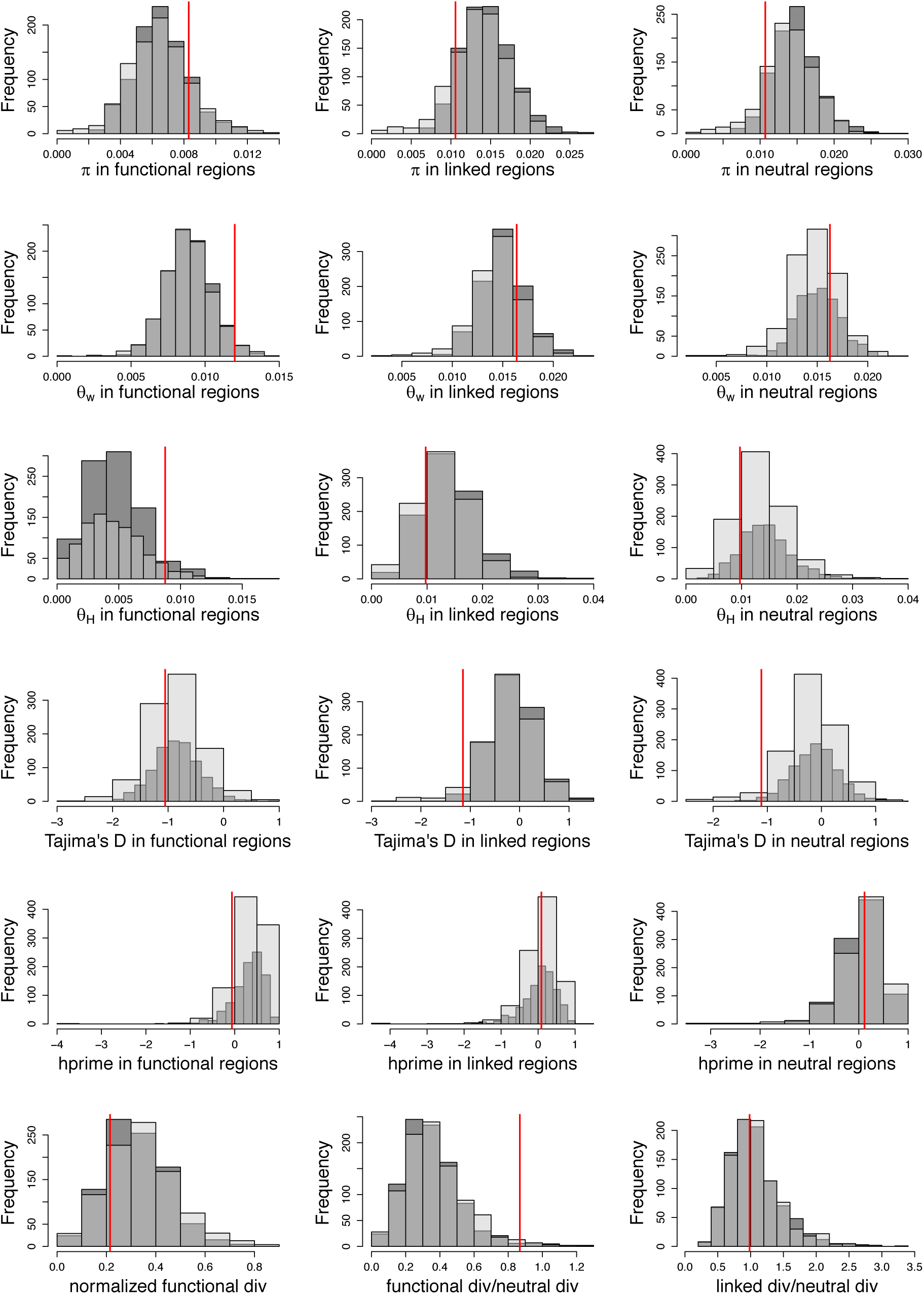
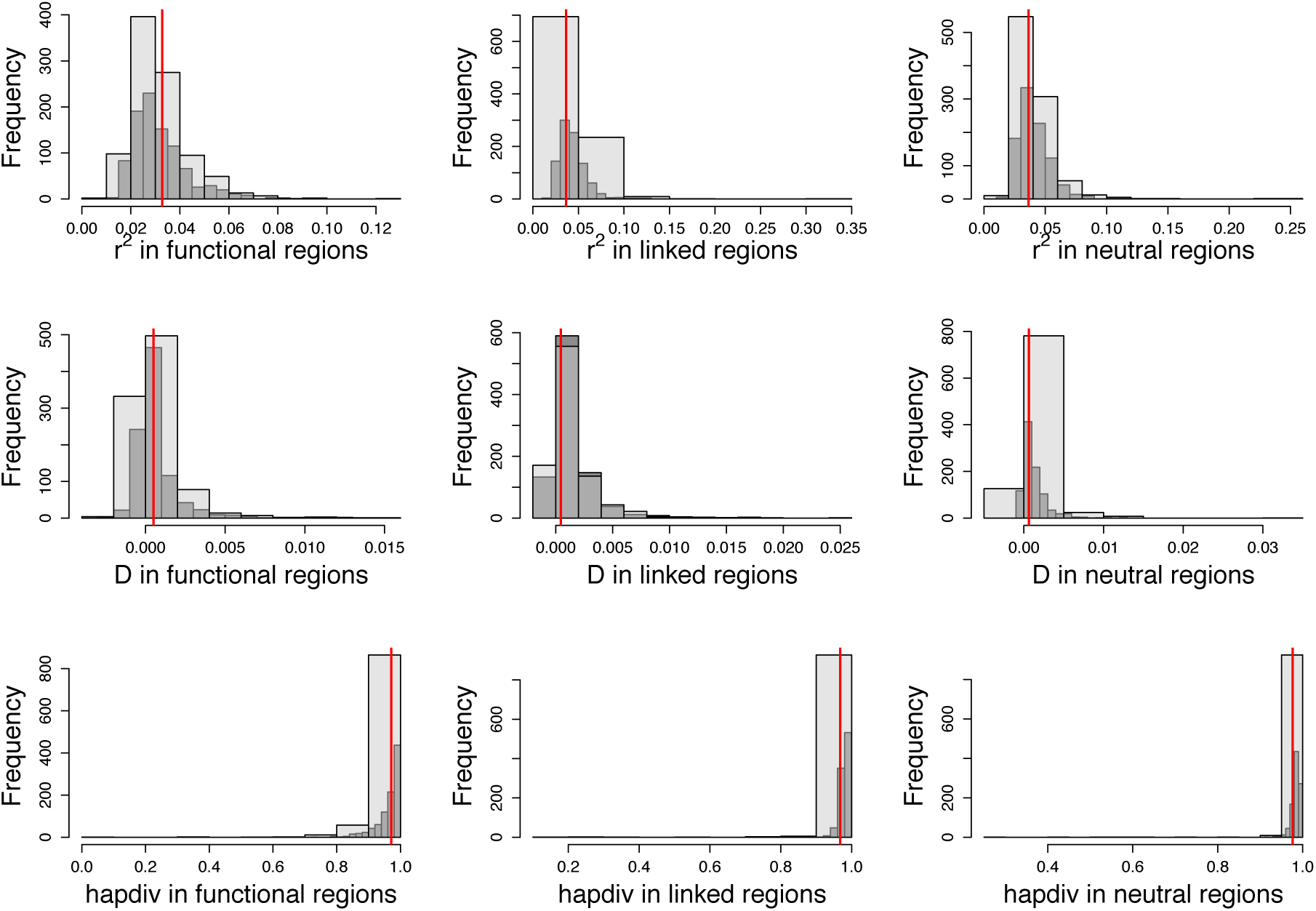
Distribution of summary statistics calculated from 94 exons simulated with 100 replicates each using our inferred model (*i.e*., *f*_0_ = 0.25, *f*_1_=0.49, *f*_2_=0.04, *f*_3_=0.22, *N*_anc_= 1,225,393, *N*_cur_ = 1,357,760). Functional regions were simulated to experience rare (1.28 x 10^-4^ %) and strong positive selection (2*N*_anc_*s* = 10000) as in Lange and Pool (2018). Red lines indicate the value observed in 76 individuals of *D. melanogaster* from Zambia, after excluding sites with phastCons score **≥** 0.8. Dark grey bars represent no positive selection and light grey bars represent simulations with positive selection in functional regions.

**Figure S24:**
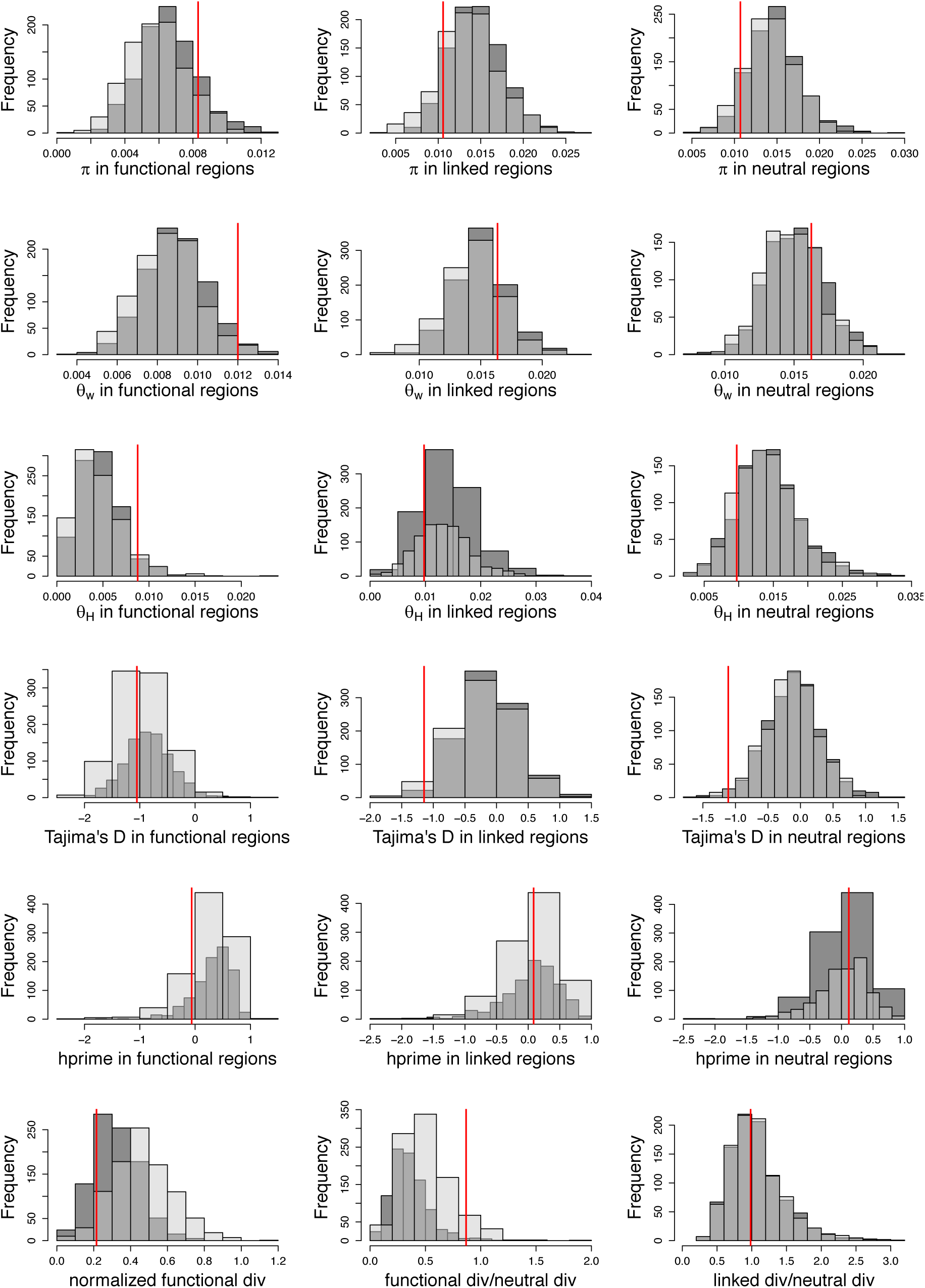
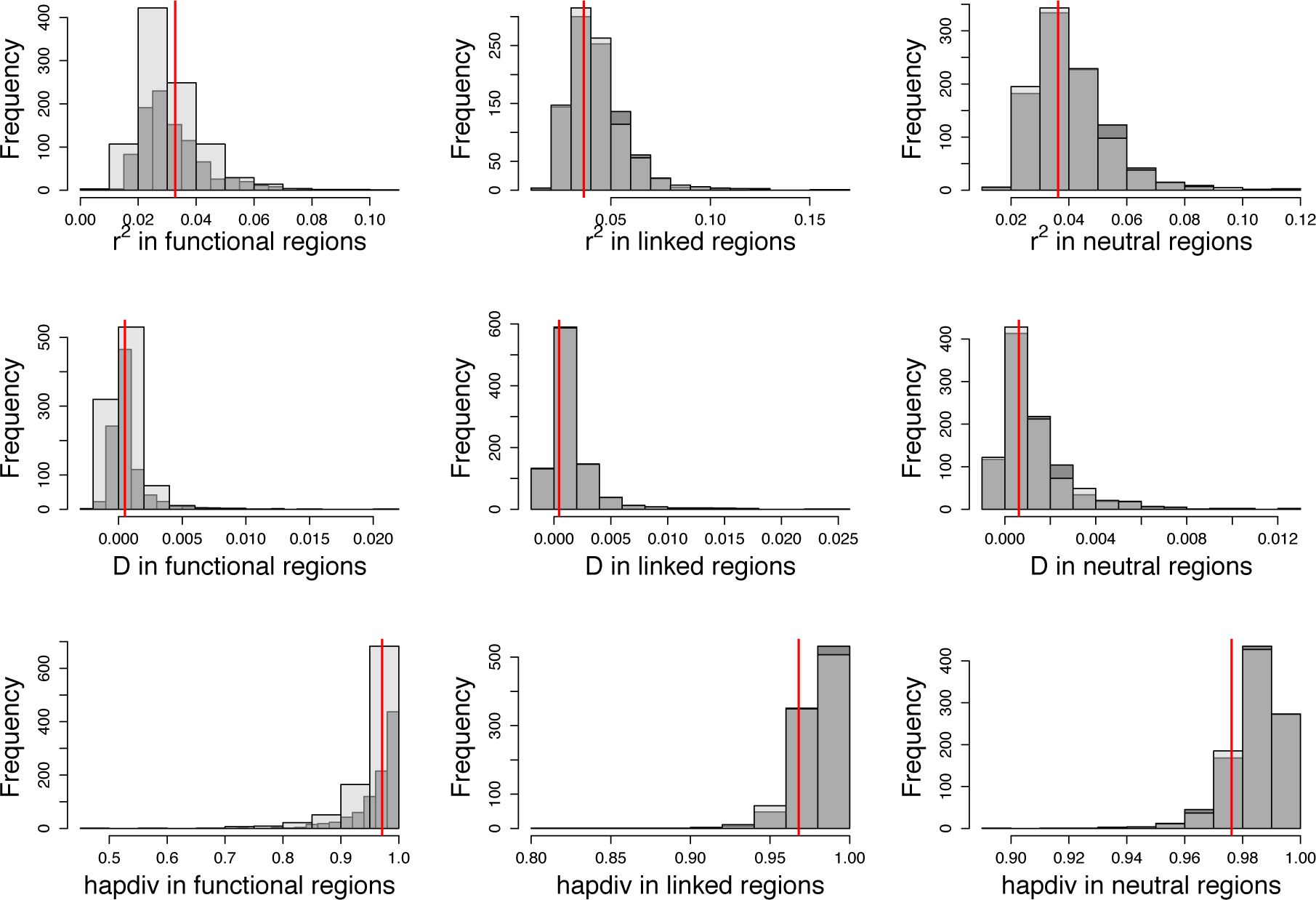
Distribution of summary statistics calculated from 94 exons simulated with 100 replicates each using our inferred model (*i.e f*_0_ = 0.25, *f*_1_=0.49, *f*_2_=0.04, *f*_3_=0.22, *N*_anc_= 1,225,393, *N*_cur_ = 1,357,760). Functional regions were simulated to experience rare (0.2%) and weak positive selection (2*N*_anc_*s* = 60) as in Lange and Pool (2018). Red lines indicate the value observed in 76 individuals of *D. melanogaster* from Zambia, after excluding sites with phastCons score **≥** 0.8. Dark grey bars represent no positive selection and light grey bars represent simulations with positive selection in functional regions.

